# Fasting-mimicking diet counteracts gut microbial dysbiosis in experimental Lynch syndrome

**DOI:** 10.1101/2025.03.14.643045

**Authors:** Lorena Garcia-Castillo, Giulio Ferrero, Olga Blaževitš, Giulia Francescato, Eliass Azra Thaseen, Natalia Erika Cortez, Marc Beltrà, Sonia Tarallo, Barbara Pardini, Paola Costelli, Alessio Naccarati, Valter D. Longo, Fabio Penna

## Abstract

The development of colorectal cancer (CRC) is largely influenced by hereditary factors, with up to one-third of cases linked to genetic predisposition. In parallel, environmental factors such as diet and intestinal microbiota play a significant role. Lynch syndrome (LS), the most common form of hereditary CRC, is due to mutations in DNA mismatch repair genes. Diet interventions such as calorie restriction (CR) can modify the course of the disease, altering nutrient supply and promoting beneficial microbial populations. Fasting-mimicking diets (FMD) are plant-based CR regimens that showed promise in modulating the gut microbiota and suppressing CRC progression in pre-clinical ectopic cancer models.

In this study, Villin-Cre/Msh2-floxed (VCM) mice, modelling LS, were subjected to periodic FMD cycles for 10 months.

Although not impacting on macroscopic tumor development, FMD influenced animal weight in a sexually dimorphic manner. Moreover, shotgun metagenomic sequencing revealed that FMD mitigated the dysbiotic longitudinal changes associated with cancer onset, preserving beneficial species, such as *Lactobacillus johnsonii*, and reducing adverse species, such as *Escherichia coli*.

Metabolic pathway analysis also showed significant differences, with FMD preventing the upregulation of pathways involved in amino acid and nucleotide synthesis, potentially promoting tumor growth.

Overall, the findings suggest that periodic FMD may be adopted as an adjuvant therapy in LS management, counteracting gut microbiota alterations.

## Introduction

Colorectal cancer (CRC) is the third most diagnosed cancer in the world and represents the second leading cause of cancer deaths [1,2]. Currently, it is estimated that 10-35% of CRCs are linked to a hereditary predisposition, with a specific genetic cause identified in 3-5% of these cases [3,4]. The most frequent form of hereditary CRC is known as Lynch syndrome (LS) [5]. LS, also referred to as hereditary nonpolyposis CRC (HNPCC), is characterized by an autosomal dominant inheritance pattern that significantly increases the risk of developing colorectal and endometrial cancers, among others. It is caused by a mutation in one of the DNA mismatch repair (MMR) genes *MLH1, MSH2, MSH6, PMS2* or a deletion in the 3’ region of the *EPCAM* gene. Defective MMR pathway causes microsatellite instability (MSI), which leads to the early onset of colorectal neoplastic lesions [1,3,5]. In addition to hereditary predisposition, environmental factors such as dietary habits, sedentary lifestyle, obesity, alcohol consumption, and intestinal microbiota significantly aid in the development of sporadic CRC [6,7].

Beyond intrinsic enterocyte genetic alterations driving tumorigenesis, several studies have demonstrated a correlation between gut dysbiosis and the risk of onset and progression of sporadic CRC [8–11]. Indeed, some specific bacterial species, including *Fusobacterium nucleatum, Escherichia coli, Bacteroides fragilis*, and *Streptococcus gallolyticus* have been associated with colon tumorigenesis [12]. Mechanistically, one of the most common pathways in CRC development involves the inflammatory and immune responses triggered by the interaction between CRC-promoting bacteria and the host [12]. In addition, microbe-derived metabolites in the intestine can act as a genotoxic agent by promoting inflammation and DNA damage in the colon, altering genome stability [13]. For instance, secondary bile acids, such as deoxycholic acid (DCA), enhanced the development of CRC in a cancer-resistant mouse strain after treatment with azoxymethane [14]. In another study, higher levels of DCA were observed in the serum of men with colorectal adenomas compared to healthy subjects [15]. In addition, the presence in the colon of colibactin-producing *E. coli pks+ (polyketide synthases)* has been shown to accelerate the mutations and the relative contribution of mutational signatures associated with MMR deficiency [16,17].

Dietary habits affect the composition and metabolism of the gut microbiota. In this respect, the availability of undigested food influences the abundance, diversity, function, and balance among gut microbial species [18,19]. Calorie restriction (CR) and fasting may alter the gut environment by changing the nutrient supply and energy sources, which can directly influence the growth of certain microbial taxa and microbiota-derived metabolite production [20]. In mice subjected to nutrient deprivation for 24 hours, the abundance of *Akkermansia, Parabacteroides*, and *Muribaculaceae* increases, while the populations of *Lactobacillus* and *Bifidobacterium* decrease. Interestingly, this modulation is reversed upon refeeding [21]. Given that CR impacts on the microbiota and in parallel interferes with CRC tumorigenesis, the two events may be causally linked; however, information on this area remains limited. So far, preclinical data have shown that increased *Lactobacillus* [22] and *Bifidobacteria* [23] in the gut during CR have key roles in CRC suppression.

The fasting-mimicking diet (FMD), a plant-based, calorie restricted diet, was designed to achieve the advantages of fasting while providing micronutrient nourishment and minimizing the stress of fasting [24]. In a murine model of inflammatory bowel disease (IBD), FMD reduced systemic inflammation and increased *Lactobacillaceae* levels approximately 45-fold compared to normal chow [25]. In a recent study, the increase of *Lactobacillus johnsonii* in the intestine of animals subjected to FMD was essential for the anticancer effect of FMD in an ectopic CRC model [26].

Here, we explore the effects of FMD in the gut microbial population before and after tumor onset in an experimental genetic model of spontaneous intestinal tumorigenesis. Our findings indicate that periodic cycles of FMD, although not impacting tumor appearance, counteract gut microbiota alterations observed during the progression of CRC.

## Methods

### Animal model and experimental design

The study was performed in accordance with the ethical standards of the Helsinki Declaration and all the experimental animal procedures were performed in compliance with the Italian Ministry of Health Guidelines and the Policy on Humane Care and Use of Laboratory Animals (NRC, 2011). The Bioethical Committee of the University of Torino approved the experimental protocol. The animals were maintained on a regular dark-light cycle of 12:12 h with controlled temperature (20-23°C) and humidity (40-60%).

The Villin-Cre/ Msh2loxP/loxP (VCM) mouse model was developed by crossing B6.Cg-Tg (Vil1-cre) 997Gum/J (Villin-Cre) and B6.Cg-Msh2tm2.1Rak/J (Msh2loxP) mice purchased from The Jackson Laboratory (Bar Harbor, CA, USA). The offspring bears a conditional knockout of the *Msh2* gene in the enterocytes, accelerating the spontaneous formation of intestinal adenomas/adenocarcinomas [27]. The presence of each transgene construct was assessed on DNA extracted from the distal tail tip (2 mm) of conscious, non-anesthetized animals [28] using Melt Curve Analysis (RT-PCR) with the following primers: Villin-Cre (forward: 5′-TTCTCCTCTAGGCTCGTCCA-3′ and reverse: 5′-CATGTCCATCAGGTTCTTGC-3′) and Msh2loxP (wild-type: 5′-GATGATGTGTGAAGCCTGCAT-3′, mutant: 5′-CCTCTTGAGGGGAATTGAAGT-3′ and common: 5′-AGGTTAAAAACCAGAGCCTCAACT-3′).

After reaching three months of age, male and female VCM mice were randomized and divided into four groups; 1) VCM fed with standard diet (SD; from hereby called VCM+SD) males and 2) VCM+SD females, 3) VCM fed with FMD (from hereby called VCM+FMD) males and 4) VCM+FMD females. Endpoint was set at ∼13 months of age, a time point when, in general, more than 80% of VCM mice show macroscopic tumors [29]. Hence, animals underwent ∼20 cycles of FMD over a ten-month experimental period. Not all the animals were included simultaneously in the experimental set-up, rather they were included as they reached three months of age from our in-house breeding colony. Body weight and food intake were recorded regularly. Blood glucose and ketones were assessed when the animals reached twelve months of age (Figure 1). Fecal samples were collected at two time points (always between 3-5 pm): T1 (corresponding to 6 months of age) and T2 (at 12 months of age). Mice were examined daily for signs of distress and eventually euthanized when they reached the humane endpoint.

**Figure 1.**
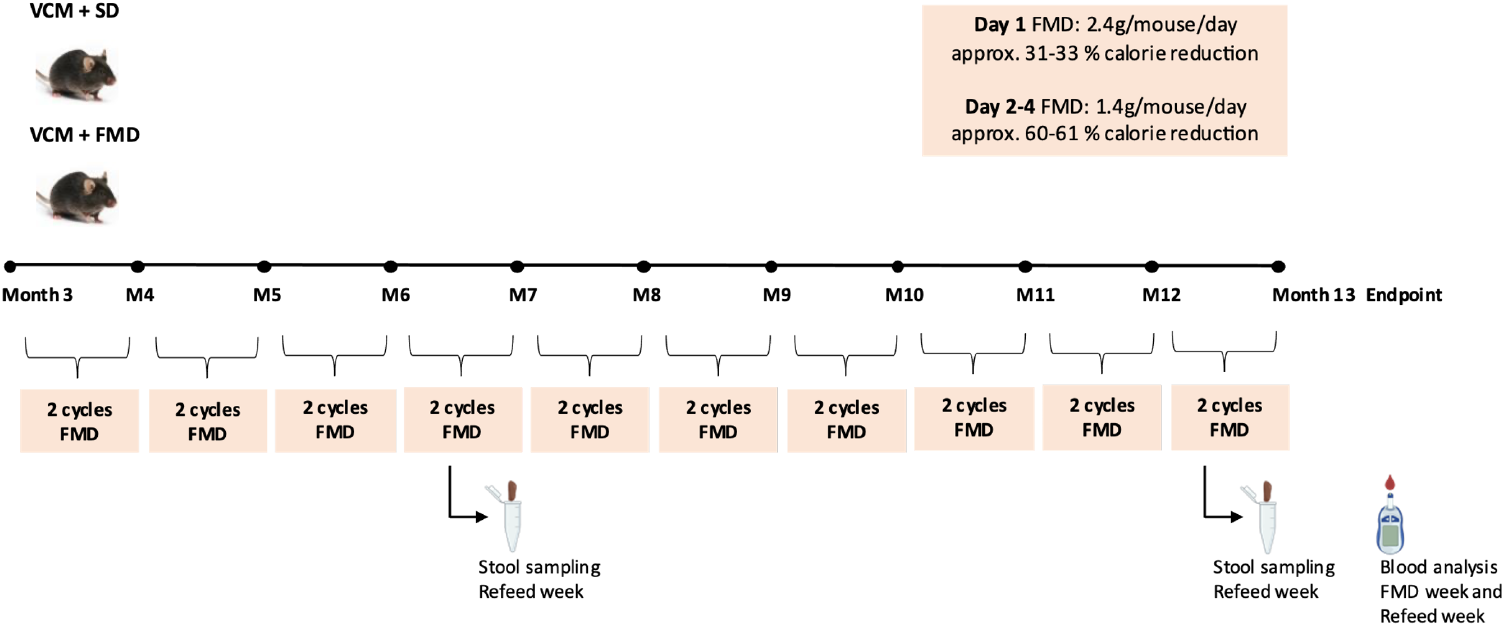
Schematic representation of the experimental design. The cartoon depicts the study experimental plan, including the Fast-Mimicking Diet (FMD) protocol (twice every month until the endpoint), fecal collection, blood ketone and glucose analysis, and sacrifice time point. The experimental groups consisted of 1) Male VCM mice fed with standard diet (SD) (n=12); 2) Female VCM mice fed with SD (n=12); 3) Male VCM mice fed with the FMD (n=14); 4) Female VCM mice fed with the FMD (n=12).

### FMD diet

In humans, the FMD is designed to provide sufficient nourishment during periods of low caloric intake consisting of a 5-day regimen, as described in Brandhorst et al., 2015 [30]. In our study, the FMD regimen was adapted for just 4 days of diet as the animals achieved a ∼10-20% body weight loss by the fourth day (Figure 1). On day one, animals received 2.4g of FMD/mouse/day to adapt to a period of low caloric intake. On days 2-4 animals received 1.4g FMD/mouse/day. Hence, mice in the FMD group experienced a ∼31% and ∼61% calorie reduction on day one and days 2-4, respectively. Before supplying the FMD, the animals were transferred into fresh cages to avoid coprophagy and/or feeding with residual chow diet. Mice consumed all the supplied food of their daily FMD regimen and showed no signs of food aversion. After the end of the FMD cycle, animals were supplied with a standard laboratory chow (SD, Special Diets Services; VRFI) *ad libitum* before starting another FMD cycle.

### Assessment of blood ketones and glucose levels

To assess the proper intake of FMD regimen, blood glucose and ketone levels were evaluated in a small amount of peripheral blood, obtained by the removal of the distal tail tip (2 mm) of conscious, non-anesthetized animals [28]. Tail tip sampling is a minimally invasive procedure and was carried out by snipping less than 2 mm of tissue from the tail tip with sharp scissors. Blood was obtained by direct flow or by gently massaging the tail and then collecting the blood directly on a glucose or ketone test strip (Nova Pro device).

### Free fatty acid assay

Free fatty acid (FFA) concentration in plasma was quantified using a commercial kit (MAK044, Sigma), according to the manufacturer’s instructions. All samples, blanks, and standards were prepared in duplicate.

### DNA extraction and shotgun metagenomic library preparation

Stool DNA extraction was performed with the DNeasy PowerSoil Pro Kit (Qiagen) according to the manufacturer’s instructions. Libraries were prepared starting from 24ng of fecal DNA using the Illumina® DNA Prep, (M) Tagmentation kit (Illumina) as described in [31,32]. The procedure followed the manufacturer’s protocol, except for two in-house modifications as performed by Thomas et al., 2019 [32]: 1) a final clean-up of the pool to be sequenced was performed with 0.6X AMPure XP beads (Beckman-Coulter), and 2) the final library pool was resuspended after the clean-up with 1/3 of the initial pool volume. These adjustments allowed to retrieve higher quality reads with a lower adapter content. The sequencing on the NovaSeq 6000 System was performed at the Italian Institute for Genomic Medicine (IIGM) sequencing facility.

### Shotgun metagenomic data analysis

Sequenced metagenomes were pre-processed using the pipeline available at https://github.com/SegataLab/preprocessing: for i) removal of low-quality reads (quality Q<20), too short fragments (length <75 bp), and reads with 2 or more ambiguous nucleotides; ii) host and contaminant DNA removal using Bowtie 2.77 (--sensitive-local) for filtering the phiX174 Illumina spike-in and mice-associated reads (mm10 assembly); iii) creation of paired forward and reverse and unpaired reads output files.

MetaPhlAn 4 (version 4.1, database vJun23, with the “--statq 0.1”) [33] and HUMAnN 3.9 [34] profiling tools were applied to define the microbial taxonomic and functional profiles, respectively. Microbial diversity analysis was performed using the functions implemented in the vegan v2.6.4 R package, while differential abundance analysis was performed with SIAMCAT v2.4 [35]. The analysis was performed on Species-levels Genomic Bins (SGBs) by removing the lowly abundant SGBs (nearZeroVar function of caret R package) and those without a defined genus annotation.

### Statistical analyses

R v4.3.1 was used for statistical analyses and graphical representation of the results. Non-parametric tests were performed with the Wilcoxon Rank-Sum method. Multidimensional scaling (MDS) statistical analysis was performed using the Bray-Curtis dissimilarity index with the Vegan R package. Paired differential analyses were performed among time-points while comparisons between SD and FMD animals were adjusted for sex. A test was considered significant if associated with a p-value lower than 0.05.

## Results

In this longitudinal study, VCM mice were subjected to periodic cycles of FMD for ten months as described in the ‘methods’ section (experimental protocol depicted in Figure 1). VCM mice develop heterogeneous and non-synchronized tumors, thus not all mice reached the established 13-month endpoint or developed macroscopic tumors at the endpoint. However, the sample size was not powered to draw conclusive evidence on the effect of FMD on tumor incidence in VCM mice. In the VCM+SD group, either males or females (n= 12 each), only 8 males and 6 females showed macroscopic tumors at the endpoint or died prematurely, likely due to tumor presence. In the group subjected to FMD (comprising 14 males and 12 females), a total of 12 males and 8 females exhibited the presence of tumors at endpoint or died prematurely (Supplementary Table 1).

### FMD impacted on systemic metabolism of VCM mice

FMD was shown to stimulate molecular hallmarks of fasting, like low blood glucose levels and increased ketone bodies in circulation. Aiming to shed light on the metabolic shift caused by FMD, blood glucose and β-Hydroxybutyrate (β-HB) levels were assessed when mice reached twelve months of age (Figure 2) either the last day of the FMD week or one week after the refeeding. During the FMD cycle, both males (Figure 2A) and females (Figure 2B) presented a significant decrease in blood glucose levels that was restored during the refeeding.

**Figure 2.**
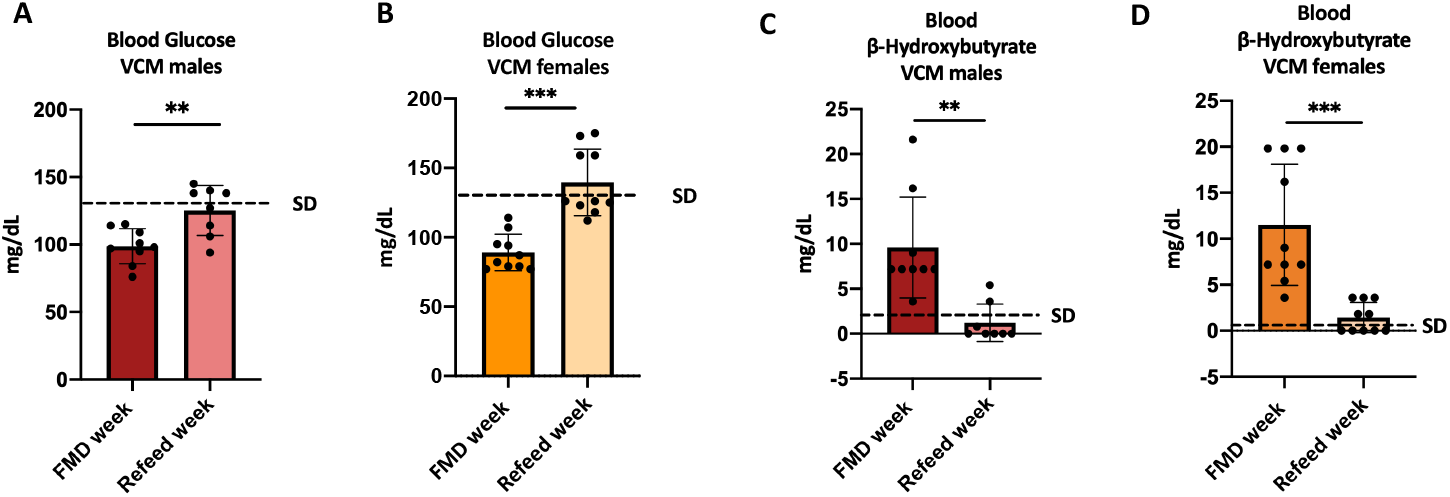
Blood glucose and β-HB levels in VCM mice. **A-B)** Blood glucose (means ± SD) levels expressed in mg/dL on the last day of the FMD week or one week after refeeding in males (**A**) or females (**B**). **C-D)** Blood β-HB (means ± SD) levels represented as mg\dL on the last day of the FMD week or one week after refeeding in males (**C**) or females (**D**). VCM+FMD males (n=9) and females (n=10). Student’s t-test: *p < 0.05; **p < 0.01; ***p<0.001.

In physiological conditions, the concentration of β-HB is very low (≤ 5.4 mg/dL) while during nutritional ketosis this concentration increases over 9 mg/dL [36,37]. β-HB levels were significantly different during the FMD cycles in both sexes (males, p=0.0012; females, p=0.0002) although there was high variability among samples. Circulating levels of β-HB increased during the FMD week reaching “mild” ketosis in males (9.54±5.58mg/dL; Figure 2C) and females (11.52±6.48mg/dL; Figure 2D). After refeeding, the blood glucose and β-HB levels in the VCM+FMD group were comparable to the VCM+SD group in both males (glucose: 133.56±17.55mg/dL, ketones: 2.16±2.52mg/dL) and females (glucose: 132.89±20.32mg/dL, ketones: 0.36±0.72mg/dL). Overall, these results suggest that FMD impacted the metabolic state and stimulated a fasting environment in VCM mice.

### Effect of FMD on body weight and plasma FFA levels in VCM mice

Fasting and CR have been popular strategies to treat obesity and protect against metabolic syndrome [38,39]. Indeed, alternate day fasting in rodent models of obesity has been shown to decrease total plasma cholesterol and triglyceride concentration [39]. The monitoring of VCM animal body weight shows significant differences between male and female animals. Male FMD-fed mice were characterized by a lower weight than male VCM+SD animals (Figure 3A). Conversely, the weight of female mice increased on average in the FMD group (Figure 3B). Notably, during FMD week both males and females had reduced energy intake which returned to the same levels as controls after several days of refeeding (Figure S1).

**Figure 3.**
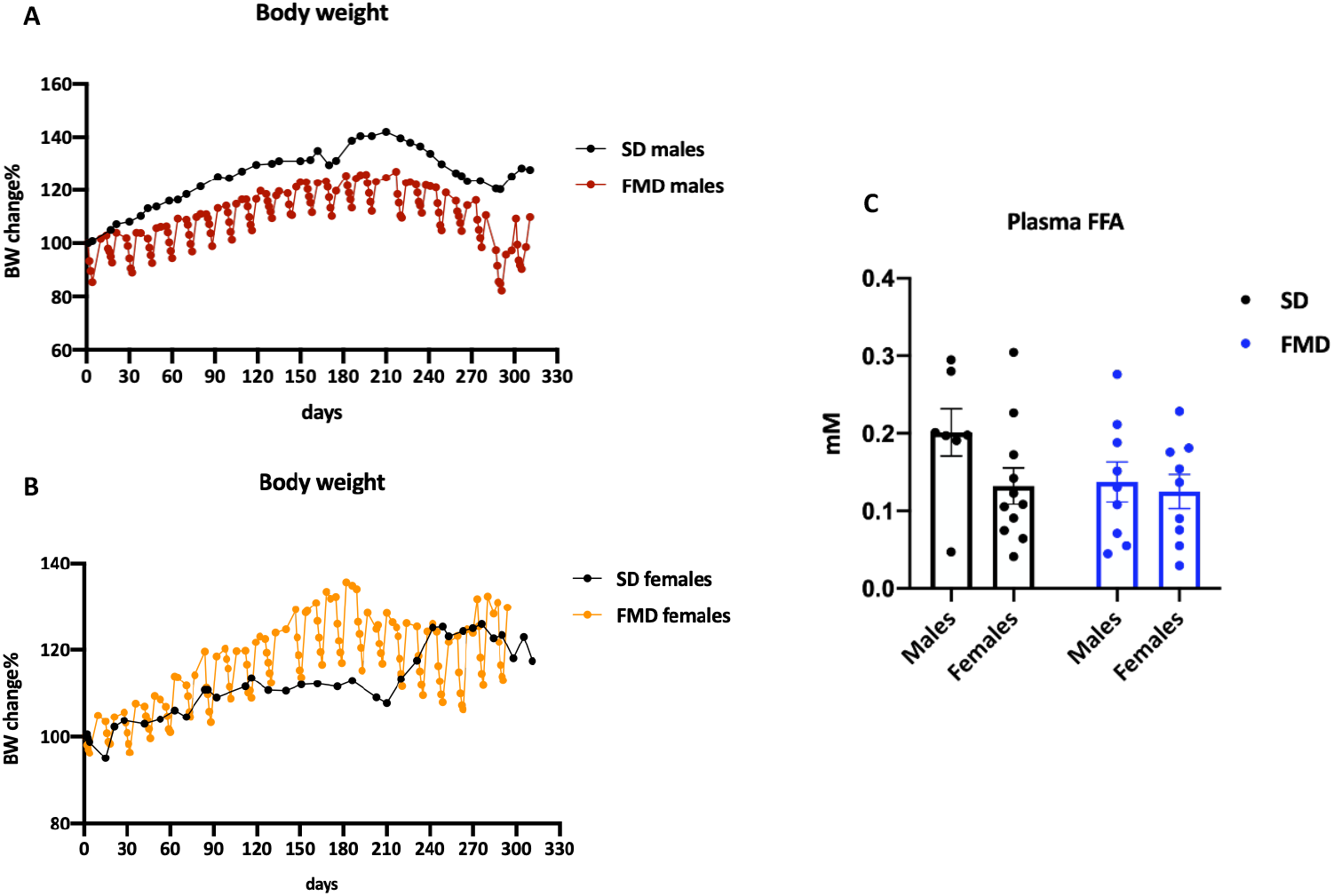
Body weight and plasma free fatty acid (FFA) levels in VCM. **A-B)** Body weight change in VCM+SD and VCM+FMD males (**A**) and females (**B)**. Body weight is expressed as a percentage of the initial body weight of the average weight of the experimental animals. Only mice showing macroscopic tumor masses at necropsy or died prematurely are represented. VCM mice fed with SD males (n=8) and females (n=6). VCM mice fed with FMD males (n=12) and females (n=7). **C**) Histograms reporting plasma FFA levels in VCM+SD and VCM+FMD mice, stratified for sex. Plasma FFA levels are expressed in mM, measured at the endpoint during the refeeding. VCM+SD males (n=7) and females (n=11); VCM+FMD males (n=9) and females (n=9).

To investigate the effect of FMD on circulating FFA, the plasma of VCM mice was analyzed at the endpoint, during the refeeding week. FMD fed animals showed similar plasma FFA to those fed a SD in both males and females (Figure 3C). However, when comparing only males and females in the SD group, males had a significant increase (p=0.0461) of FFA in comparison to females.

### Modulation of gut microbiota and microbial metabolic pathways by FMD

Since possible beneficial effects on the gut microbiota by dietary restrictions [20], including FMD [25,26], have been previously reported, fecal samples were collected at two timepoints (Figure 1) in order to evaluate gut microbiota composition after the FMD. Shotgun metagenomic sequencing was performed on fecal samples collected for both groups at the age of six months, before the frankly appearance of tumors (T1), and at twelve months, the time in which tumors are likely present in the majority of the mice (T2) (for the VCM+FMD fed group one week after refeeding).

No significant changes in microbial richness (number of detected species) were observed neither between T1 and T2, nor between VCM+SD and VCM+FMD samples (Figure 4A). Similarly, no significant differences among groups were observed with the other two alpha diversity metrics, the Shannon Index and Inverse Simpson index (Figure S2A-C and Supplementary Table 2A). On the other hand, the VCM+SD group at T2 showed a higher beta diversity (measuring the differences in microbial composition between samples) than T1 samples of the same animals but also than VCM+FMD in all time points and the time variable was significantly associated with the beta diversity (PERMANOVA p=0.001; Figure 4B and Supplementary Table 2B).

**Figure 4.**
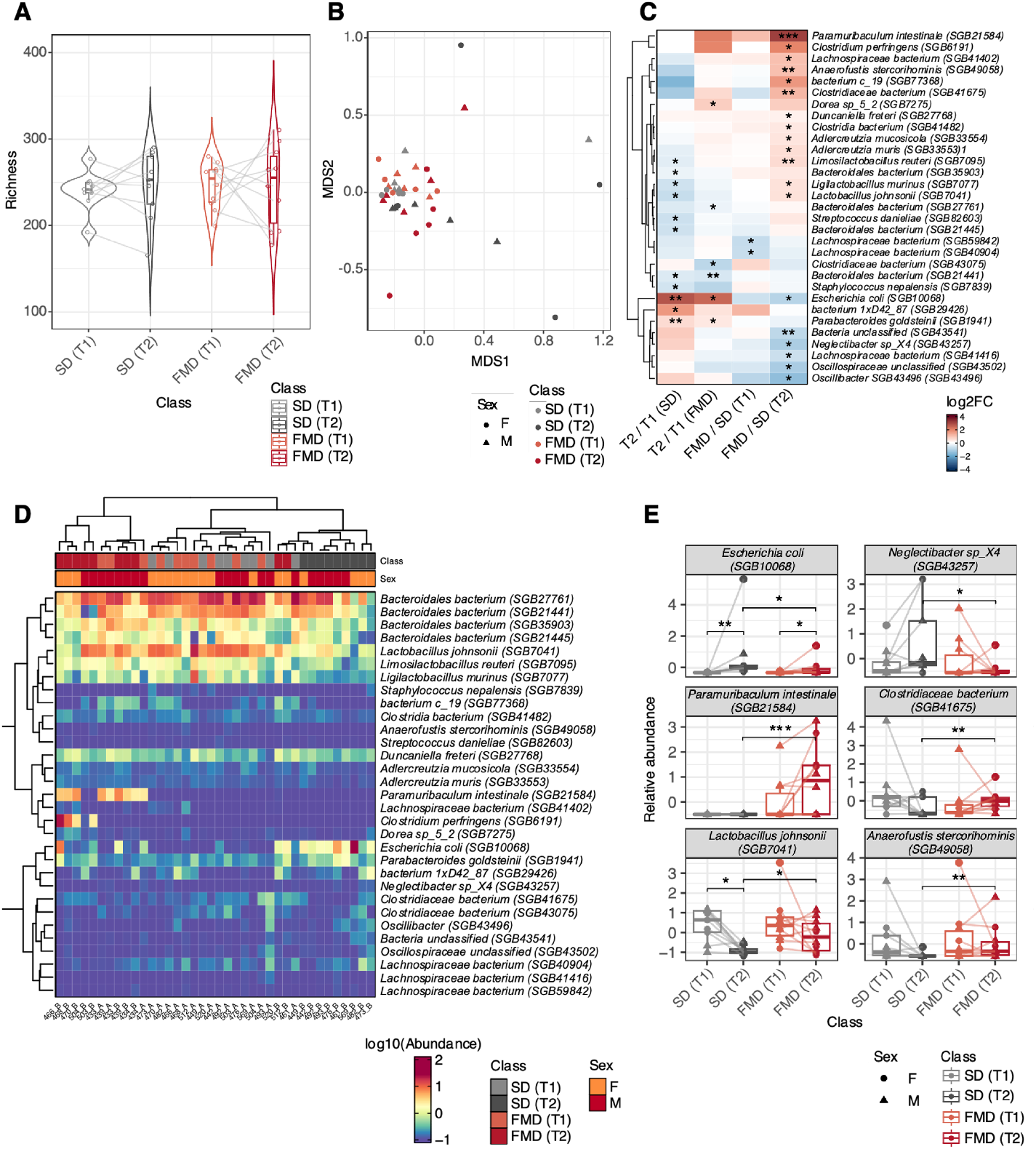
Analysis of the fecal microbial composition among samples of VCM+SD and VCM+FMD mice collected at T1 and T2. **A)** Violin plots of the microbial richness computed for each study group at the two time points. P-value by Wilcoxon Rank-Sum test. **B)** Multidimensional scaling (MDS) plots representing the beta diversity samples from SD and FMD mice at T1 and T2. VCM+SD males and females (n=9) and VCM+FMD males and females (n=10). **C)** Heat map reporting the species-level genome bins (SGBs) with significantly different abundances among the study groups. The color-code represents the log2FC of SGB abundances as computed in the comparisons reported in columns. Significance of the differences by SIAMCAT test: *p<0.05; **p<0.01; ***p<0.001. Paired analysis was performed for the T2 vs T1 comparisons. **D)** Heat map with relative abundance levels (reported in log10) of the differentially abundant SGBs. Heat map columns are annotated with the study groups and sex information. VCM+ SD (n=9) and VCM+FMD (n=10). **E)** Box plots reporting the relative abundances of six bacterial species in the VCM+FMD and VCM+SD samples. Significance of the differences by Wilcoxon Rank-Sum test: *p<0.05; **p<0.01.

Differential analysis of the relative abundances by microbial phyla showed an increase of Firmicutes and Proteobacteria at T2 of both groups (Figure S2D-E). Differentially abundant SGBs, time or diet-dependent, are reported in Figure 4C and Supplementary Table 2C. Observing the VCM+SD group over time, some species, such as *E. coli (SGB10068)* and *Parabacteroides goldsteinii (SGB1941)* increased (paired Wilcoxon test p<0.05), while others such as *Bacteroidales bacterium (SGB21441)* and *Lactobacillus johnsonii (SGB7041)* decreased at T2. Similarly, in the VCM+FMD group, the abundance of the same species increased when animals approached the endpoint.

The clustering analysis showed a clear separation between the samples at T2, highlighting SGBs such as *E. coli (SGB10068)* in VCM+SD animals or *Paramuribaculum intestinale (SGB21584)* in VCM+FMD with specifically higher abundance (Figure 4D). Moreover, significant differences (p<0.05) in the abundance of 19 species were observed between SD and FMD at T2. Details of some of these taxa are reported in Figure 4E. For instance, the *E. coli (SGB10068)* abundance increased at T2 in both VCM+SD and VCM+FMD; however, this increase was significantly smaller in animals subjected to FMD cycles. Similarly, the increase of *Neglectibacter sp_X4 (SGB43257)* observed in SD-fed mice at T2 was not found in animals subjected to the FMD regimen. Conversely, *P. intestinale (SGB21584)* and *Clostridiaceae bacterium (SGB41675)* abundance increased in VCM+FMD at T2 compared with the VCM+SD. Furthermore, the FMD prevented the general decrease in the abundance of *L. johnsonii (SGB7041) and Anaerofustis stercorihominis (SGB49058)*, as observed at month 12.

Given the known relevance of *E. coli* colonization in CRC development and to gain better insight into the possible influence of animal sex, we evaluated the relative abundance of these bacteria. Our analyses revealed an increased abundance of *E. coli (SGB10068)* in VCM+SD animals at T2 compared to VCM+FMD mice, particularly in males (p=0.016) (Figure 5A). Since among the *E. coli* strains, those encoding for the colibactin (*pks*^+^) are involved in CRC development [40], we performed the quantification of this gene with HUMAnN v3.9, observing a lack of genes belonging to the *pks* island in all tested samples.

**Figure 5.**
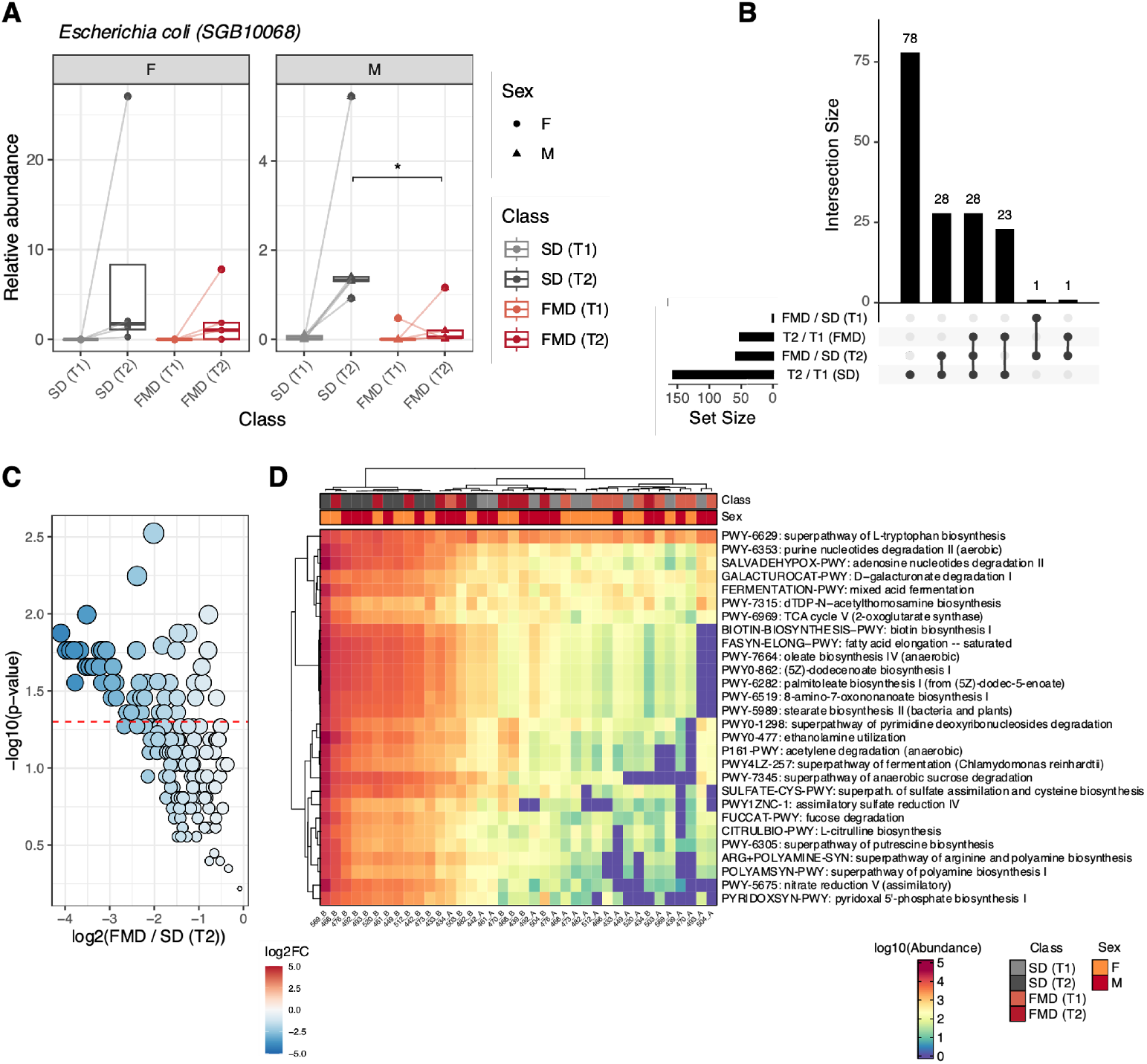
**A)** Box plots reporting the relative abundance of *E. coli (SGB10068)* in samples from animals in the VCM+FMD or VCM+SD groups stratified by sex. Significance of the differences by Wilcoxon Rank-Sum test: *p< 0.05. Paired analysis was performed for the T2 vs T1 comparisons. **B)** Upset plot of the number of microbial pathways associated with significantly different levels between study time points and VCM+SD and VCM+FMD animals. **C)** Volcano plot of the change of microbial pathway levels between T2 time points of VCM+SD and VCM+FMD animals considering 157 pathways observed as significantly modulated between T2 and T1 time points of VCM+SD mice. P-value by Wilcoxon Rank-Sum test. The red dashed lines represent the p-value threshold of 0.05. **D)** Heatmap of the levels of microbial pathways, showing the rescue determined by FMD at T2.

The analysis of microbial metabolic pathways showed significant differences among the study groups (Supplementary Table 2D). Specifically, among the 234 microbial pathways analyzed, respectively 157 and 52, were associated with a significant (paired Wilcoxon test p<0.05) modulation in VCM+SD and VCM+FMD, with the levels of all of them increasing at T2 (Figure 5B). Notably, the levels of all 157 pathways modulated in VCM+SD, including the 52 pathways observed in VCM+FMD, decreased at T2 in VCM+FMD animals (Figure 5C). Among them, 29 were associated with levels significantly lower at T2 of VCM+FMD with respect to T2 of VCM+SD mice (Figure 5C-D). The pathways most significantly decreased in VCM+FMD were *PWY-6549: L-glutamine biosynthesis III, P108-PWY: pyruvate fermentation to propanoate I, PWY-6507: 4-deoxy-L-threo-hex-4-enopyranuronate degradation, VALSYN-PWY: L-valine biosynthesis, PWY-6122: 5-aminoimidazole ribonucleotide biosynthesis II, PWY-6277: superpathway of 5-aminoimidazole ribonucleotide biosynthesis, PWY-7198: pyrimidine deoxyribonucleotides de novo biosynthesis IV*.

## Discussion

The FMD is an emerging nutritional regimen focused on implementing specific nutrient combinations and complex food compositions to improve healthspan and lifespan and optimize therapies [41]. Most studies investigating FMD in animal models of cancer have focused on its potential to improve the efficacy of antineoplastic treatments by modulating cancer and host metabolism or enhancing antitumor responses [42–45]. The current study, adopting a mouse model of spontaneous cancer with late and low pace tumor onset, aimed at understanding the impact of long term FMD on host health and microbiota.

FMD did not prevent tumor development, suggesting that, at least in this model with constitutive *Msh2* deletion, FMD alone is insufficient to block the tumorigenesis. In accordance with the literature [30,46], the administration of FMD led to metabolic changes also in VCM animals, with a marked decrease of blood glucose and an increase of β-HB levels. Notably, our study has provided interesting data regarding the sex-based differences in body weight change upon FMD, with males benefiting significantly more from FMD in limiting body weight gain. Several biological factors may explain the dimorphic effect, likely impinging on whole body metabolism, such as hormone levels and differences in adipose tissue distribution and function [47]. Females exhibit higher thermogenic activity than males, while males generally have higher resting energy expenditure [48]. This difference in resting energy may explain why males tend to lose more body weight during FMD-dependent CR. Finally, CR may help prevent cardiometabolic risk factors [49,50]; indeed, elevated plasma FFA levels were observed exclusively in SD-fed male mice, likely as a consequence of increased adipose tissue.

Gut microbiota undergoes significant changes in the course of cancer and intestinal inflammatory diseases [25,51]. More specifically, increased Proteobacteria and Firmicutes/Bacteroides ratio together with a decrease in *Lactobacilli* [52] and *Muribaculaceae* [53] have been observed in humans and mice with CRC. In this research, we observed that variations in beta diversity may suggest a possible connection between tumor progression and microbial diversity in SD-fed mice. FMD previously showed the ability to reduce intestinal inflammation and promote beneficial gut microbiota colonization in an experimental model of inflammatory bowel disease [25]. In the present study, we showed that a long-term intervention with FMD may prevent the microbiota shift observed during CRC progression, increasing the abundance of species belonging to the probiotic bacterial families *P. intestinale* (a species of *Muribaculaceae), P. goldsteinii and L. johnsonii*, crucial for intestinal health [54,55]. *Lactobacillus* and *Bifidobacterium* strains have been implicated, among others, in the suppression of NF-κB activation and IL-6 secretion, along with the production of short-chain fatty acids (SCFAs), exerting a protective action on the intestinal barrier against inflammatory pathologies [56]. In addition, as demonstrated by Luo et al. (2024) [26], FMD promotes the expansion of *L. johnsonii*, which plays a role in modulating the immune microenvironment in mice subcutaneously implanted with MC38 cells, enhancing CD45+ and CD8+ T cell populations. Bacterial species such as *E. coli* have been identified as CRC-promoting bacteria [12,16,17]. Chronic exposure of healthy human intestinal organoids to genotoxic *pks*^*+*^ *E. coli* promoted mutagenesis by inducing specific single base substitution (SBS-*pks*) and small insertions and deletions (ID-*pks*) mutational signatures [57]. Although *pks+ E. coli* has been found in both normal and cancerous intestinal tissues, CRC patients have a higher abundance compared to healthy individuals [16,17]. In our data, the abundance of *E. coli* was increased in the late sampling (T2), especially in the males of the SD group, but no genes related to the *pks* island have been detected. Laboratory mice are typically bred and raised in isolated, germ-free environments that do not naturally harbor specific bacterial strains like *pks*+ *E. coli*. To study the role of these bacteria in cancer [58] and other diseases [59], animals are often deliberately colonized with *pks*+ *E. coli*. In the current study, it remains unclear whether the increase in *E. coli* is a result of changes caused by the tumor microenvironment or if it is an earlier event, potentially acting as a promoter of CRC development. Further research is needed to determine the underlying cause.

The differences in microbial species observed at T2 resulted in the upregulation of microbial metabolic pathways in the SD group, likely resulting in dysbiotic microbiota-derived metabolites. This would suggest that unrestricted feeding in the SD group may have promoted more active microbial metabolism over time compared to FMD, negatively impacting on intestinal health. In contrast, FMD may have selected for a more specialized microbial population that is resilient to reduced gut nutrient availability, which aligns with the increased beta diversity observed in the SD group as it approached the endpoint. Among the most significantly modulated metabolic pathways are those involved in amino acid and nucleotide synthesis, which potentially contribute to the production of alternative energy sources and biosynthetic molecules essential for tumor growth in the SD group. Conversely, the increased metabolic pathway for producing SCFAs (*pyruvate fermentation to propanoate I*) observed in SD group, may counteract CRC proliferation [60], as observed for propionate production in the gut which has been shown to induce CRC cells apoptosis in 3D spheroid culture models [61]. Despite these findings, the activity of these metabolic pathways cannot be determined, requiring further validation through metabolomics.

Overall, our findings indicate that FMD differentially impacts the body weight of animals in a sex-dimorphic manner, highlighting the importance of considering the sex even in the context of nutritional regimen response as it is already emerging for humans [62]. Consequently, it is crucial to carefully consider and address sex-based differences in nutritional studies, as these factors may confound the results and limit their generalizability. Furthermore, this study suggests that chronic intervention with FMD may slow down the metabolic activity of the intestinal microbial population and prevent the microbiota modulations observed during the development of CRC. However, establishing a direct cause between this modulation and specific aspects of the development or progression of CRC remains challenging, especially in light of the lack of a macroscopic impact on the tumor.

In conclusion, while there is evidence suggesting that FMD influences the gut microbiota, the potential implications of these alterations for cancer patients remain to be fully understood. Furthermore, the use of FMD is not recommended for individuals with high susceptibility to malnutrition and cachexia, emphasizing the need for specialized monitoring and prudent implementation. Ultimately, while FMD cannot replace standard cancer treatments, it may serve as a complementary approach to support gut health by promoting beneficial bacterial strains. This could contribute to a more balanced intestinal microbiota, potentially offering an alternative to probiotics within a broader, multimodal strategy aimed at improving the overall health of individuals at risk for or affected by intestinal cancers. However, further research is needed to fully understand its clinical implications and long-term effects.

## Supporting information

Supplementary Table 2. Metagenomic analyses

## ACKNOWLEDGEMENTS

This work was supported by Fondazione AIRC (IG 2018—ID. 21963 project, PI: F.P. This project has partially received funding from the European Union`s Horizon 2020 research and innovation program under grant agreement No 825410 (ONCOBIOME project to AN, BP, and ST).

N.E.C is supported by an EMBO Postdoctoral Fellowship (ALTF 498-2023).

## Supplemental data

**Figure S1.**
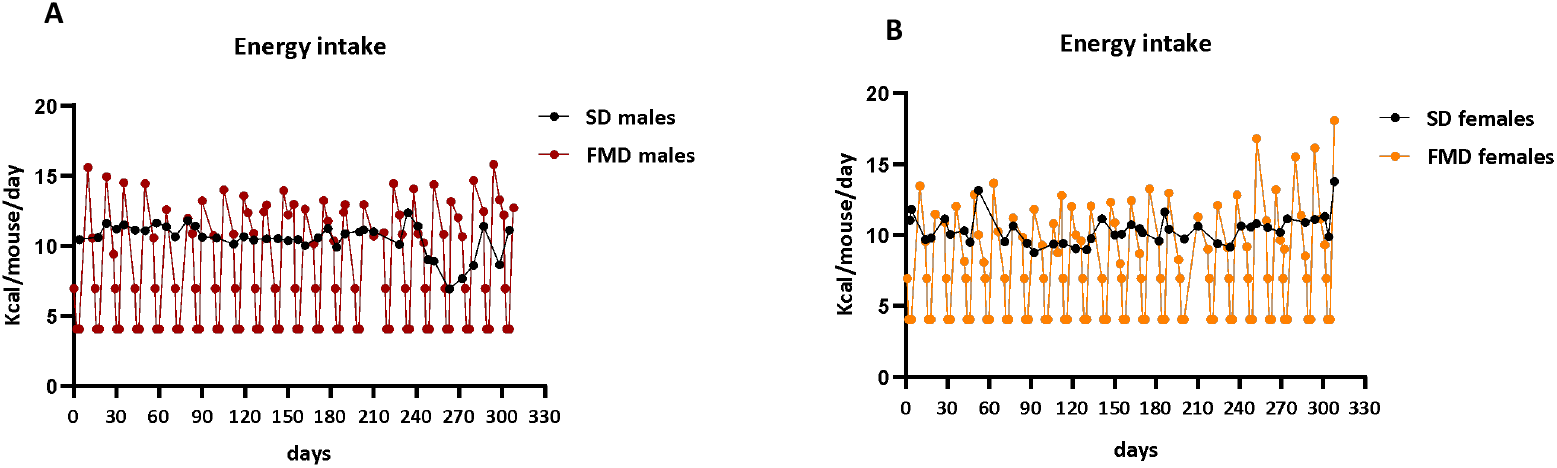
**A-B)** Energy intake measured in VCM+SD or VCM+FMD stratified for **A)** males and **B)** females. Energy intake is expressed as average kcal/mouse/day. VCM+SD males; n=12 and females; n=12. VCM+FMD diet males; n=14 and females; n=12.

**Figure S2.**
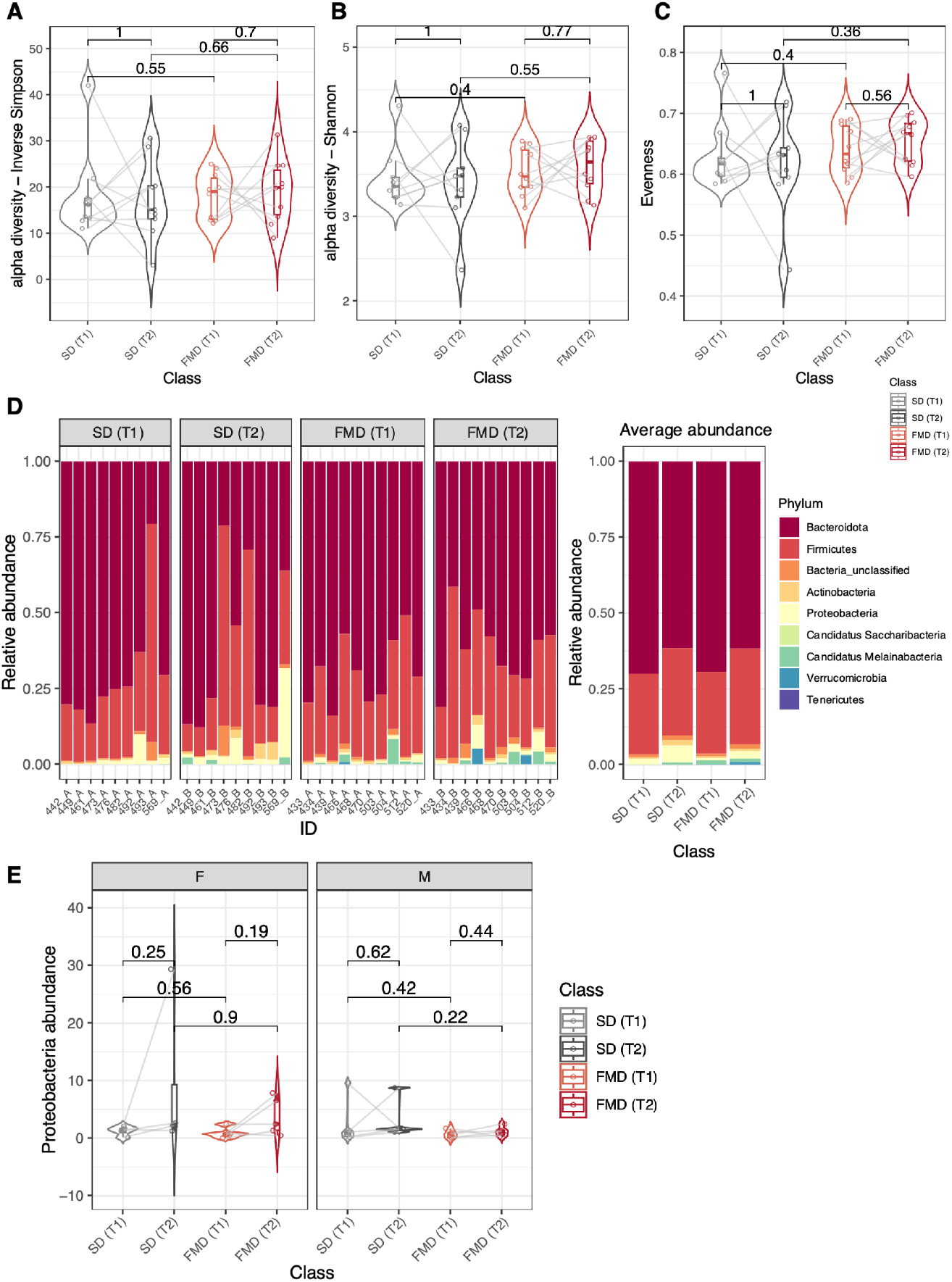
Microbial diversity and phyla relative abundances. **A-C)** Violin plots reporting the alpha diversity computed as **A)** Inverse Simpson index and **B)** Shannon index. **C)** Violin plots of Evenness in VCM+SD (n= 9 all animals) and VCM+FMD mice. (n=10; all animals). Evenness measures how evenly the microbial abundance is distributed in a sample without considering the number of species. Statistical analysis was performed using the Vegan R package, p-value by Wilcoxon Rank-Sum test. **D)** Bar plots showing the samples-specific (left) and the average (right) relative abundance of microbial Phyla as measured in samples collected from mice fed with SD or with FMD at T1 and T2. **E)** Violin plots showing the relative abundance of Proteobacteria among the dietary groups stratified by sex. Abundancies are expressed as % of the total Phylum. Significance of the differences by Wilcoxon Rank-Sum test. Paired analysis was performed for the T2 vs T1 comparisons.

**Supplementary Table 1.**
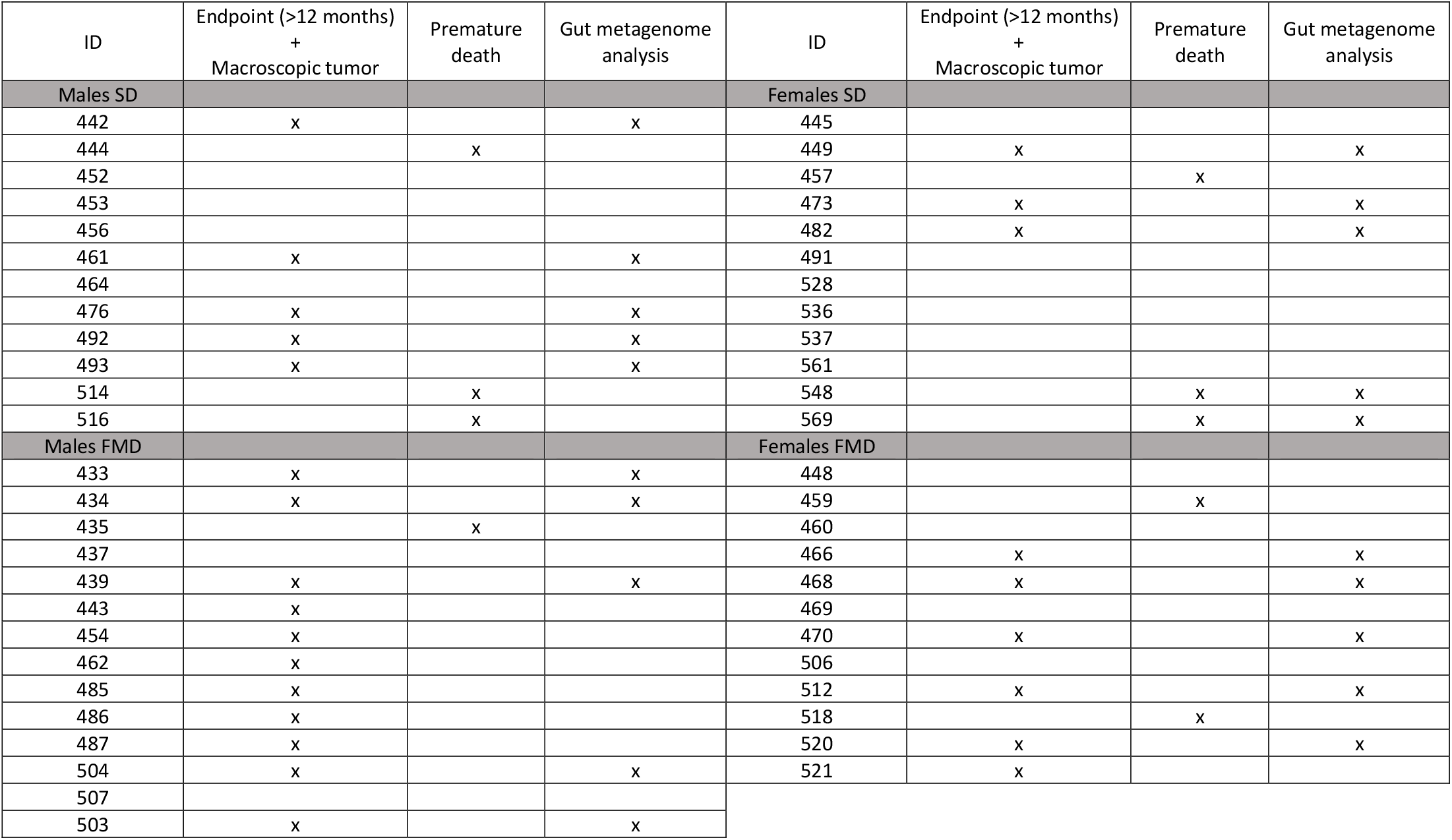
Animal data.

