## Supplementary Table 2. Metagenomic analyses for "Fasting-mimicking diet counteracts gut microbial dysbiosis in experimental Lynch syndrome"

### A_Summary statistics

| ID | Model | Diet | Time | Sex | Toremove | Class |
|---|---|---|---|---|---|---|
| 433_A | 433 | FMD | T1 | M | No | FMD (T1) |
| 433_B | 433 | FMD | T2 | M | No | FMD (T2) |
| 434_A | 434 | FMD | T1 | M | No | FMD (T1) |
| 434_B | 434 | FMD | T2 | M | No | FMD (T2) |
| 439_A | 439 | FMD | T1 | M | No | FMD (T1) |
| 439_B | 439 | FMD | T2 | M | No | FMD (T2) |
| 442_A | 442 | SD | T1 | M | No | SD (T1) |
| 442_B | 442 | SD | T2 | M | No | SD (T2) |
| 449_A | 449 | SD | T1 | F | No | SD (T1) |
| 449_B | 449 | SD | T2 | F | No | SD (T2) |
| 461_A | 461 | SD | T1 | M | No | SD (T1) |
| 461_B | 461 | SD | T2 | M | No | SD (T2) |
| 466_A | 466 | FMD | T1 | F | No | FMD (T1) |
| 466_B | 466 | FMD | T2 | F | No | FMD (T2) |
| 468_A | 468 | FMD | T1 | F | No | FMD (T1) |
| 468_B | 468 | FMD | T2 | F | No | FMD (T2) |
| 470_A | 470 | FMD | T1 | F | No | FMD (T1) |
| 470_B | 470 | FMD | T2 | F | No | FMD (T2) |
| 473_A | 473 | SD | T1 | F | No | SD (T1) |
| 473_B | 473 | SD | T2 | F | No | SD (T2) |
| 476_A | 476 | SD | T1 | M | No | SD (T1) |
| 476_B | 476 | SD | T2 | M | No | SD (T2) |
| 482_A | 482 | SD | T1 | F | No | SD (T1) |
| 482_B | 482 | SD | T2 | F | No | SD (T2) |
| 492_A | 492 | SD | T1 | M | No | SD (T1) |
| 492_B | 492 | SD | T2 | M | No | SD (T2) |
| 493_A | 493 | SD | T1 | M | No | SD (T1) |
| 493_B | 493 | SD | T2 | M | No | SD (T2) |
| 503_A | 503 | FMD | T1 | M | No | FMD (T1) |
| 503_B | 503 | FMD | T2 | M | No | FMD (T2) |
| 504_A | 504 | FMD | T1 | M | No | FMD (T1) |
| 504_B | 504 | FMD | T2 | M | No | FMD (T2) |
| 512_A | 512 | FMD | T1 | F | No | FMD (T1) |
| 512_B | 512 | FMD | T2 | F | No | FMD (T2) |
| 520_A | 520 | FMD | T1 | F | No | FMD (T1) |
| 520_B | 520 | FMD | T2 | F | No | FMD (T2) |
| 569_A | 569 | SD | T1 | F | No | SD (T1) |
| 569_B | 569 | SD | T2 | F | No | SD (T2) |

| Richness | Diversity_inverse_simpson | Diversity_shannon | Evenness_simpson |
|---|---|---|---|
| 265 | 18,56 | 3,45 | 0,62 |
| 192 | 13,56 | 3,14 | 0,60 |
| 273 | 25,01 | 3,86 | 0,69 |
| 246 | 9,02 | 3,44 | 0,63 |
| 226 | 19,54 | 3,50 | 0,65 |
| 265 | 19,82 | 3,78 | 0,68 |
| 238 | 13,33 | 3,22 | 0,59 |
| 225 | 20,28 | 3,48 | 0,64 |
| 192 | 12,76 | 3,14 | 0,60 |
| 166 | 13,16 | 3,10 | 0,61 |
| 250 | 11,07 | 3,23 | 0,58 |
| 246 | 20,05 | 3,48 | 0,63 |
| 280 | 22,62 | 3,89 | 0,69 |
| 177 | 11,98 | 3,18 | 0,61 |
| 262 | 24,13 | 3,80 | 0,68 |
| 285 | 24,68 | 3,94 | 0,70 |
| 199 | 12,84 | 3,10 | 0,59 |
| 311 | 24,72 | 3,93 | 0,69 |
| 229 | 16,63 | 3,36 | 0,62 |
| 290 | 28,78 | 4,08 | 0,72 |
| 246 | 16,26 | 3,41 | 0,62 |
| 285 | 30,60 | 4,03 | 0,71 |
| 251 | 17,66 | 3,46 | 0,63 |
| 209 | 3,11 | 2,37 | 0,44 |
| 238 | 21,90 | 3,66 | 0,67 |
| 262 | 15,14 | 3,32 | 0,60 |
| 277 | 42,05 | 4,31 | 0,77 |
| 253 | 14,17 | 3,23 | 0,58 |
| 218 | 12,19 | 3,35 | 0,62 |
| 194 | 20,03 | 3,50 | 0,67 |
| 263 | 19,92 | 3,74 | 0,67 |
| 229 | 15,67 | 3,37 | 0,62 |
| 247 | 13,46 | 3,35 | 0,61 |
| 266 | 31,35 | 3,91 | 0,70 |
| 231 | 12,88 | 3,24 | 0,60 |
| 298 | 20,83 | 3,81 | 0,67 |
| 242 | 13,63 | 3,31 | 0,60 |
| 280 | 10,58 | 3,57 | 0,63 |

### B_Permanova results

| Variable | Df | SumOfSqs | R2 | F | Pr(>F) |
|---|---|---|---|---|---|
| Diet | 1 | 0,190106967 | 0,03677251 | 1,52001297 | 0,087 |
| Time | 1 | 0,433975632 | 0,08394418 | 3,4698812 | 0,001 |
| Sex | 1 | 0,167162144 | 0,03233428 | 1,3365561 | 0,158 |
| Model | 1 | 0,251280023 | 0,04860525 | 2,00912623 | 0,028 |
| Residual | 33 | 4,127287093 | 0,79834377 | NA | NA |
| Total | 37 | 5,169811859 | 1 | NA | NA |

#### C_Species analysis

| | Mean | | |
|---|---|---|---|
| **ID** | **SD (T1)** | **SD (T2)** | **FMD (T1)** |
| Escherichia coli (SGB10068) | 0,03 | 4,59 | 0,05 |
| Bacteroidales bacterium (SGB21441) | 6,66 | 2,86 | 5,56 |
| Lactobacillus johnsonii (SGB7041) | 6,71 | 1,30 | 7,36 |
| Ligilactobacillus murinus (SGB7077) | 0,80 | 0,15 | 1,88 |
| Limosilactobacillus reuteri (SGB7095) | 1,07 | 0,39 | 1,27 |
| Parabacteroides goldsteinii (SGB1941) | 0,11 | 0,50 | 0,11 |
| Adlercreutzia mucosicola (SGB33554) | 0,04 | 0,03 | 0,04 |
| Adlercreutzia muris (SGB33553) | 0,03 | 0,02 | 0,03 |
| Anaerofustis stercorihominis (SGB49058) | 0,00 | 0,00 | 0,00 |
| Bacteria unclassified_SGB43541 (SGB43541) | 0,01 | 0,03 | 0,01 |
| bacterium 1xD42_87 (SGB29426) | 0,01 | 0,24 | 0,11 |
| bacterium c_19 (SGB77368) | 0,07 | 0,00 | 0,08 |
| Bacteroidales bacterium (SGB21445) | 2,10 | 0,76 | 1,38 |
| Bacteroidales bacterium (SGB27761) | 14,22 | 7,34 | 12,52 |
| Bacteroidales bacterium (SGB35903) | 1,67 | 0,94 | 2,45 |
| Clostridia bacterium (SGB41482) | 0,04 | 0,04 | 0,08 |
| Clostridiaceae bacterium (SGB41675) | 0,11 | 0,03 | 0,05 |
| Clostridiaceae bacterium (SGB43075) | 0,07 | 0,04 | 0,04 |
| Clostridium perfringens (SGB6191) | 0,00 | 0,00 | 0,00 |
| Dorea sp_5_2 (SGB7275) | 0,01 | 0,01 | 0,01 |
| Duncaniella freteri (SGB27768) | 0,26 | 0,29 | 0,44 |
| Lachnospiraceae bacterium (SGB40904) | 0,11 | 0,11 | 0,03 |
| Lachnospiraceae bacterium (SGB41402) | 0,01 | 0,00 | 0,02 |
| Lachnospiraceae bacterium (SGB41416) | 0,01 | 0,00 | 0,00 |
| Lachnospiraceae bacterium (SGB59842) | 0,00 | 0,00 | 0,00 |
| Neglectibacter sp_X4 (SGB43257) | 0,00 | 0,01 | 0,00 |
| Oscillibacter SGB43496 (SGB43496) | 0,04 | 0,05 | 0,01 |
| Oscillospiraceae unclassified_SGB43502 (SGB43502) | 0,03 | 0,02 | 0,01 |
| Paramuribaculum intestinale (SGB21584) | 0,00 | 0,00 | 0,96 |
| Staphylococcus nepalensis (SGB7839) | 0,01 | 0,01 | 0,01 |
| Streptococcus danieliae (SGB82603) | 0,00 | 0,00 | 0,00 |
| Acetatifactor SGB41545 (SGB41545) | 0,00 | 0,01 | 0,00 |
| Acetatifactor SGB41546 (SGB41546) | 0,03 | 0,03 | 0,02 |
| Acetatifactor SGB41549 (SGB41549) | 0,00 | 0,00 | 0,01 |
| Acetatifactor SGB94872 (SGB94872) | 0,00 | 0,01 | 0,00 |
| Acetatifactor sp_DSM_110981 (SGB41548) | 0,02 | 0,02 | 0,02 |
| Acutalibacter muris (SGB7269) | 0,05 | 0,04 | 0,06 |
| Acutalibacter SGB43254 (SGB43254) | 0,00 | 0,00 | 0,00 |
| Acutalibacter sp_1XD8_33 (SGB29419) | 0,00 | 0,00 | 0,00 |
| Adlercreutzia caecimuris (SGB14802) | 0,10 | 0,12 | 0,14 |
| Adlercreutzia sp_DSM_108611 (SGB86230) | 0,01 | 0,01 | 0,01 |
| Akkermansia muciniphila (SGB9226) | 0,00 | 0,00 | 0,07 |
| Anaerocaecibacter sp_CLA_AA_M11 (SGB41016) | 0,00 | 0,02 | 0,00 |
| Anaerotruncus colihominis (SGB82507) | 0,00 | 0,00 | 0,00 |

| | | | |
|---|---|---|---|
| Anaerotruncus sp_1XD42_93 (SGB43003) | 0,04 | 0,08 | 0,10 |
| Bacteria unclassified_SGB102200 (SGB102200) | 0,01 | 0,03 | 0,01 |
| Bacteria unclassified_SGB41677 (SGB41677) | 0,21 | 0,02 | 0,06 |
| Bacteria unclassified_SGB43539 (SGB43539) | 0,01 | 0,01 | 0,00 |
| Bacteria unclassified_SGB43544 (SGB43544) | 0,00 | 0,00 | 0,01 |
| Bacteria unclassified_SGB43546 (SGB43546) | 0,01 | 0,03 | 0,01 |
| Bacteria unclassified_SGB63211 (SGB63211) | 0,03 | 0,01 | 0,01 |
| bacterium 0_1xD8_82 (SGB29425) | 0,02 | 0,01 | 0,00 |
| bacterium 1XD21_13 (SGB82510) | 0,01 | 0,00 | 0,00 |
| bacterium 1XD21_70 (SGB29423) | 0,00 | 0,00 | 0,00 |
| bacterium 1XD42_1 (SGB29420) | 0,05 | 0,09 | 0,04 |
| bacterium 1XD42_54 (SGB29418) | 0,01 | 0,01 | 0,01 |
| bacterium 1xD42_62 (SGB82509) | 0,00 | 0,00 | 0,00 |
| bacterium 1xD42_67 (SGB29335) | 0,00 | 0,00 | 0,00 |
| bacterium 1XD42_76 (SGB41647) | 0,01 | 0,02 | 0,01 |
| bacterium 1xD8_48 (SGB41627) | 0,06 | 0,04 | 0,02 |
| bacterium 1xD8_6 (SGB29416) | 0,29 | 0,67 | 0,20 |
| bacterium 1XD8_76 (SGB29429) | 0,00 | 0,00 | 0,01 |
| bacterium D16_76 (SGB43258) | 0,00 | 0,00 | 0,00 |
| Bacteroidales bacterium (SGB35892) | 0,84 | 0,75 | 0,58 |
| Bacteroidales bacterium (SGB40205) | 1,06 | 1,26 | 1,13 |
| Bacteroidales bacterium (SGB40306) | 0,73 | 0,47 | 0,62 |
| Bacteroides acidifaciens (SGB27814) | 1,61 | 2,05 | 1,57 |
| Bacteroides caecimuris (SGB1863) | 0,19 | 0,21 | 0,21 |
| Bacteroides muris (SGB27790) | 3,17 | 3,87 | 4,54 |
| Candidatus Arthromitus_sp_SFB_mouse (SGB8526) | 0,01 | 0,02 | 0,00 |
| Clostridia bacterium (SGB102314) | 0,01 | 0,00 | 0,01 |
| Clostridia bacterium (SGB109668) | 0,00 | 0,00 | 0,00 |
| Clostridia bacterium (SGB109669) | 0,00 | 0,00 | 0,00 |
| Clostridia bacterium (SGB109671) | 0,01 | 0,02 | 0,01 |
| Clostridia bacterium (SGB41508) | 0,03 | 0,02 | 0,06 |
| Clostridia bacterium (SGB41510) | 0,01 | 0,01 | 0,01 |
| Clostridia bacterium (SGB41594) | 0,00 | 0,01 | 0,01 |
| Clostridia bacterium (SGB41692) | 0,03 | 0,04 | 0,02 |
| Clostridia bacterium (SGB45231) | 0,10 | 0,15 | 0,14 |
| Clostridia bacterium (SGB82506) | 0,00 | 0,00 | 0,00 |
| Clostridiaceae bacterium (SGB41251) | 0,03 | 0,01 | 0,01 |
| Clostridiaceae bacterium (SGB41645) | 0,04 | 0,08 | 0,04 |
| Clostridiaceae bacterium (SGB41667) | 0,02 | 0,02 | 0,01 |
| Clostridiaceae unclassified_SGB41663 (SGB41663) | 0,03 | 0,07 | 0,10 |
| Clostridiales bacterium (SGB42493) | 0,18 | 0,06 | 0,26 |
| Clostridiales bacterium (SGB59899) | 0,16 | 0,09 | 0,19 |
| Clostridiales bacterium (SGB82513) | 0,00 | 0,01 | 0,00 |
| Clostridium cocleatum (SGB6748) | 0,02 | 0,05 | 0,04 |
| Clostridium cuniculi (SGB29095) | 0,00 | 0,80 | 0,00 |
| Clostridium SGB65123 (SGB65123) | 0,00 | 0,00 | 0,00 |

| | | | |
|---|---|---|---|
| Coriobacteriaceae bacterium (SGB42958) | 0,04 | 0,03 | 0,04 |
| Duncaniella dubosii (SGB2089) | 8,37 | 4,60 | 8,58 |
| Duncaniella muris (SGB21446) | 2,98 | 2,26 | 2,87 |
| Enterococcus faecalis (SGB7962) | 0,01 | 1,07 | 0,11 |
| Erysipelotrichales bacterium (SGB44461) | 0,03 | 0,03 | 0,11 |
| Erysipelotrichales bacterium (SGB44463) | 0,01 | 0,02 | 0,02 |
| Erysipelotrichales bacterium (SGB44464) | 0,00 | 0,00 | 0,00 |
| Eubacteriaceae bacterium (SGB41268) | 0,00 | 0,00 | 0,00 |
| Eubacteriaceae bacterium (SGB41444) | 0,20 | 0,05 | 0,14 |
| Eubacteriaceae bacterium (SGB41701) | 0,22 | 0,04 | 0,07 |
| Eubacteriaceae bacterium (SGB41703) | 0,14 | 0,11 | 0,05 |
| Fumia xinanensis (SGB25547) | 0,00 | 0,00 | 0,00 |
| Granulimonas faecalis (SGB33580) | 0,16 | 1,55 | 0,23 |
| Jeotgalicoccus halotolerans (SGB24956) | 0,00 | 0,00 | 0,00 |
| Lachnospiraceae bacterium (SGB109670) | 0,00 | 0,00 | 0,00 |
| Lachnospiraceae bacterium (SGB109672) | 0,00 | 0,00 | 0,00 |
| Lachnospiraceae bacterium (SGB29415) | 0,00 | 0,00 | 0,00 |
| Lachnospiraceae bacterium (SGB41239) | 0,00 | 0,04 | 0,01 |
| Lachnospiraceae bacterium (SGB41401) | 0,04 | 0,02 | 0,15 |
| Lachnospiraceae bacterium (SGB41411) | 0,01 | 0,03 | 0,02 |
| Lachnospiraceae bacterium (SGB41412) | 0,01 | 0,00 | 0,00 |
| Lachnospiraceae bacterium (SGB41415) | 0,00 | 0,11 | 0,02 |
| Lachnospiraceae bacterium (SGB41417) | 0,05 | 0,03 | 0,03 |
| Lachnospiraceae bacterium (SGB41420) | 0,01 | 0,01 | 0,02 |
| Lachnospiraceae bacterium (SGB41421) | 0,00 | 0,00 | 0,01 |
| Lachnospiraceae bacterium (SGB41425) | 0,00 | 0,00 | 0,00 |
| Lachnospiraceae bacterium (SGB41427) | 0,00 | 0,00 | 0,00 |
| Lachnospiraceae bacterium (SGB41461) | 0,03 | 0,02 | 0,07 |
| Lachnospiraceae bacterium (SGB41462) | 0,00 | 0,00 | 0,00 |
| Lachnospiraceae bacterium (SGB41475) | 0,00 | 0,07 | 0,01 |
| Lachnospiraceae bacterium (SGB41480) | 0,14 | 0,04 | 0,05 |
| Lachnospiraceae bacterium (SGB41513) | 0,11 | 0,08 | 0,04 |
| Lachnospiraceae bacterium (SGB41516) | 0,03 | 0,04 | 0,01 |
| Lachnospiraceae bacterium (SGB41519) | 0,03 | 0,01 | 0,01 |
| Lachnospiraceae bacterium (SGB41569) | 0,23 | 0,16 | 0,10 |
| Lachnospiraceae bacterium (SGB41586) | 0,22 | 0,69 | 0,12 |
| Lachnospiraceae bacterium (SGB41609) | 0,01 | 0,04 | 0,07 |
| Lachnospiraceae bacterium (SGB41617) | 0,05 | 0,01 | 0,01 |
| Lachnospiraceae bacterium (SGB41629) | 0,08 | 0,02 | 0,10 |
| Lachnospiraceae bacterium (SGB41654) | 0,03 | 0,02 | 0,05 |
| Lachnospiraceae bacterium (SGB41672) | 0,00 | 0,03 | 0,02 |
| Lachnospiraceae bacterium (SGB59798) | 0,00 | 0,01 | 0,01 |
| Lachnospiraceae bacterium (SGB7268) | 0,23 | 0,17 | 0,32 |
| Lachnospiraceae bacterium (SGB7271) | 0,00 | 0,02 | 0,00 |
| Lachnospiraceae bacterium (SGB7272) | 0,55 | 0,01 | 0,08 |
| Lachnospiraceae bacterium (SGB94870) | 0,00 | 0,00 | 0,00 |

| | | | |
|---|---|---|---|
| Lachnospiraceae bacterium_A2 (SGB7277) | 0,08 | 0,09 | 0,06 |
| Lachnospiraceae bacterium_MD308 (SGB7279) | 0,00 | 0,00 | 0,00 |
| Lachnospiraceae bacterium_MD329 (SGB42354) | 0,03 | 0,01 | 0,01 |
| Lachnospiraceae bacterium_MD335 (SGB5155) | 0,14 | 0,04 | 0,03 |
| Lachnospiraceae unclassified_SGB36958 (SGB36958) | 0,00 | 6,37 | 0,03 |
| Lachnospiraceae unclassified_SGB41413 (SGB41413) | 0,00 | 0,00 | 0,01 |
| Lachnospiraceae unclassified_SGB41418 (SGB41418) | 0,00 | 0,00 | 0,00 |
| Lachnospiraceae unclassified_SGB41424 (SGB41424) | 0,01 | 0,02 | 0,02 |
| Lachnospiraceae unclassified_SGB41521 (SGB41521) | 0,02 | 0,05 | 0,03 |
| Lachnospiraceae unclassified_SGB41589 (SGB41589) | 0,03 | 0,01 | 0,03 |
| Lactobacillus taiwanensis (SGB7040) | 1,84 | 1,15 | 1,25 |
| Muribaculaceae bacterium (SGB21431) | 0,96 | 0,94 | 1,12 |
| Muribaculaceae bacterium (SGB27763) | 5,94 | 6,32 | 6,22 |
| Muribaculaceae bacterium (SGB35953) | 0,50 | 0,68 | 0,45 |
| Muribaculaceae bacterium (SGB40323) | 1,33 | 1,33 | 1,19 |
| Muribaculaceae bacterium_Isolate_013_NCI (SGB2776 | 0,35 | 0,37 | 0,38 |
| Muribaculum intestinale (SGB2088) | 3,45 | 6,28 | 3,17 |
| Oscillospiraceae bacterium (SGB102199) | 0,04 | 0,00 | 0,01 |
| Oscillospiraceae bacterium (SGB109673) | 0,00 | 0,00 | 0,00 |
| Oscillospiraceae bacterium (SGB43504) | 0,09 | 0,05 | 0,06 |
| Oscillospiraceae bacterium (SGB43532) | 0,05 | 0,05 | 0,04 |
| Oscillospiraceae bacterium (SGB63364) | 0,01 | 0,01 | 0,01 |
| Oscillospiraceae bacterium (SGB95005) | 0,00 | 0,00 | 0,00 |
| Oscillospiraceae unclassified_SGB43505 (SGB43505) | 0,00 | 0,00 | 0,00 |
| Oscillospiraceae unclassified_SGB43542 (SGB43542) | 0,00 | 0,00 | 0,00 |
| Paeniclostridium sordellii (SGB6146) | 0,00 | 0,05 | 0,00 |
| Pumilibacter sp_CLA_AA_M08 (SGB41021) | 0,00 | 0,01 | 0,00 |
| Pumilibacter sp_CLA_AA_M10 (SGB41020) | 0,02 | 0,01 | 0,02 |
| Rikenellaceae bacterium (SGB35894) | 3,73 | 3,58 | 2,11 |
| Romboutsia ilealis (SGB24891) | 0,00 | 0,16 | 0,01 |
| Schaedlerella arabinosiphila (SGB7270) | 0,01 | 0,00 | 0,01 |
| Sporofaciens musculi (SGB41410) | 0,08 | 0,01 | 0,01 |
| Turicibacter sp_1E2 (SGB39153) | 0,29 | 0,48 | 0,56 |
| Turicimonas muris (SGB9258) | 0,08 | 0,01 | 0,00 |
| Xylanibacter rodentium (SGB40522) | 1,06 | 0,58 | 0,74 |

| | Standard deviation | | | | Preva | |
|---|---|---|---|---|---|---|
| FMD (T2) | SD (T1) | SD (T2) | FMD (T1) | FMD (T2) | SD (T1) | SD (T2) |
| 1,22 | 0,04 | 8,57 | 0,15 | 2,40 | 0,78 | 1,00 |
| 2,00 | 2,33 | 2,10 | 2,32 | 1,41 | 1,00 | 1,00 |
| 4,40 | 3,44 | 1,00 | 5,04 | 3,48 | 1,00 | 1,00 |
| 0,65 | 0,57 | 0,07 | 3,56 | 0,65 | 1,00 | 1,00 |
| 1,41 | 0,72 | 0,24 | 0,92 | 0,76 | 1,00 | 1,00 |
| 0,30 | 0,07 | 0,47 | 0,11 | 0,32 | 1,00 | 1,00 |
| 0,06 | 0,04 | 0,04 | 0,02 | 0,04 | 1,00 | 1,00 |
| 0,04 | 0,01 | 0,02 | 0,02 | 0,02 | 1,00 | 0,89 |
| 0,00 | 0,01 | 0,00 | 0,01 | 0,00 | 0,78 | 0,11 |
| 0,01 | 0,01 | 0,04 | 0,01 | 0,01 | 0,78 | 1,00 |
| 0,16 | 0,01 | 0,49 | 0,21 | 0,23 | 0,44 | 0,89 |
| 0,16 | 0,09 | 0,01 | 0,10 | 0,18 | 0,78 | 0,33 |
| 0,83 | 1,20 | 0,75 | 1,16 | 0,71 | 1,00 | 1,00 |
| 7,15 | 6,90 | 6,05 | 5,51 | 4,50 | 1,00 | 1,00 |
| 1,87 | 1,64 | 1,02 | 1,89 | 1,43 | 1,00 | 1,00 |
| 0,10 | 0,02 | 0,02 | 0,05 | 0,06 | 1,00 | 1,00 |
| 0,07 | 0,13 | 0,05 | 0,09 | 0,05 | 1,00 | 0,78 |
| 0,02 | 0,12 | 0,07 | 0,04 | 0,02 | 0,78 | 1,00 |
| 3,80 | 0,00 | 0,00 | 0,00 | 7,55 | 0,00 | 0,00 |
| 0,04 | 0,01 | 0,02 | 0,02 | 0,05 | 1,00 | 0,89 |
| 0,53 | 0,22 | 0,11 | 0,37 | 0,29 | 1,00 | 1,00 |
| 0,03 | 0,09 | 0,17 | 0,05 | 0,03 | 1,00 | 0,89 |
| 0,01 | 0,01 | 0,00 | 0,04 | 0,02 | 0,33 | 0,11 |
| 0,00 | 0,02 | 0,01 | 0,01 | 0,00 | 0,44 | 0,67 |
| 0,00 | 0,00 | 0,01 | 0,00 | 0,01 | 0,89 | 0,44 |
| 0,00 | 0,01 | 0,01 | 0,01 | 0,00 | 0,56 | 0,78 |
| 0,00 | 0,09 | 0,05 | 0,03 | 0,01 | 0,78 | 0,89 |
| 0,00 | 0,05 | 0,04 | 0,01 | 0,01 | 1,00 | 0,78 |
| 2,60 | 0,00 | 0,00 | 1,75 | 2,66 | 0,00 | 0,00 |
| 0,00 | 0,03 | 0,02 | 0,01 | 0,00 | 0,67 | 0,22 |
| 0,00 | 0,01 | 0,00 | 0,00 | 0,00 | 0,89 | 0,67 |
| 0,00 | 0,00 | 0,01 | 0,01 | 0,00 | 0,22 | 0,56 |
| 0,02 | 0,05 | 0,05 | 0,02 | 0,03 | 0,89 | 1,00 |
| 0,00 | 0,00 | 0,00 | 0,03 | 0,01 | 0,11 | 0,11 |
| 0,00 | 0,00 | 0,01 | 0,01 | 0,01 | 0,33 | 0,22 |
| 0,01 | 0,03 | 0,03 | 0,03 | 0,02 | 0,67 | 0,67 |
| 0,05 | 0,11 | 0,05 | 0,07 | 0,06 | 0,89 | 1,00 |
| 0,00 | 0,00 | 0,00 | 0,00 | 0,01 | 0,11 | 0,11 |
| 0,00 | 0,00 | 0,00 | 0,00 | 0,00 | 0,33 | 0,33 |
| 0,12 | 0,05 | 0,08 | 0,07 | 0,05 | 1,00 | 1,00 |
| 0,01 | 0,00 | 0,01 | 0,01 | 0,01 | 1,00 | 0,89 |
| 0,82 | 0,00 | 0,00 | 0,23 | 1,79 | 0,00 | 0,00 |
| 0,01 | 0,00 | 0,03 | 0,00 | 0,01 | 0,67 | 0,67 |
| 0,00 | 0,00 | 0,01 | 0,00 | 0,01 | 0,22 | 0,56 |

| | | | | | | |
|---|---|---|---|---|---|---|
| 0,13 | 0,06 | 0,10 | 0,20 | 0,17 | 1,00 | 1,00 |
| 0,01 | 0,02 | 0,05 | 0,01 | 0,02 | 0,89 | 0,89 |
| 0,04 | 0,55 | 0,03 | 0,05 | 0,05 | 0,89 | 0,67 |
| 0,00 | 0,03 | 0,01 | 0,00 | 0,00 | 0,78 | 0,67 |
| 0,00 | 0,00 | 0,00 | 0,02 | 0,00 | 0,22 | 0,22 |
| 0,01 | 0,03 | 0,05 | 0,02 | 0,01 | 0,44 | 0,89 |
| 0,01 | 0,08 | 0,02 | 0,01 | 0,01 | 1,00 | 1,00 |
| 0,00 | 0,05 | 0,01 | 0,01 | 0,00 | 0,56 | 0,33 |
| 0,01 | 0,02 | 0,01 | 0,00 | 0,01 | 0,89 | 0,78 |
| 0,00 | 0,00 | 0,00 | 0,00 | 0,00 | 0,00 | 0,22 |
| 0,11 | 0,10 | 0,11 | 0,04 | 0,22 | 0,89 | 1,00 |
| 0,02 | 0,01 | 0,01 | 0,00 | 0,02 | 1,00 | 1,00 |
| 0,00 | 0,00 | 0,00 | 0,00 | 0,00 | 0,00 | 0,00 |
| 0,00 | 0,00 | 0,00 | 0,00 | 0,00 | 0,22 | 0,33 |
| 0,02 | 0,02 | 0,02 | 0,01 | 0,02 | 1,00 | 0,89 |
| 0,02 | 0,17 | 0,07 | 0,03 | 0,02 | 0,78 | 0,89 |
| 0,71 | 0,49 | 1,89 | 0,32 | 1,25 | 0,78 | 0,78 |
| 0,01 | 0,00 | 0,00 | 0,04 | 0,02 | 0,00 | 0,00 |
| 0,00 | 0,00 | 0,01 | 0,00 | 0,01 | 0,33 | 0,22 |
| 0,51 | 0,76 | 0,92 | 0,42 | 0,38 | 1,00 | 1,00 |
| 1,29 | 0,45 | 1,00 | 1,06 | 0,96 | 1,00 | 1,00 |
| 0,60 | 0,30 | 0,30 | 0,20 | 0,32 | 1,00 | 1,00 |
| 1,81 | 1,44 | 1,41 | 1,19 | 1,81 | 1,00 | 1,00 |
| 0,34 | 0,11 | 0,14 | 0,13 | 0,23 | 1,00 | 1,00 |
| 5,07 | 1,79 | 2,87 | 1,56 | 4,37 | 1,00 | 1,00 |
| 0,04 | 0,02 | 0,03 | 0,01 | 0,08 | 0,67 | 0,56 |
| 0,01 | 0,02 | 0,01 | 0,01 | 0,01 | 0,44 | 0,56 |
| 0,00 | 0,00 | 0,00 | 0,00 | 0,00 | 0,44 | 0,67 |
| 0,00 | 0,00 | 0,01 | 0,01 | 0,00 | 0,44 | 0,33 |
| 0,01 | 0,01 | 0,02 | 0,02 | 0,01 | 1,00 | 1,00 |
| 0,02 | 0,03 | 0,02 | 0,10 | 0,02 | 1,00 | 0,89 |
| 0,03 | 0,01 | 0,02 | 0,01 | 0,07 | 0,44 | 0,89 |
| 0,01 | 0,01 | 0,01 | 0,02 | 0,03 | 0,56 | 0,78 |
| 0,02 | 0,04 | 0,06 | 0,03 | 0,02 | 1,00 | 0,78 |
| 0,50 | 0,06 | 0,15 | 0,14 | 0,66 | 1,00 | 1,00 |
| 0,00 | 0,00 | 0,00 | 0,00 | 0,00 | 0,56 | 0,44 |
| 0,01 | 0,05 | 0,02 | 0,01 | 0,01 | 0,78 | 0,67 |
| 0,12 | 0,03 | 0,09 | 0,03 | 0,11 | 1,00 | 0,89 |
| 0,01 | 0,01 | 0,02 | 0,01 | 0,01 | 1,00 | 1,00 |
| 0,07 | 0,02 | 0,08 | 0,10 | 0,08 | 1,00 | 1,00 |
| 0,12 | 0,23 | 0,10 | 0,36 | 0,17 | 0,89 | 0,89 |
| 0,12 | 0,10 | 0,08 | 0,15 | 0,12 | 1,00 | 1,00 |
| 0,00 | 0,00 | 0,01 | 0,00 | 0,00 | 0,67 | 0,89 |
| 0,23 | 0,03 | 0,08 | 0,05 | 0,50 | 0,67 | 0,67 |
| 0,06 | 0,00 | 1,70 | 0,00 | 0,13 | 0,00 | 0,44 |
| 0,03 | 0,00 | 0,00 | 0,00 | 0,09 | 0,11 | 0,00 |

| | | | | | | |
|---|---|---|---|---|---|---|
| 0,05 | 0,02 | 0,01 | 0,01 | 0,02 | 1,00 | 1,00 |
| 6,93 | 4,57 | 3,63 | 4,78 | 3,28 | 1,00 | 1,00 |
| 2,74 | 1,38 | 1,90 | 1,55 | 1,20 | 1,00 | 1,00 |
| 0,19 | 0,01 | 1,34 | 0,34 | 0,40 | 0,56 | 0,67 |
| 0,03 | 0,06 | 0,05 | 0,13 | 0,04 | 0,22 | 0,67 |
| 0,01 | 0,01 | 0,04 | 0,04 | 0,01 | 0,44 | 0,67 |
| 0,00 | 0,01 | 0,01 | 0,01 | 0,01 | 0,67 | 0,44 |
| 0,00 | 0,01 | 0,00 | 0,01 | 0,00 | 0,56 | 0,33 |
| 0,30 | 0,33 | 0,08 | 0,34 | 0,73 | 1,00 | 0,89 |
| 0,10 | 0,42 | 0,04 | 0,08 | 0,12 | 0,89 | 0,78 |
| 0,05 | 0,25 | 0,21 | 0,06 | 0,06 | 1,00 | 0,67 |
| 0,01 | 0,00 | 0,00 | 0,00 | 0,02 | 0,00 | 0,00 |
| 0,48 | 0,30 | 2,10 | 0,29 | 0,88 | 1,00 | 0,78 |
| 0,00 | 0,00 | 0,01 | 0,00 | 0,00 | 0,11 | 0,22 |
| 0,00 | 0,00 | 0,00 | 0,00 | 0,00 | 0,67 | 0,56 |
| 0,00 | 0,00 | 0,00 | 0,00 | 0,01 | 0,78 | 0,33 |
| 0,05 | 0,00 | 0,00 | 0,00 | 0,13 | 0,00 | 0,00 |
| 0,02 | 0,01 | 0,06 | 0,01 | 0,02 | 0,78 | 1,00 |
| 0,04 | 0,05 | 0,02 | 0,23 | 0,04 | 1,00 | 1,00 |
| 0,07 | 0,02 | 0,05 | 0,05 | 0,11 | 0,67 | 0,78 |
| 0,00 | 0,02 | 0,00 | 0,01 | 0,00 | 0,44 | 0,33 |
| 0,01 | 0,01 | 0,31 | 0,04 | 0,02 | 0,44 | 0,67 |
| 0,09 | 0,04 | 0,05 | 0,03 | 0,11 | 1,00 | 1,00 |
| 0,02 | 0,02 | 0,02 | 0,03 | 0,04 | 0,89 | 0,89 |
| 0,03 | 0,00 | 0,01 | 0,02 | 0,07 | 0,89 | 0,78 |
| 0,00 | 0,00 | 0,00 | 0,00 | 0,00 | 0,00 | 0,00 |
| 0,00 | 0,00 | 0,00 | 0,00 | 0,00 | 0,89 | 0,67 |
| 0,04 | 0,06 | 0,03 | 0,16 | 0,08 | 0,56 | 0,56 |
| 0,03 | 0,00 | 0,00 | 0,00 | 0,08 | 0,00 | 0,00 |
| 0,02 | 0,01 | 0,14 | 0,03 | 0,05 | 0,33 | 0,78 |
| 0,02 | 0,30 | 0,06 | 0,06 | 0,02 | 0,89 | 0,78 |
| 0,07 | 0,24 | 0,10 | 0,04 | 0,06 | 1,00 | 0,89 |
| 0,01 | 0,06 | 0,08 | 0,01 | 0,01 | 0,89 | 1,00 |
| 0,00 | 0,04 | 0,02 | 0,01 | 0,01 | 0,89 | 0,44 |
| 0,11 | 0,45 | 0,22 | 0,07 | 0,13 | 1,00 | 1,00 |
| 0,05 | 0,30 | 1,53 | 0,29 | 0,12 | 0,67 | 0,67 |
| 0,06 | 0,02 | 0,08 | 0,09 | 0,14 | 0,89 | 1,00 |
| 0,03 | 0,09 | 0,02 | 0,03 | 0,05 | 0,67 | 0,78 |
| 0,06 | 0,18 | 0,03 | 0,23 | 0,14 | 0,78 | 0,78 |
| 0,06 | 0,04 | 0,02 | 0,05 | 0,08 | 1,00 | 0,89 |
| 0,03 | 0,00 | 0,04 | 0,02 | 0,04 | 1,00 | 1,00 |
| 0,03 | 0,01 | 0,02 | 0,01 | 0,07 | 0,78 | 0,78 |
| 0,16 | 0,35 | 0,16 | 0,28 | 0,23 | 0,78 | 1,00 |
| 0,00 | 0,00 | 0,04 | 0,01 | 0,01 | 0,22 | 0,44 |
| 0,09 | 1,43 | 0,01 | 0,21 | 0,13 | 0,56 | 0,67 |
| 0,00 | 0,01 | 0,00 | 0,00 | 0,00 | 0,22 | 0,22 |

| | | | | | | |
|---|---|---|---|---|---|---|
| 0,23 | 0,17 | 0,13 | 0,12 | 0,38 | 1,00 | 0,56 |
| 0,02 | 0,01 | 0,00 | 0,01 | 0,04 | 0,11 | 0,22 |
| 0,03 | 0,06 | 0,02 | 0,02 | 0,06 | 0,67 | 0,78 |
| 0,10 | 0,24 | 0,08 | 0,05 | 0,19 | 0,44 | 0,33 |
| 3,26 | 0,00 | 18,56 | 0,05 | 9,81 | 0,11 | 0,33 |
| 0,01 | 0,00 | 0,00 | 0,02 | 0,01 | 0,22 | 0,22 |
| 0,00 | 0,00 | 0,00 | 0,00 | 0,00 | 0,22 | 0,11 |
| 0,02 | 0,01 | 0,03 | 0,03 | 0,03 | 1,00 | 0,89 |
| 0,03 | 0,03 | 0,08 | 0,04 | 0,07 | 0,89 | 0,89 |
| 0,16 | 0,07 | 0,02 | 0,07 | 0,30 | 0,22 | 0,11 |
| 2,78 | 1,92 | 0,97 | 1,32 | 3,10 | 0,89 | 0,89 |
| 0,53 | 0,75 | 0,77 | 1,01 | 0,27 | 1,00 | 1,00 |
| 8,71 | 3,67 | 4,81 | 4,47 | 4,34 | 1,00 | 1,00 |
| 0,77 | 0,22 | 0,39 | 0,18 | 0,38 | 1,00 | 1,00 |
| 1,04 | 0,78 | 1,37 | 0,63 | 0,55 | 1,00 | 1,00 |
| 0,57 | 0,24 | 0,24 | 0,19 | 0,36 | 1,00 | 1,00 |
| 3,21 | 1,62 | 5,17 | 1,45 | 1,75 | 1,00 | 1,00 |
| 0,00 | 0,12 | 0,00 | 0,03 | 0,00 | 0,78 | 1,00 |
| 0,00 | 0,00 | 0,00 | 0,00 | 0,00 | 0,44 | 0,67 |
| 0,06 | 0,12 | 0,05 | 0,08 | 0,07 | 1,00 | 1,00 |
| 0,04 | 0,07 | 0,04 | 0,05 | 0,07 | 1,00 | 1,00 |
| 0,01 | 0,01 | 0,02 | 0,01 | 0,01 | 0,89 | 1,00 |
| 0,00 | 0,00 | 0,00 | 0,00 | 0,00 | 0,00 | 0,11 |
| 0,00 | 0,00 | 0,00 | 0,00 | 0,00 | 0,00 | 0,33 |
| 0,00 | 0,00 | 0,00 | 0,00 | 0,00 | 0,22 | 0,44 |
| 0,14 | 0,00 | 0,14 | 0,00 | 0,45 | 0,00 | 0,22 |
| 0,00 | 0,00 | 0,01 | 0,00 | 0,01 | 0,56 | 0,56 |
| 0,02 | 0,02 | 0,01 | 0,02 | 0,04 | 1,00 | 1,00 |
| 0,98 | 2,12 | 3,76 | 1,53 | 0,61 | 1,00 | 1,00 |
| 0,13 | 0,00 | 0,22 | 0,01 | 0,25 | 0,11 | 0,44 |
| 0,02 | 0,01 | 0,00 | 0,01 | 0,04 | 0,33 | 0,33 |
| 0,01 | 0,14 | 0,01 | 0,02 | 0,02 | 0,67 | 0,89 |
| 0,28 | 0,34 | 0,45 | 0,68 | 0,37 | 0,89 | 0,67 |
| 0,25 | 0,25 | 0,02 | 0,00 | 0,54 | 0,11 | 0,11 |
| 0,47 | 0,78 | 0,36 | 0,48 | 0,42 | 1,00 | 1,00 |

| lence | | log2FC | | |
|---|---|---|---|---|
| FMD (T1) | FMD (T2) | T2 / T1 (SD) | T2 / T1 (FMD) | FMD / SD (T1) |
| 0,50 | 1,00 | 2,76 | 2,46 | -0,70 |
| 1,00 | 1,00 | -0,49 | -0,56 | -0,10 |
| 1,00 | 1,00 | -0,79 | -0,30 | -0,02 |
| 1,00 | 1,00 | -0,67 | -0,19 | 0,00 |
| 1,00 | 1,00 | -0,41 | 0,10 | 0,06 |
| 1,00 | 1,00 | 0,57 | 0,36 | -0,05 |
| 1,00 | 1,00 | -0,13 | 0,21 | 0,08 |
| 1,00 | 1,00 | -0,36 | 0,12 | 0,05 |
| 0,70 | 0,70 | -1,02 | 0,06 | -0,07 |
| 0,80 | 0,90 | 0,83 | -0,18 | 0,10 |
| 0,90 | 0,90 | 1,66 | 0,45 | 1,17 |
| 0,70 | 0,60 | -1,70 | 0,08 | -0,12 |
| 0,90 | 1,00 | -0,69 | -0,17 | -0,35 |
| 1,00 | 1,00 | -0,49 | -0,27 | -0,08 |
| 1,00 | 1,00 | -0,28 | -0,13 | 0,21 |
| 1,00 | 1,00 | -0,04 | 0,09 | 0,24 |
| 0,90 | 1,00 | -1,23 | 0,59 | -0,66 |
| 1,00 | 0,70 | 0,14 | -0,93 | 0,63 |
| 0,00 | 0,50 | 0,00 | 1,97 | 0,00 |
| 0,90 | 0,90 | -0,06 | 0,79 | -0,05 |
| 1,00 | 1,00 | 0,15 | 0,16 | 0,20 |
| 1,00 | 0,90 | -0,40 | 0,05 | -0,78 |
| 0,60 | 0,50 | -0,48 | 0,24 | 0,29 |
| 0,30 | 0,20 | 0,21 | -0,25 | -0,27 |
| 0,30 | 0,50 | -0,48 | 0,09 | -0,81 |
| 0,30 | 0,50 | 0,73 | -0,06 | -0,36 |
| 0,70 | 0,50 | 0,69 | 0,14 | -0,96 |
| 0,70 | 0,40 | 0,06 | -0,58 | -0,54 |
| 0,30 | 0,70 | 0,00 | 2,07 | 1,04 |
| 0,50 | 0,10 | -0,63 | -0,98 | 0,19 |
| 1,00 | 1,00 | -0,71 | -0,11 | -0,24 |
| 0,20 | 0,30 | 0,85 | 0,00 | 0,05 |
| 1,00 | 0,80 | 0,32 | -0,30 | 0,39 |
| 0,50 | 0,40 | 0,00 | -0,25 | 0,72 |
| 0,50 | 0,60 | 0,05 | 0,25 | 0,52 |
| 0,80 | 0,60 | -0,30 | -0,32 | -0,10 |
| 1,00 | 1,00 | 0,42 | -0,26 | 0,64 |
| 0,20 | 0,20 | 0,02 | 0,16 | 0,08 |
| 0,00 | 0,20 | -0,12 | 0,06 | -0,24 |
| 1,00 | 1,00 | 0,11 | -0,04 | 0,16 |
| 1,00 | 1,00 | -0,14 | 0,26 | -0,09 |
| 0,10 | 0,30 | 0,00 | 0,79 | 0,02 |
| 0,60 | 0,50 | 0,73 | -0,12 | 0,07 |
| 0,10 | 0,30 | 0,90 | 0,25 | -0,01 |

| | | | | |
|---|---|---|---|---|
| 1,00 | 1,00 | 0,40 | 0,36 | 0,20 |
| 0,90 | 0,50 | 0,05 | -0,75 | -0,03 |
| 1,00 | 0,70 | -0,51 | -0,94 | 0,53 |
| 0,60 | 0,60 | -0,35 | -0,22 | -0,36 |
| 0,40 | 0,40 | 0,04 | -0,05 | 0,58 |
| 0,60 | 0,80 | 0,77 | 0,54 | -0,04 |
| 1,00 | 0,90 | 0,01 | -0,26 | 0,00 |
| 0,40 | 0,40 | -0,08 | -0,23 | 0,12 |
| 0,80 | 0,80 | 0,08 | 0,00 | 0,23 |
| 0,00 | 0,10 | 0,02 | 0,00 | 0,00 |
| 1,00 | 0,90 | 0,69 | -0,17 | 0,65 |
| 1,00 | 0,90 | -0,16 | 0,10 | -0,26 |
| 0,30 | 0,20 | 0,00 | 0,00 | 0,11 |
| 0,20 | 0,20 | 0,22 | 0,02 | -0,01 |
| 1,00 | 0,90 | 0,07 | 0,40 | 0,05 |
| 0,90 | 0,80 | 0,60 | 0,08 | 0,37 |
| 0,70 | 0,70 | -0,17 | 0,40 | -0,52 |
| 0,10 | 0,20 | 0,00 | 0,16 | 0,02 |
| 0,30 | 0,60 | -0,16 | 0,78 | -0,07 |
| 0,80 | 0,90 | -0,18 | 0,23 | -0,42 |
| 1,00 | 1,00 | -0,03 | 0,13 | -0,10 |
| 1,00 | 1,00 | -0,27 | -0,05 | -0,08 |
| 1,00 | 1,00 | 0,17 | 0,00 | -0,01 |
| 1,00 | 1,00 | -0,02 | 0,21 | -0,01 |
| 1,00 | 1,00 | 0,04 | -0,09 | 0,21 |
| 0,40 | 0,40 | 0,26 | 0,50 | -0,61 |
| 0,80 | 0,50 | 0,24 | -0,63 | 1,08 |
| 0,60 | 0,70 | 0,51 | 0,19 | 0,21 |
| 0,10 | 0,00 | -0,23 | -0,01 | -0,49 |
| 1,00 | 1,00 | 0,20 | 0,00 | 0,15 |
| 1,00 | 0,80 | -0,09 | -0,63 | 0,15 |
| 0,60 | 0,80 | 0,60 | 0,63 | 0,24 |
| 0,50 | 0,40 | 0,83 | -0,15 | 0,07 |
| 0,70 | 0,80 | -0,27 | 0,34 | -0,70 |
| 1,00 | 0,90 | -0,08 | 0,49 | -0,14 |
| 0,60 | 0,80 | 0,28 | 0,48 | 0,12 |
| 0,70 | 0,70 | -0,25 | 0,09 | -0,33 |
| 1,00 | 1,00 | -0,05 | 0,39 | 0,04 |
| 0,90 | 0,90 | -0,21 | -0,20 | -0,27 |
| 0,90 | 0,90 | 0,06 | -0,22 | 0,37 |
| 1,00 | 0,70 | -0,74 | -0,74 | -0,01 |
| 0,90 | 0,80 | -0,32 | -0,63 | 0,01 |
| 0,80 | 0,60 | 0,70 | -0,28 | 0,49 |
| 0,60 | 0,60 | 0,61 | -0,10 | 0,42 |
| 0,00 | 0,20 | 1,57 | 0,37 | 0,00 |
| 0,10 | 0,20 | -0,01 | 0,26 | 0,01 |

| | | | | |
|---|---|---|---|---|
| 1,00 | 1,00 | -0,08 | 0,02 | 0,06 |
| 1,00 | 1,00 | -0,35 | -0,08 | 0,00 |
| 1,00 | 1,00 | -0,27 | 0,01 | -0,06 |
| 0,40 | 0,70 | 1,52 | 1,41 | -0,68 |
| 0,70 | 0,70 | 1,08 | -0,49 | 1,81 |
| 0,60 | 0,80 | 0,49 | 0,54 | 0,34 |
| 0,60 | 0,40 | -0,25 | -0,35 | -0,09 |
| 0,50 | 0,60 | -0,57 | 0,06 | -0,06 |
| 0,80 | 0,90 | -0,61 | 0,79 | -0,76 |
| 0,90 | 0,90 | -0,52 | 0,21 | -0,05 |
| 0,70 | 0,90 | -1,15 | 0,15 | -0,81 |
| 0,00 | 0,30 | 0,00 | 0,61 | 0,00 |
| 0,90 | 1,00 | 0,54 | 0,17 | 0,83 |
| 0,10 | 0,00 | 0,05 | 0,00 | -0,02 |
| 0,40 | 0,50 | -0,24 | -0,04 | -0,24 |
| 0,50 | 0,40 | -0,47 | 0,22 | -0,41 |
| 0,00 | 0,30 | 0,00 | 0,50 | 0,00 |
| 0,80 | 0,80 | 0,97 | 0,20 | 0,51 |
| 0,90 | 0,90 | -0,19 | 0,03 | 0,01 |
| 0,80 | 1,00 | 0,52 | 0,99 | 0,28 |
| 0,30 | 0,00 | -0,44 | -0,38 | -0,17 |
| 0,30 | 0,30 | 0,90 | -0,11 | -0,15 |
| 0,90 | 1,00 | -0,38 | 0,38 | -0,21 |
| 0,80 | 0,90 | 0,66 | 0,17 | 0,33 |
| 0,70 | 0,80 | 0,02 | 0,55 | -0,12 |
| 0,20 | 0,10 | 0,00 | -0,08 | 0,09 |
| 0,40 | 0,60 | -0,40 | 0,22 | -0,61 |
| 0,70 | 0,60 | 0,03 | -0,17 | 0,53 |
| 0,00 | 0,30 | 0,00 | 0,60 | 0,00 |
| 0,30 | 0,50 | 0,97 | 0,56 | -0,07 |
| 0,70 | 0,80 | -0,42 | 0,05 | -0,37 |
| 1,00 | 1,00 | 0,08 | 0,13 | 0,02 |
| 0,80 | 0,90 | 0,30 | 0,64 | -0,37 |
| 0,60 | 0,60 | -1,24 | -0,39 | -0,60 |
| 1,00 | 1,00 | 0,00 | -0,13 | 0,17 |
| 0,30 | 0,50 | -0,14 | 0,10 | -1,42 |
| 0,90 | 0,90 | 0,67 | -0,35 | 0,84 |
| 0,70 | 0,60 | -0,25 | -0,12 | -0,36 |
| 0,70 | 0,60 | -0,03 | -0,35 | -0,45 |
| 1,00 | 0,90 | -0,45 | 0,09 | 0,27 |
| 0,90 | 0,80 | 0,55 | -0,27 | 0,39 |
| 0,70 | 0,90 | 0,80 | 0,41 | 0,18 |
| 1,00 | 0,90 | 0,22 | -0,71 | 0,53 |
| 0,10 | 0,10 | 0,52 | 0,00 | -0,05 |
| 0,70 | 0,70 | -0,44 | 0,15 | -0,07 |
| 0,50 | 0,30 | -0,07 | -0,18 | 0,27 |

| | | | | |
|---|---|---|---|---|
| 0,50 | 0,80 | -0,44 | 1,31 | -1,07 |
| 0,20 | 0,40 | 0,18 | 0,37 | 0,14 |
| 0,80 | 0,80 | 0,24 | 0,14 | 0,37 |
| 0,60 | 0,50 | -0,61 | -0,04 | -0,01 |
| 0,40 | 0,70 | 1,09 | 1,44 | 0,85 |
| 0,50 | 0,40 | 0,01 | -0,09 | 0,68 |
| 0,30 | 0,20 | -0,08 | -0,25 | 0,21 |
| 0,70 | 0,70 | 0,08 | -0,18 | -0,24 |
| 0,70 | 0,50 | 0,05 | -0,52 | -0,48 |
| 0,30 | 0,40 | -0,30 | 0,74 | 0,12 |
| 0,90 | 0,90 | -0,09 | 0,08 | -0,16 |
| 1,00 | 1,00 | -0,09 | -0,16 | -0,04 |
| 1,00 | 1,00 | 0,01 | 0,21 | -0,02 |
| 1,00 | 1,00 | 0,12 | 0,21 | -0,03 |
| 1,00 | 1,00 | -0,14 | -0,06 | -0,03 |
| 1,00 | 1,00 | 0,01 | 0,13 | 0,08 |
| 1,00 | 1,00 | 0,11 | -0,01 | -0,05 |
| 0,80 | 0,40 | 0,12 | -0,62 | -0,07 |
| 0,40 | 0,40 | 0,42 | 0,15 | -0,02 |
| 0,90 | 0,90 | -0,11 | 0,07 | -0,21 |
| 1,00 | 1,00 | 0,26 | -0,16 | -0,03 |
| 0,80 | 0,80 | 0,52 | 0,33 | 0,01 |
| 0,10 | 0,30 | 0,01 | 0,05 | 0,00 |
| 0,00 | 0,10 | 0,18 | 0,01 | 0,00 |
| 0,30 | 0,50 | 0,42 | 0,21 | 0,07 |
| 0,00 | 0,10 | 0,36 | 0,02 | 0,00 |
| 0,40 | 0,60 | 0,28 | 0,74 | -0,29 |
| 0,90 | 0,90 | -0,18 | -0,06 | -0,07 |
| 1,00 | 1,00 | -0,20 | -0,43 | -0,29 |
| 0,20 | 0,40 | 1,45 | 0,68 | 0,23 |
| 0,70 | 0,70 | -0,03 | 0,33 | 0,61 |
| 0,90 | 0,50 | 0,10 | -0,78 | 0,02 |
| 0,80 | 0,90 | -0,20 | 0,06 | 0,18 |
| 0,00 | 0,20 | -0,01 | 0,44 | -0,05 |
| 1,00 | 1,00 | -0,21 | -0,27 | -0,14 |

| | P-value | | |
|---|---|---|---|
| FMD / SD (T2) | T2 / T1 (SD) | T2 / T1 (FMD) | FMD / SD (T1) |
| -1,00 | 9,15E-03 | 3,23E-02 | 3,36E-01 |
| -0,17 | 2,44E-02 | 8,05E-03 | 2,70E-01 |
| 0,47 | 1,78E-02 | 2,62E-01 | 7,75E-01 |
| 0,47 | 1,29E-02 | 2,62E-01 | 9,67E-01 |
| 0,57 | 3,30E-02 | 6,83E-01 | 7,13E-01 |
| -0,27 | 9,15E-03 | 3,23E-02 | 6,53E-01 |
| 0,42 | 9,06E-01 | 1,26E-01 | 8,38E-01 |
| 0,53 | 2,36E-01 | 4,15E-01 | 4,88E-01 |
| 1,01 | 7,59E-02 | 1,00E+00 | 9,67E-01 |
| -0,91 | 1,55E-01 | 5,41E-01 | 9,02E-01 |
| -0,04 | 1,78E-02 | 5,41E-01 | 8,16E-02 |
| 1,67 | 5,87E-02 | 3,08E-01 | 9,67E-01 |
| 0,16 | 1,78E-02 | 2,21E-01 | 1,53E-01 |
| 0,13 | 1,55E-01 | 4,15E-02 | 5,96E-01 |
| 0,36 | 4,40E-02 | 2,21E-01 | 2,36E-01 |
| 0,36 | 9,06E-01 | 4,15E-01 | 1,31E-01 |
| 1,16 | 1,55E-01 | 2,62E-01 | 5,50E-02 |
| -0,45 | 6,36E-01 | 2,49E-02 | 4,38E-01 |
| 1,97 | NA | 5,91E-02 | NA |
| 0,79 | 8,13E-01 | 1,44E-02 | 7,44E-01 |
| 0,22 | 4,77E-01 | 6,83E-01 | 3,07E-01 |
| -0,33 | 5,54E-01 | 1,00E+00 | 1,01E-02 |
| 1,01 | 2,01E-01 | 5,54E-01 | 5,08E-01 |
| -0,73 | 1,00E+00 | 1,81E-01 | 5,71E-01 |
| -0,23 | 6,24E-01 | 5,53E-01 | 4,17E-02 |
| -1,15 | 7,59E-02 | 6,73E-01 | 6,17E-01 |
| -1,51 | 2,34E-01 | 7,26E-01 | 1,28E-01 |
| -1,18 | 6,36E-01 | 6,73E-01 | 2,70E-01 |
| 3,11 | NA | 5,19E-02 | 9,47E-02 |
| -0,16 | 3,60E-02 | 5,91E-02 | 1,00E+00 |
| 0,37 | 4,40E-02 | 8,38E-01 | 2,36E-01 |
| -0,80 | 1,78E-01 | 7,87E-01 | 1,00E+00 |
| -0,23 | 8,13E-01 | 6,10E-01 | 3,07E-01 |
| 0,46 | 1,00E+00 | 7,26E-01 | 8,33E-02 |
| 0,71 | 4,23E-01 | 6,24E-01 | 2,56E-01 |
| -0,12 | 9,33E-01 | 4,41E-01 | 9,02E-01 |
| -0,04 | 9,06E-01 | 6,10E-01 | 1,31E-01 |
| 0,22 | 1,00E+00 | 3,71E-01 | 5,63E-01 |
| -0,07 | 1,00E+00 | 3,71E-01 | 6,24E-02 |
| 0,01 | 7,22E-01 | 1,00E+00 | 2,06E-01 |
| 0,31 | 5,54E-01 | 1,85E-01 | 1,00E+00 |
| 0,82 | NA | 1,81E-01 | 3,99E-01 |
| -0,79 | 1,07E-01 | 6,24E-01 | 8,34E-01 |
| -0,66 | 5,92E-02 | 3,61E-01 | 6,07E-01 |

| | | | |
|---|---|---|---|
| 0,16 | 1,93E-01 | 4,76E-01 | 4,38E-01 |
| -0,82 | 7,22E-01 | 4,76E-01 | 9,35E-01 |
| 0,10 | 4,77E-01 | 4,76E-01 | 2,70E-01 |
| -0,22 | 6,24E-01 | 7,60E-01 | 4,31E-01 |
| 0,49 | 1,00E+00 | 6,75E-01 | 2,99E-01 |
| -0,28 | 2,94E-01 | 1,00E+00 | 9,66E-01 |
| -0,27 | 6,36E-01 | 3,59E-01 | 7,75E-01 |
| -0,04 | 6,74E-01 | 8,00E-01 | 8,95E-01 |
| 0,15 | 9,44E-01 | 2,62E-01 | 7,13E-01 |
| -0,02 | 3,71E-01 | 1,00E+00 | NA |
| -0,21 | 3,43E-01 | 7,60E-01 | 2,06E-01 |
| 0,00 | 4,77E-01 | 6,10E-01 | 2,36E-01 |
| 0,11 | NA | 7,89E-01 | 9,47E-02 |
| -0,22 | 7,87E-01 | 5,84E-01 | 9,09E-01 |
| 0,38 | 7,22E-01 | 1,85E-01 | 7,75E-01 |
| -0,15 | 1,83E-01 | 1,00E+00 | 4,87E-01 |
| 0,06 | 8,34E-01 | 5,41E-01 | 5,37E-01 |
| 0,18 | NA | 1,00E+00 | 3,99E-01 |
| 0,87 | 7,89E-01 | 1,41E-01 | 8,82E-01 |
| -0,01 | 9,06E-01 | 7,60E-01 | 7,75E-01 |
| 0,07 | 7,22E-01 | 6,83E-01 | 5,40E-01 |
| 0,14 | 1,24E-01 | 4,76E-01 | 2,70E-01 |
| -0,18 | 6,36E-01 | 1,00E+00 | 1,00E+00 |
| 0,22 | 7,22E-01 | 2,21E-01 | 9,67E-01 |
| 0,07 | 4,77E-01 | 8,38E-01 | 9,42E-02 |
| -0,36 | 2,72E-01 | 2,05E-01 | 2,81E-01 |
| 0,21 | 1,00E+00 | 5,54E-01 | 7,18E-02 |
| -0,12 | 7,59E-02 | 8,34E-01 | 3,00E-01 |
| -0,27 | 5,90E-01 | 1,00E+00 | 1,71E-01 |
| -0,04 | 1,93E-01 | 6,83E-01 | 7,13E-01 |
| -0,39 | 8,13E-01 | 2,62E-01 | 5,96E-01 |
| 0,27 | 5,29E-01 | 1,55E-01 | 7,63E-01 |
| -0,91 | 3,63E-01 | 1,00E+00 | 9,66E-01 |
| -0,09 | 1,00E+00 | 3,43E-01 | 2,70E-01 |
| 0,43 | 8,13E-01 | 8,31E-02 | 1,00E+00 |
| 0,32 | 2,95E-01 | 5,41E-01 | 7,67E-01 |
| 0,01 | 7,26E-01 | 6,10E-01 | 4,34E-01 |
| 0,48 | 9,06E-01 | 8,31E-02 | 8,38E-01 |
| -0,27 | 6,36E-01 | 4,76E-01 | 1,31E-01 |
| 0,09 | 4,77E-01 | 4,15E-01 | 1,31E-01 |
| -0,01 | 3,63E-01 | 2,21E-01 | 7,75E-01 |
| -0,30 | 7,56E-02 | 4,76E-01 | 7,13E-01 |
| -0,49 | 2,34E-01 | 1,00E+00 | 3,03E-01 |
| -0,28 | 2,72E-01 | 1,00E+00 | 3,79E-01 |
| -1,20 | 1,00E-01 | 3,71E-01 | NA |
| 0,27 | 1,00E+00 | 3,71E-01 | 1,00E+00 |

| | | | |
|---|---|---|---|
| 0,16 | 5,54E-01 | 5,41E-01 | 4,88E-01 |
| 0,26 | 2,36E-01 | 4,76E-01 | 9,03E-01 |
| 0,22 | 4,77E-01 | 7,60E-01 | 8,38E-01 |
| -0,80 | 5,19E-02 | 2,05E-01 | 4,53E-01 |
| 0,24 | 4,02E-01 | 1,07E-01 | 6,37E-02 |
| 0,38 | 1,00E+00 | 2,36E-01 | 6,35E-01 |
| -0,19 | 9,33E-01 | 9,33E-01 | 7,69E-01 |
| 0,57 | 5,91E-02 | 1,00E+00 | 9,66E-01 |
| 0,64 | 9,72E-02 | 3,08E-01 | 1,53E-01 |
| 0,68 | 1,55E-01 | 6,10E-01 | 8,70E-01 |
| 0,50 | 6,36E-01 | 8,38E-01 | 3,47E-01 |
| 0,61 | NA | 1,81E-01 | NA |
| 0,46 | 4,07E-01 | 1,03E-01 | 2,06E-01 |
| -0,08 | 7,89E-01 | 1,00E+00 | 9,39E-01 |
| -0,04 | 6,36E-01 | 7,67E-01 | 4,12E-01 |
| 0,28 | 6,24E-01 | 4,47E-01 | 2,58E-01 |
| 0,50 | NA | 1,81E-01 | NA |
| -0,25 | 5,80E-02 | 3,43E-01 | 1,76E-01 |
| 0,22 | 4,77E-01 | 4,76E-01 | 9,03E-01 |
| 0,75 | 1,51E-01 | 6,65E-02 | 7,11E-01 |
| -0,11 | 2,08E-01 | 1,81E-01 | 6,37E-01 |
| -1,16 | 1,07E-01 | 7,89E-01 | 7,77E-01 |
| 0,55 | 1,24E-01 | 1,26E-01 | 5,40E-01 |
| -0,16 | 5,54E-01 | 8,38E-01 | 3,90E-01 |
| 0,41 | 5,29E-01 | 1,93E-01 | 9,35E-01 |
| 0,01 | NA | 1,00E+00 | 1,93E-01 |
| 0,01 | 7,26E-01 | 6,73E-01 | 1,94E-01 |
| 0,33 | 9,33E-01 | 9,06E-01 | 5,86E-01 |
| 0,60 | NA | 1,81E-01 | NA |
| -0,49 | 2,34E-01 | 6,73E-01 | 1,00E+00 |
| 0,10 | 2,36E-01 | 2,34E-01 | 5,66E-01 |
| 0,08 | 7,22E-01 | 2,62E-01 | 1,00E+00 |
| -0,03 | 8,13E-01 | 2,62E-01 | 4,87E-01 |
| 0,25 | 2,36E-01 | 1,83E-01 | 3,43E-01 |
| 0,04 | 9,06E-01 | 4,76E-01 | 5,96E-01 |
| -1,18 | 7,26E-01 | 9,33E-01 | 2,16E-01 |
| -0,18 | 3,43E-01 | 4,07E-01 | 1,21E-01 |
| -0,23 | 1,51E-01 | 1,00E+00 | 3,62E-01 |
| -0,76 | 9,44E-01 | 1,00E+00 | 6,51E-01 |
| 0,81 | 2,86E-01 | 6,83E-01 | 4,38E-01 |
| -0,43 | 7,56E-02 | 3,59E-01 | 1,78E-01 |
| -0,20 | 1,07E-01 | 1,00E+00 | 7,11E-01 |
| -0,40 | 9,06E-01 | 1,85E-01 | 3,91E-01 |
| -0,57 | 2,81E-01 | 1,00E+00 | 6,07E-01 |
| 0,53 | 1,07E-01 | 9,44E-01 | 5,86E-01 |
| 0,16 | 4,23E-01 | 6,75E-01 | 3,22E-01 |

| | | | |
|---|---|---|---|
| 0,68 | 7,22E-01 | 8,01E-02 | 1,74E-01 |
| 0,33 | 1,00E+00 | 5,29E-01 | 6,53E-01 |
| 0,27 | 8,34E-01 | 9,19E-01 | 6,51E-01 |
| 0,56 | 3,61E-01 | 5,29E-01 | 8,29E-01 |
| 1,20 | 2,01E-01 | 5,80E-02 | 1,55E-01 |
| 0,58 | 8,55E-01 | 8,34E-01 | 1,87E-01 |
| 0,04 | 7,89E-01 | 2,01E-01 | 5,27E-01 |
| -0,51 | 2,86E-01 | 9,06E-01 | 1,00E+00 |
| -1,05 | 1,00E+00 | 5,29E-01 | 6,82E-01 |
| 1,15 | 3,71E-01 | 3,53E-01 | 8,33E-01 |
| 0,01 | 6,24E-01 | 2,21E-01 | 5,13E-01 |
| -0,11 | 1,00E+00 | 1,26E-01 | 9,03E-01 |
| 0,19 | 7,22E-01 | 1,85E-01 | 9,03E-01 |
| 0,06 | 2,36E-01 | 1,03E-01 | 7,75E-01 |
| 0,05 | 6,36E-01 | 5,41E-01 | 8,38E-01 |
| 0,19 | 8,13E-01 | 2,62E-01 | 5,96E-01 |
| -0,17 | 4,07E-01 | 6,83E-01 | 7,75E-01 |
| -0,81 | 1,00E+00 | 2,86E-01 | 7,74E-01 |
| -0,29 | 4,47E-01 | 4,47E-01 | 8,92E-01 |
| -0,02 | 2,86E-01 | 1,00E+00 | 7,75E-01 |
| -0,44 | 4,07E-01 | 8,38E-01 | 7,75E-01 |
| -0,19 | 1,93E-01 | 4,07E-01 | 7,75E-01 |
| 0,05 | 1,00E+00 | 7,89E-01 | 3,99E-01 |
| -0,18 | 1,81E-01 | 1,00E+00 | NA |
| -0,13 | 5,90E-01 | 6,73E-01 | 8,33E-01 |
| -0,33 | 3,71E-01 | 1,00E+00 | NA |
| 0,16 | 5,54E-01 | 8,01E-02 | 4,02E-01 |
| 0,05 | 6,36E-01 | 8,38E-01 | 7,13E-01 |
| -0,52 | 1,00E+00 | 6,65E-02 | 6,62E-02 |
| -0,54 | 1,06E-01 | 1,78E-01 | 5,63E-01 |
| 0,98 | 1,00E+00 | 4,41E-01 | 2,81E-01 |
| -0,86 | 4,07E-01 | 7,22E-01 | 1,00E+00 |
| 0,44 | 3,43E-01 | 1,85E-01 | 4,87E-01 |
| 0,41 | 1,00E+00 | 3,71E-01 | 3,43E-01 |
| -0,20 | 5,54E-01 | 8,31E-02 | 3,48E-01 |

| | Adjusted p-value | | |
|---|---|---|---|
| FMD / SD (T2) | T2 / T1 (SD) | T2 / T1 (FMD) | FMD / SD (T1) |
| 1,46E-02 | 4,81E-01 | 7,87E-01 | 1,00E+00 |
| 4,22E-01 | 5,64E-01 | 7,87E-01 | 1,00E+00 |
| 1,35E-02 | 4,81E-01 | 7,87E-01 | 1,00E+00 |
| 2,46E-02 | 4,81E-01 | 7,87E-01 | 1,00E+00 |
| 2,59E-03 | 5,86E-01 | 9,58E-01 | 1,00E+00 |
| 1,72E-01 | 4,81E-01 | 7,87E-01 | 1,00E+00 |
| 3,65E-02 | 1,00E+00 | 7,87E-01 | 1,00E+00 |
| 2,94E-02 | 7,22E-01 | 9,14E-01 | 1,00E+00 |
| 9,22E-03 | 5,86E-01 | 1,00E+00 | 1,00E+00 |
| 4,14E-03 | 6,98E-01 | 9,38E-01 | 1,00E+00 |
| 9,12E-01 | 4,81E-01 | 9,38E-01 | 1,00E+00 |
| 3,79E-02 | 5,86E-01 | 8,78E-01 | 1,00E+00 |
| 2,58E-01 | 4,81E-01 | 7,87E-01 | 1,00E+00 |
| 3,19E-01 | 6,98E-01 | 7,87E-01 | 1,00E+00 |
| 6,93E-02 | 5,86E-01 | 7,87E-01 | 1,00E+00 |
| 1,13E-02 | 1,00E+00 | 9,14E-01 | 1,00E+00 |
| 4,55E-03 | 6,98E-01 | 7,87E-01 | 1,00E+00 |
| 3,02E-01 | 9,53E-01 | 7,87E-01 | 1,00E+00 |
| 1,65E-02 | NA | 7,87E-01 | NA |
| 9,48E-02 | 1,00E+00 | 7,87E-01 | 1,00E+00 |
| 4,32E-02 | 9,09E-01 | 9,58E-01 | 1,00E+00 |
| 4,69E-01 | 9,53E-01 | 1,00E+00 | 1,00E+00 |
| 3,17E-02 | 7,22E-01 | 9,38E-01 | 1,00E+00 |
| 1,87E-02 | 1,00E+00 | 7,87E-01 | 1,00E+00 |
| 6,40E-01 | 9,53E-01 | 9,38E-01 | 1,00E+00 |
| 2,21E-02 | 5,86E-01 | 9,58E-01 | 1,00E+00 |
| 1,04E-02 | 7,22E-01 | 9,94E-01 | 1,00E+00 |
| 4,80E-02 | 9,53E-01 | 9,58E-01 | 1,00E+00 |
| 9,50E-04 | NA | 7,87E-01 | 1,00E+00 |
| 2,81E-01 | 5,86E-01 | 7,87E-01 | 1,00E+00 |
| 1,30E-01 | 5,86E-01 | 1,00E+00 | 1,00E+00 |
| 6,17E-02 | 7,22E-01 | 1,00E+00 | 1,00E+00 |
| 4,13E-01 | 1,00E+00 | 9,58E-01 | 1,00E+00 |
| 3,20E-01 | 1,00E+00 | 9,94E-01 | 1,00E+00 |
| 3,09E-01 | 8,89E-01 | 9,58E-01 | 1,00E+00 |
| 8,29E-01 | 1,00E+00 | 9,14E-01 | 1,00E+00 |
| 7,74E-01 | 1,00E+00 | 9,58E-01 | 1,00E+00 |
| 5,41E-01 | 1,00E+00 | 8,94E-01 | 1,00E+00 |
| 7,25E-01 | 1,00E+00 | 8,94E-01 | 1,00E+00 |
| 5,46E-01 | 9,80E-01 | 1,00E+00 | 1,00E+00 |
| 1,12E-01 | 9,53E-01 | 7,87E-01 | 1,00E+00 |
| 9,30E-02 | NA | 7,87E-01 | 1,00E+00 |
| 2,26E-01 | 6,21E-01 | 9,58E-01 | 1,00E+00 |
| 2,66E-01 | 5,86E-01 | 8,94E-01 | 1,00E+00 |

| | | | |
|---|---|---|---|
| 5,37E-01 | 7,22E-01 | 9,14E-01 | 1,00E+00 |
| 1,87E-01 | 9,80E-01 | 9,14E-01 | 1,00E+00 |
| 8,63E-01 | 9,09E-01 | 9,14E-01 | 1,00E+00 |
| 5,76E-01 | 9,53E-01 | 1,00E+00 | 1,00E+00 |
| 3,36E-01 | 1,00E+00 | 9,58E-01 | 1,00E+00 |
| 3,60E-01 | 7,82E-01 | 1,00E+00 | 1,00E+00 |
| 2,13E-01 | 9,53E-01 | 8,94E-01 | 1,00E+00 |
| 6,22E-01 | 9,80E-01 | 1,00E+00 | 1,00E+00 |
| 7,34E-01 | 1,00E+00 | 7,87E-01 | 1,00E+00 |
| 6,30E-01 | 8,59E-01 | 1,00E+00 | NA |
| 5,84E-01 | 8,59E-01 | 1,00E+00 | 1,00E+00 |
| 8,35E-01 | 9,09E-01 | 9,58E-01 | 1,00E+00 |
| 1,51E-01 | NA | 1,00E+00 | 1,00E+00 |
| 2,05E-01 | 1,00E+00 | 9,58E-01 | 1,00E+00 |
| 4,33E-01 | 9,80E-01 | 7,87E-01 | 1,00E+00 |
| 6,73E-01 | 7,22E-01 | 1,00E+00 | 1,00E+00 |
| 9,48E-01 | 1,00E+00 | 9,38E-01 | 1,00E+00 |
| 1,94E-01 | NA | 1,00E+00 | 1,00E+00 |
| 1,44E-01 | 1,00E+00 | 7,87E-01 | 1,00E+00 |
| 4,65E-01 | 1,00E+00 | 1,00E+00 | 1,00E+00 |
| 7,96E-01 | 9,80E-01 | 9,58E-01 | 1,00E+00 |
| 1,82E-01 | 6,67E-01 | 9,14E-01 | 1,00E+00 |
| 6,35E-01 | 9,53E-01 | 1,00E+00 | 1,00E+00 |
| 1,94E-01 | 9,80E-01 | 7,87E-01 | 1,00E+00 |
| 5,58E-01 | 9,09E-01 | 1,00E+00 | 1,00E+00 |
| 7,41E-01 | 7,82E-01 | 7,87E-01 | 1,00E+00 |
| 7,40E-01 | 1,00E+00 | 9,38E-01 | 1,00E+00 |
| 6,95E-01 | 5,86E-01 | 1,00E+00 | 1,00E+00 |
| 8,23E-02 | 9,53E-01 | 1,00E+00 | 1,00E+00 |
| 5,76E-01 | 7,22E-01 | 9,58E-01 | 1,00E+00 |
| 4,66E-01 | 1,00E+00 | 7,87E-01 | 1,00E+00 |
| 5,86E-01 | 9,53E-01 | 7,87E-01 | 1,00E+00 |
| 2,06E-01 | 8,59E-01 | 1,00E+00 | 1,00E+00 |
| 8,23E-01 | 1,00E+00 | 8,94E-01 | 1,00E+00 |
| 6,66E-01 | 1,00E+00 | 7,87E-01 | 1,00E+00 |
| 5,27E-01 | 7,82E-01 | 9,38E-01 | 1,00E+00 |
| 9,60E-01 | 9,80E-01 | 9,58E-01 | 1,00E+00 |
| 1,02E-01 | 1,00E+00 | 7,87E-01 | 1,00E+00 |
| 1,42E-01 | 9,53E-01 | 9,14E-01 | 1,00E+00 |
| 7,33E-01 | 9,09E-01 | 9,14E-01 | 1,00E+00 |
| 1,00E+00 | 8,59E-01 | 7,87E-01 | 1,00E+00 |
| 2,98E-01 | 5,86E-01 | 9,14E-01 | 1,00E+00 |
| 2,73E-01 | 7,22E-01 | 1,00E+00 | 1,00E+00 |
| 8,61E-01 | 7,82E-01 | 1,00E+00 | 1,00E+00 |
| 1,95E-01 | 6,21E-01 | 8,94E-01 | NA |
| 1,81E-01 | 1,00E+00 | 8,94E-01 | 1,00E+00 |

| | | | |
|---|---|---|---|
| 9,73E-02 | 9,53E-01 | 9,38E-01 | 1,00E+00 |
| 9,71E-02 | 7,22E-01 | 9,14E-01 | 1,00E+00 |
| 1,67E-01 | 9,09E-01 | 1,00E+00 | 1,00E+00 |
| 4,37E-01 | 5,86E-01 | 7,87E-01 | 1,00E+00 |
| 7,75E-01 | 8,79E-01 | 7,87E-01 | 1,00E+00 |
| 6,06E-01 | 1,00E+00 | 7,87E-01 | 1,00E+00 |
| 7,52E-01 | 1,00E+00 | 1,00E+00 | 1,00E+00 |
| 1,44E-01 | 5,86E-01 | 1,00E+00 | 1,00E+00 |
| 2,35E-01 | 6,21E-01 | 8,78E-01 | 1,00E+00 |
| 2,39E-01 | 6,98E-01 | 9,58E-01 | 1,00E+00 |
| 6,22E-01 | 9,53E-01 | 1,00E+00 | 1,00E+00 |
| 6,88E-02 | NA | 7,87E-01 | NA |
| 4,76E-01 | 8,79E-01 | 7,87E-01 | 1,00E+00 |
| 2,03E-01 | 1,00E+00 | 1,00E+00 | 1,00E+00 |
| 7,43E-01 | 9,53E-01 | 1,00E+00 | 1,00E+00 |
| 5,54E-01 | 9,53E-01 | 9,14E-01 | 1,00E+00 |
| 9,83E-02 | NA | 7,87E-01 | NA |
| 4,76E-01 | 5,86E-01 | 8,94E-01 | 1,00E+00 |
| 8,91E-01 | 9,09E-01 | 9,14E-01 | 1,00E+00 |
| 9,82E-02 | 6,98E-01 | 7,87E-01 | 1,00E+00 |
| 1,24E-01 | 7,22E-01 | 7,87E-01 | 1,00E+00 |
| 7,65E-02 | 6,21E-01 | 1,00E+00 | 1,00E+00 |
| 1,43E-01 | 6,67E-01 | 7,87E-01 | 1,00E+00 |
| 8,08E-01 | 9,53E-01 | 1,00E+00 | 1,00E+00 |
| 3,39E-01 | 9,53E-01 | 7,87E-01 | 1,00E+00 |
| 3,58E-01 | NA | 1,00E+00 | 1,00E+00 |
| 9,53E-01 | 9,80E-01 | 9,58E-01 | 1,00E+00 |
| 5,22E-01 | 1,00E+00 | 1,00E+00 | 1,00E+00 |
| 8,59E-02 | NA | 7,87E-01 | NA |
| 4,74E-01 | 7,22E-01 | 9,58E-01 | 1,00E+00 |
| 9,38E-01 | 7,22E-01 | 7,87E-01 | 1,00E+00 |
| 6,28E-01 | 9,80E-01 | 7,87E-01 | 1,00E+00 |
| 5,01E-01 | 1,00E+00 | 7,87E-01 | 1,00E+00 |
| 8,03E-01 | 7,22E-01 | 7,87E-01 | 1,00E+00 |
| 8,11E-01 | 1,00E+00 | 9,14E-01 | 1,00E+00 |
| 2,04E-01 | 9,80E-01 | 1,00E+00 | 1,00E+00 |
| 5,69E-01 | 8,59E-01 | 9,14E-01 | 1,00E+00 |
| 8,23E-01 | 6,98E-01 | 1,00E+00 | 1,00E+00 |
| 3,83E-01 | 1,00E+00 | 1,00E+00 | 1,00E+00 |
| 9,90E-02 | 7,82E-01 | 9,58E-01 | 1,00E+00 |
| 2,63E-01 | 5,86E-01 | 8,94E-01 | 1,00E+00 |
| 7,99E-01 | 6,21E-01 | 1,00E+00 | 1,00E+00 |
| 2,72E-01 | 1,00E+00 | 7,87E-01 | 1,00E+00 |
| 2,45E-01 | 7,82E-01 | 1,00E+00 | 1,00E+00 |
| 3,82E-01 | 6,21E-01 | 1,00E+00 | 1,00E+00 |
| 2,47E-01 | 8,89E-01 | 9,58E-01 | 1,00E+00 |

| | | | |
|---|---|---|---|
| 4,27E-01 | 9,80E-01 | 7,87E-01 | 1,00E+00 |
| 4,82E-01 | 1,00E+00 | 9,38E-01 | 1,00E+00 |
| 5,77E-01 | 1,00E+00 | 1,00E+00 | 1,00E+00 |
| 5,19E-01 | 8,59E-01 | 9,38E-01 | 1,00E+00 |
| 3,68E-01 | 7,22E-01 | 7,87E-01 | 1,00E+00 |
| 3,17E-01 | 1,00E+00 | 1,00E+00 | 1,00E+00 |
| 6,21E-01 | 1,00E+00 | 7,87E-01 | 1,00E+00 |
| 4,05E-01 | 7,82E-01 | 1,00E+00 | 1,00E+00 |
| 9,09E-02 | 1,00E+00 | 9,38E-01 | 1,00E+00 |
| 1,35E-01 | 8,59E-01 | 8,94E-01 | 1,00E+00 |
| 9,91E-01 | 9,53E-01 | 7,87E-01 | 1,00E+00 |
| 5,19E-01 | 1,00E+00 | 7,87E-01 | 1,00E+00 |
| 1,50E-01 | 9,80E-01 | 7,87E-01 | 1,00E+00 |
| 5,71E-01 | 7,22E-01 | 7,87E-01 | 1,00E+00 |
| 9,93E-01 | 9,53E-01 | 9,38E-01 | 1,00E+00 |
| 2,52E-01 | 1,00E+00 | 7,87E-01 | 1,00E+00 |
| 5,19E-01 | 8,79E-01 | 9,58E-01 | 1,00E+00 |
| 6,29E-02 | 1,00E+00 | 8,44E-01 | 1,00E+00 |
| 5,02E-01 | 9,09E-01 | 9,14E-01 | 1,00E+00 |
| 5,78E-01 | 7,82E-01 | 1,00E+00 | 1,00E+00 |
| 9,39E-02 | 8,79E-01 | 1,00E+00 | 1,00E+00 |
| 5,24E-01 | 7,22E-01 | 9,14E-01 | 1,00E+00 |
| 7,59E-01 | 1,00E+00 | 1,00E+00 | 1,00E+00 |
| 2,99E-01 | 7,22E-01 | 1,00E+00 | NA |
| 7,67E-01 | 9,53E-01 | 9,58E-01 | 1,00E+00 |
| 6,55E-01 | 8,59E-01 | 1,00E+00 | NA |
| 9,05E-01 | 9,53E-01 | 7,87E-01 | 1,00E+00 |
| 9,09E-01 | 9,53E-01 | 1,00E+00 | 1,00E+00 |
| 7,44E-02 | 1,00E+00 | 7,87E-01 | 1,00E+00 |
| 6,29E-01 | 6,21E-01 | 7,87E-01 | 1,00E+00 |
| 5,58E-02 | 1,00E+00 | 9,14E-01 | 1,00E+00 |
| 1,50E-01 | 8,79E-01 | 9,94E-01 | 1,00E+00 |
| 5,49E-01 | 8,59E-01 | 7,87E-01 | 1,00E+00 |
| 4,47E-01 | 1,00E+00 | 8,94E-01 | 1,00E+00 |
| 5,08E-01 | 9,53E-01 | 7,87E-01 | 1,00E+00 |

| | Prevalence shift | | |
|---|---|---|---|
| FMD / SD (T2) | T2 / T1 (SD) | T2 / T1 (FMD) | FMD / SD (T1) |
| 2,78E-01 | 0,22 | 0,50 | -0,28 |
| 7,93E-01 | 0,00 | 0,00 | 0,00 |
| 2,78E-01 | 0,00 | 0,00 | 0,00 |
| 3,24E-01 | 0,00 | 0,00 | 0,00 |
| 1,94E-01 | 0,00 | 0,00 | 0,00 |
| 5,76E-01 | 0,00 | 0,00 | 0,00 |
| 3,81E-01 | 0,00 | 0,00 | 0,00 |
| 3,60E-01 | -0,11 | 0,00 | 0,00 |
| 2,77E-01 | -0,67 | 0,10 | -0,18 |
| 1,94E-01 | 0,22 | 0,00 | 0,02 |
| 9,51E-01 | 0,44 | 0,00 | 0,46 |
| 3,81E-01 | -0,44 | -0,10 | -0,08 |
| 6,40E-01 | 0,00 | 0,10 | -0,10 |
| 6,75E-01 | 0,00 | 0,00 | 0,00 |
| 4,57E-01 | 0,00 | 0,00 | 0,00 |
| 2,77E-01 | 0,00 | 0,00 | 0,00 |
| 1,94E-01 | -0,22 | 0,10 | -0,10 |
| 6,70E-01 | 0,22 | -0,30 | 0,22 |
| 2,82E-01 | 0,00 | 0,50 | 0,00 |
| 4,57E-01 | -0,11 | 0,00 | -0,10 |
| 4,11E-01 | 0,00 | 0,00 | 0,00 |
| 8,01E-01 | -0,11 | -0,10 | 0,00 |
| 3,62E-01 | -0,22 | -0,10 | 0,27 |
| 2,91E-01 | 0,22 | -0,20 | -0,14 |
| 8,17E-01 | -0,44 | 0,00 | -0,59 |
| 3,15E-01 | 0,22 | 0,20 | -0,26 |
| 2,77E-01 | 0,11 | -0,10 | -0,18 |
| 4,32E-01 | -0,22 | -0,30 | -0,30 |
| 1,62E-01 | 0,00 | 0,40 | 0,30 |
| 6,48E-01 | -0,33 | -0,40 | -0,06 |
| 5,26E-01 | -0,22 | 0,00 | 0,11 |
| 4,57E-01 | 0,33 | 0,10 | -0,02 |
| 7,85E-01 | 0,11 | -0,20 | 0,11 |
| 6,75E-01 | 0,00 | -0,10 | 0,39 |
| 6,75E-01 | -0,11 | 0,10 | 0,17 |
| 9,03E-01 | 0,00 | -0,20 | 0,13 |
| 8,90E-01 | 0,11 | -0,10 | 0,11 |
| 8,01E-01 | 0,00 | 0,00 | 0,09 |
| 8,82E-01 | -0,11 | 0,20 | -0,33 |
| 8,01E-01 | 0,00 | 0,00 | 0,00 |
| 4,93E-01 | -0,22 | 0,00 | 0,00 |
| 4,57E-01 | 0,00 | 0,20 | 0,10 |
| 6,14E-01 | 0,00 | -0,10 | -0,07 |
| 6,40E-01 | 0,44 | 0,10 | -0,01 |

| | | | |
|---|---|---|---|
| 8,01E-01 | 0,11 | 0,00 | 0,11 |
| 5,78E-01 | 0,00 | -0,40 | 0,01 |
| 9,23E-01 | -0,11 | -0,30 | 0,22 |
| 8,01E-01 | -0,11 | 0,00 | -0,18 |
| 6,98E-01 | 0,00 | 0,00 | 0,18 |
| 7,25E-01 | 0,44 | 0,20 | 0,16 |
| 5,88E-01 | 0,00 | -0,10 | 0,00 |
| 8,16E-01 | -0,22 | 0,00 | -0,04 |
| 8,82E-01 | 0,00 | -0,10 | 0,02 |
| 8,16E-01 | 0,11 | 0,10 | 0,00 |
| 8,01E-01 | 0,11 | -0,10 | 0,11 |
| 9,04E-01 | 0,00 | 0,00 | -0,10 |
| 5,26E-01 | 0,00 | 0,00 | 0,20 |
| 5,78E-01 | 0,22 | 0,20 | -0,11 |
| 7,95E-01 | -0,11 | -0,10 | 0,00 |
| 8,41E-01 | 0,11 | -0,10 | 0,12 |
| 9,75E-01 | 0,00 | 0,00 | -0,08 |
| 5,78E-01 | 0,00 | 0,10 | 0,10 |
| 5,26E-01 | -0,11 | 0,30 | -0,03 |
| 8,01E-01 | 0,00 | 0,10 | -0,20 |
| 9,00E-01 | 0,00 | 0,00 | 0,00 |
| 5,78E-01 | 0,00 | 0,00 | 0,00 |
| 8,16E-01 | 0,00 | 0,00 | 0,00 |
| 5,78E-01 | 0,00 | 0,00 | 0,00 |
| 8,01E-01 | 0,00 | 0,00 | 0,00 |
| 8,82E-01 | -0,11 | 0,00 | -0,27 |
| 8,82E-01 | 0,00 | -0,30 | 0,36 |
| 8,61E-01 | 0,33 | 0,20 | 0,07 |
| 4,57E-01 | -0,11 | -0,10 | -0,34 |
| 8,01E-01 | 0,00 | 0,00 | 0,00 |
| 8,01E-01 | 0,00 | -0,20 | 0,11 |
| 8,01E-01 | 0,44 | 0,20 | 0,16 |
| 5,78E-01 | 0,22 | -0,10 | -0,06 |
| 9,02E-01 | -0,22 | 0,10 | -0,30 |
| 8,37E-01 | 0,00 | -0,10 | 0,00 |
| 8,01E-01 | 0,00 | 0,20 | 0,06 |
| 9,77E-01 | -0,11 | 0,00 | -0,08 |
| 4,59E-01 | 0,00 | 0,00 | 0,11 |
| 5,26E-01 | 0,00 | 0,00 | -0,10 |
| 8,82E-01 | 0,00 | 0,00 | -0,10 |
| 1,00E+00 | 0,00 | -0,30 | 0,11 |
| 6,70E-01 | 0,00 | -0,10 | -0,10 |
| 6,40E-01 | 0,33 | -0,20 | 0,24 |
| 9,23E-01 | 0,00 | 0,00 | -0,07 |
| 5,78E-01 | 0,44 | 0,20 | 0,00 |
| 5,78E-01 | -0,11 | 0,10 | -0,01 |

| | | | |
|---|---|---|---|
| 4,57E-01 | 0,00 | 0,00 | 0,00 |
| 4,57E-01 | 0,00 | 0,00 | 0,00 |
| 5,71E-01 | 0,00 | 0,00 | 0,00 |
| 7,95E-01 | 0,11 | 0,40 | -0,26 |
| 8,90E-01 | 0,44 | 0,00 | 0,48 |
| 8,16E-01 | 0,22 | 0,20 | 0,16 |
| 8,87E-01 | -0,22 | -0,20 | -0,07 |
| 5,26E-01 | -0,22 | 0,10 | -0,06 |
| 6,28E-01 | -0,11 | 0,10 | -0,20 |
| 6,28E-01 | -0,11 | 0,00 | 0,01 |
| 8,16E-01 | -0,33 | 0,00 | -0,30 |
| 4,57E-01 | 0,00 | 0,30 | 0,00 |
| 8,01E-01 | -0,22 | 0,10 | -0,10 |
| 5,78E-01 | 0,11 | 0,00 | -0,11 |
| 8,82E-01 | -0,22 | 0,10 | -0,16 |
| 8,01E-01 | -0,44 | -0,10 | -0,28 |
| 4,57E-01 | 0,00 | 0,30 | 0,00 |
| 8,01E-01 | 0,11 | 0,00 | 0,02 |
| 9,46E-01 | 0,00 | 0,00 | -0,10 |
| 4,57E-01 | 0,11 | 0,20 | 0,13 |
| 5,26E-01 | -0,22 | -0,30 | -0,14 |
| 4,57E-01 | 0,33 | 0,00 | -0,03 |
| 5,26E-01 | 0,00 | 0,10 | -0,10 |
| 9,00E-01 | 0,11 | 0,10 | 0,02 |
| 6,98E-01 | -0,11 | 0,10 | -0,19 |
| 7,25E-01 | 0,00 | -0,10 | 0,20 |
| 9,75E-01 | -0,22 | 0,20 | -0,49 |
| 8,01E-01 | 0,00 | -0,10 | 0,14 |
| 4,57E-01 | 0,00 | 0,30 | 0,00 |
| 8,01E-01 | 0,33 | 0,20 | -0,03 |
| 9,72E-01 | -0,11 | 0,10 | -0,19 |
| 8,16E-01 | -0,11 | 0,00 | 0,00 |
| 8,01E-01 | 0,11 | 0,10 | -0,09 |
| 9,00E-01 | -0,44 | 0,00 | -0,29 |
| 9,00E-01 | 0,00 | 0,00 | 0,00 |
| 5,78E-01 | -0,11 | 0,10 | -0,37 |
| 8,01E-01 | 0,11 | 0,00 | 0,01 |
| 9,02E-01 | 0,11 | -0,10 | 0,03 |
| 7,45E-01 | 0,00 | -0,10 | -0,08 |
| 4,57E-01 | -0,11 | -0,10 | 0,00 |
| 6,40E-01 | 0,00 | -0,20 | -0,10 |
| 9,00E-01 | 0,00 | 0,20 | -0,08 |
| 6,40E-01 | 0,22 | -0,10 | 0,22 |
| 6,31E-01 | 0,22 | 0,00 | -0,12 |
| 7,45E-01 | 0,11 | 0,00 | 0,14 |
| 6,31E-01 | -0,11 | -0,20 | 0,28 |

| | | | |
|---|---|---|---|
| 7,93E-01 | -0,44 | 0,30 | -0,50 |
| 8,01E-01 | 0,11 | 0,20 | 0,09 |
| 8,01E-01 | 0,11 | 0,00 | 0,13 |
| 8,01E-01 | -0,11 | -0,10 | 0,16 |
| 7,32E-01 | 0,22 | 0,30 | 0,29 |
| 6,75E-01 | 0,00 | -0,10 | 0,28 |
| 8,16E-01 | -0,11 | -0,10 | 0,08 |
| 7,77E-01 | -0,11 | 0,00 | -0,30 |
| 4,57E-01 | 0,00 | -0,20 | -0,19 |
| 5,26E-01 | -0,11 | 0,10 | 0,08 |
| 9,98E-01 | 0,00 | 0,00 | 0,01 |
| 8,01E-01 | 0,00 | 0,00 | 0,00 |
| 5,26E-01 | 0,00 | 0,00 | 0,00 |
| 8,01E-01 | 0,00 | 0,00 | 0,00 |
| 9,98E-01 | 0,00 | 0,00 | 0,00 |
| 6,34E-01 | 0,00 | 0,00 | 0,00 |
| 8,01E-01 | 0,00 | 0,00 | 0,00 |
| 4,57E-01 | 0,22 | -0,40 | 0,02 |
| 8,01E-01 | 0,22 | 0,00 | -0,04 |
| 8,01E-01 | 0,00 | 0,00 | -0,10 |
| 4,57E-01 | 0,00 | 0,00 | 0,00 |
| 8,01E-01 | 0,11 | 0,00 | -0,09 |
| 8,89E-01 | 0,11 | 0,00 | 0,10 |
| 6,70E-01 | 0,33 | 0,10 | 0,00 |
| 8,90E-01 | 0,22 | 0,20 | 0,08 |
| 8,29E-01 | 0,22 | 0,10 | 0,00 |
| 9,51E-01 | 0,00 | 0,20 | -0,04 |
| 9,51E-01 | 0,00 | 0,00 | -0,10 |
| 4,57E-01 | 0,00 | 0,00 | 0,00 |
| 8,16E-01 | 0,33 | 0,20 | 0,09 |
| 4,57E-01 | 0,00 | 0,00 | 0,37 |
| 5,26E-01 | 0,22 | -0,40 | 0,23 |
| 8,01E-01 | -0,22 | 0,10 | -0,09 |
| 8,01E-01 | 0,00 | 0,20 | -0,11 |
| 8,01E-01 | 0,00 | 0,00 | 0,00 |

| |
|---|
| **FMD / SD (T2)** |
| 0,00 |
| 0,00 |
| 0,00 |
| 0,00 |
| 0,00 |
| 0,00 |
| 0,00 |
| 0,11 |
| 0,59 |
| -0,20 |
| 0,01 |
| 0,27 |
| 0,00 |
| 0,00 |
| 0,00 |
| 0,00 |
| 0,22 |
| -0,30 |
| 0,50 |
| 0,01 |
| 0,00 |
| 0,01 |
| 0,39 |
| -0,57 |
| -0,14 |
| -0,28 |
| -0,39 |
| -0,38 |
| 0,70 |
| -0,12 |
| 0,33 |
| -0,26 |
| -0,20 |
| 0,29 |
| 0,38 |
| -0,07 |
| -0,10 |
| 0,09 |
| -0,02 |
| 0,00 |
| 0,22 |
| 0,30 |
| -0,17 |
| -0,36 |

| |
|---|
| 0,00 |
| -0,39 |
| 0,03 |
| -0,07 |
| 0,18 |
| -0,09 |
| -0,10 |
| 0,18 |
| -0,08 |
| -0,01 |
| -0,10 |
| -0,10 |
| 0,20 |
| -0,13 |
| 0,01 |
| -0,09 |
| -0,08 |
| 0,20 |
| 0,38 |
| -0,10 |
| 0,00 |
| 0,00 |
| 0,00 |
| 0,00 |
| 0,00 |
| -0,16 |
| 0,06 |
| -0,07 |
| -0,33 |
| 0,00 |
| -0,09 |
| -0,09 |
| -0,38 |
| 0,02 |
| -0,10 |
| 0,26 |
| 0,03 |
| 0,11 |
| -0,10 |
| -0,10 |
| -0,19 |
| -0,20 |
| -0,29 |
| -0,07 |
| -0,24 |
| 0,20 |

| |
|---|
| 0,00 |
| 0,00 |
| 0,00 |
| 0,03 |
| 0,03 |
| 0,13 |
| -0,04 |
| 0,27 |
| 0,01 |
| 0,12 |
| 0,03 |
| 0,30 |
| 0,22 |
| -0,22 |
| 0,17 |
| 0,07 |
| 0,30 |
| -0,09 |
| -0,10 |
| 0,22 |
| -0,22 |
| -0,37 |
| 0,00 |
| 0,01 |
| 0,02 |
| 0,10 |
| -0,07 |
| 0,04 |
| 0,30 |
| -0,17 |
| 0,02 |
| 0,11 |
| -0,10 |
| 0,16 |
| 0,00 |
| -0,16 |
| -0,10 |
| -0,18 |
| -0,18 |
| 0,01 |
| -0,30 |
| 0,12 |
| -0,10 |
| -0,34 |
| 0,03 |
| 0,19 |

| |
|---|
| 0,24 |
| 0,18 |
| 0,02 |
| 0,17 |
| 0,37 |
| 0,18 |
| 0,09 |
| -0,19 |
| -0,39 |
| 0,29 |
| 0,01 |
| 0,00 |
| 0,00 |
| 0,00 |
| 0,00 |
| 0,00 |
| 0,00 |
| -0,60 |
| -0,27 |
| -0,10 |
| 0,00 |
| -0,20 |
| -0,01 |
| -0,23 |
| 0,06 |
| -0,12 |
| 0,16 |
| -0,10 |
| 0,00 |
| -0,04 |
| 0,37 |
| -0,39 |
| 0,23 |
| 0,09 |
| 0,00 |

#### D_Phatway analysis

| | Mec | |
| --- | --- | --- |
| Pathway | SD (T1) | SD (T2) |
| PWY-6549: L-glutamine biosynthesis III | 240,07 | 1023,75 |
| P108-PWY: pyruvate fermentation to propanoate I | 81,61 | 470,17 |
| PWY-6507: 4-deoxy-L-threo-hex-4-enopyranuronate degradation | 177,63 | 3740,79 |
| VALSYN-PWY: L-valine biosynthesis | 6379,19 | 11951,10 |
| PWY-621: sucrose degradation III (sucrose invertase) | 18,20 | 3134,67 |
| PWY1ZNC-1: assimilatory sulfate reduction IV | 47,23 | 1519,21 |
| PWY-7315: dTDP-N-acetylthomosamine biosynthesis | 417,83 | 1256,93 |
| PWY-5676: acetyl-CoA fermentation to butanoate II | 53,32 | 326,23 |
| PWY-5675: nitrate reduction V (assimilatory) | 13,45 | 3625,11 |
| HOMOSER-METSYN-PWY: L-methionine biosynthesis I | 28,44 | 1552,26 |
| METSYN-PWY: superpathway of L-homoserine and L-methionine bios | 49,57 | 2490,80 |
| PWY-5347: superpathway of L-methionine biosynthesis (transsulfura | 56,53 | 2615,76 |
| PWY-5723: Rubisco shunt | 91,63 | 2642,59 |
| CITRULBIO-PWY: L-citrulline biosynthesis | 69,66 | 1746,79 |
| PWY-6305: superpathway of putrescine biosynthesis | 93,35 | 1680,56 |
| PWY-7242: D-fructuronate degradation | 203,33 | 3162,31 |
| PWY-6969: TCA cycle V (2-oxoglutarate synthase) | 199,92 | 2044,10 |
| RHAMCAT-PWY: L-rhamnose degradation I | 1112,03 | 3145,12 |
| PWY-6122: 5-aminoimidazole ribonucleotide biosynthesis II | 7525,27 | 12494,35 |
| PWY-6277: superpathway of 5-aminoimidazole ribonucleotide biosyr | 7525,27 | 12494,35 |
| PWY0-862: (5Z)-dodecenoate biosynthesis I | 107,10 | 6725,49 |
| PWY-6282: palmitoleate biosynthesis I (from (5Z)-dodec-5-enoate) | 107,02 | 6668,26 |
| PWY-7664: oleate biosynthesis IV (anaerobic) | 124,59 | 7254,07 |
| FASYN-ELONG-PWY: fatty acid elongation -- saturated | 141,97 | 7708,44 |
| PWY-5989: stearate biosynthesis II (bacteria and plants) | 105,80 | 5624,99 |
| SALVADEHYPOX-PWY: adenosine nucleotides degradation II | 231,21 | 6443,78 |
| FUCCAT-PWY: fucose degradation | 90,99 | 2334,25 |
| PWY-6121: 5-aminoimidazole ribonucleotide biosynthesis I | 5982,95 | 11876,23 |
| PWY-7198: pyrimidine deoxyribonucleotides de novo biosynthesis IV | 57,39 | 333,02 |
| PYRIDOXSYN-PWY: pyridoxal 5'-phosphate biosynthesis I | 16,16 | 3181,12 |
| PWY-7345: superpathway of anaerobic sucrose degradation | 98,63 | 6135,08 |
| P161-PWY: acetylene degradation (anaerobic) | 52,04 | 3259,58 |
| PWY-6519: 8-amino-7-oxononanoate biosynthesis I | 128,64 | 5894,20 |
| MET-SAM-PWY: superpathway of S-adenosyl-L-methionine biosynthe | 56,57 | 2602,71 |
| FERMENTATION-PWY: mixed acid fermentation | 269,38 | 3336,06 |
| TRPSYN-PWY: L-tryptophan biosynthesis | 1312,97 | 4270,86 |
| PWY-6629: superpathway of L-tryptophan biosynthesis | 2181,19 | 6036,65 |
| PWY-6608: guanosine nucleotides degradation III | 874,86 | 7084,53 |
| POLYAMSYN-PWY: superpathway of polyamine biosynthesis I | 28,80 | 1573,04 |
| BIOTIN-BIOSYNTHESIS-PWY: biotin biosynthesis I | 161,49 | 5946,06 |
| PWY4LZ-257: superpathway of fermentation (Chlamydomonas reinha | 62,18 | 2308,17 |
| PWY0-1298: superpathway of pyrimidine deoxyribonucleosides degra | 219,44 | 3263,41 |
| GALACT-GLUCUROCAT-PWY: superpathway of hexuronide and hexur | 146,32 | 2131,10 |
| GLUCUROCAT-PWY: superpathway of β-D-glucuronosides degra | 247,91 | 3372,22 |

| | | |
|---|---|---|
| P164-PWY: purine nucleobases degradation I (anaerobic) | 309,99 | 2687,56 |
| GALACTUROCAT-PWY: D-galacturonate degradation I | 413,25 | 3286,12 |
| PWY-5667: CDP-diacylglycerol biosynthesis I | 5233,26 | 10489,58 |
| PWY0-1319: CDP-diacylglycerol biosynthesis II | 5233,26 | 10489,58 |
| PWY-6163: chorismate biosynthesis from 3-dehydroquinate | 6912,72 | 10908,12 |
| PWY0-1477: ethanolamine utilization | 52,04 | 4327,38 |
| ARG+POLYAMINE-SYN: superpathway of arginine and polyamine bios | 57,07 | 2459,20 |
| P461-PWY: hexitol fermentation to lactate, formate, ethanol and ace | 135,52 | 3684,73 |
| SULFATE-CYS-PWY: superpathway of sulfate assimilation and cysteine | 190,95 | 4659,74 |
| PWY-6353: purine nucleotides degradation II (aerobic) | 467,70 | 6211,29 |
| NAGLIPASYN-PWY: lipid IVA biosynthesis (E. coli) | 701,87 | 5665,15 |
| PWY-8073: lipid IVA biosynthesis (P. putida) | 701,87 | 5665,15 |
| SO4ASSIM-PWY: assimilatory sulfate reduction I | 89,87 | 4017,12 |
| PWY-5345: superpathway of L-methionine biosynthesis (by sulfhydry | 237,85 | 5599,68 |
| P42-PWY: incomplete reductive TCA cycle | 60,25 | 852,85 |
| PWY-5497: purine nucleobases degradation II (anaerobic) | 231,53 | 2186,56 |
| COMPLETE-ARO-PWY: superpathway of aromatic amino acid biosynt | 4444,50 | 8657,85 |
| PWY-5136: fatty acid β-oxidation II (plant peroxisome) | 413,41 | 3960,32 |
| DAPLYSINESYN-PWY: L-lysine biosynthesis I | 448,25 | 3520,72 |
| PWY-702: L-methionine biosynthesis II | 320,91 | 1999,73 |
| ILEUSYN-PWY: L-isoleucine biosynthesis I (from threonine) | 3704,84 | 9866,42 |
| PWY-724: superpathway of L-lysine, L-threonine and L-methionine bi | 3512,38 | 7008,68 |
| PPGPPMET-PWY: ppGpp metabolism | 170,49 | 688,06 |
| PWY66-429: fatty acid biosynthesis initiation (mitochondria) | 5322,00 | 13339,71 |
| PWY-6609: adenine and adenosine salvage III | 9146,69 | 13191,58 |
| PWY-6834: spermidine biosynthesis III | 27,20 | 129,20 |
| P441-PWY: superpathway of N-acetylneuraminate degradation | 268,77 | 5291,82 |
| PWY-5505: L-glutamate and L-glutamine biosynthesis | 357,23 | 497,43 |
| P124-PWY: Bifidobacterium shunt | 1200,82 | 403,94 |
| PWY-8178: pentose phosphate pathway (non-oxidative branch) II | 1412,45 | 7490,54 |
| PWY-5941: glycogen degradation II | 2452,33 | 5781,71 |
| PWY-7400: L-arginine biosynthesis IV (archaebacteria) | 2373,04 | 5392,63 |
| PWY-I9: L-cysteine biosynthesis VI (from L-methionine) | 805,57 | 3680,01 |
| PWY-5384: sucrose degradation IV (sucrose phosphorylase) | 787,34 | 3335,56 |
| PWY-6731: starch degradation III | 1018,85 | 2887,19 |
| BRANCHED-CHAIN-AA-SYN-PWY: superpathway of branched chain ar | 3809,13 | 9649,95 |
| PWY-6606: guanosine nucleotides degradation II | 664,07 | 3618,57 |
| PWY0-162: superpathway of pyrimidine ribonucleotides de novo bios | 2672,51 | 6231,95 |
| ARO-PWY: chorismate biosynthesis I | 5975,20 | 9859,02 |
| GLUTORN-PWY: L-ornithine biosynthesis I | 1496,90 | 4309,76 |
| PWY-6906: chitin derivatives degradation | 225,57 | 2458,90 |
| PWY-5188: tetrapyrrole biosynthesis I (from glutamate) | 357,15 | 2851,62 |
| PWY-6387: UDP-N-acetylmuramoyl-pentapeptide biosynthesis I (mes | 9527,90 | 12136,98 |
| PWY-5913: partial TCA cycle (obligate autotrophs) | 335,84 | 2552,68 |
| NONOXIPENT-PWY: pentose phosphate pathway (non-oxidative bran | 923,01 | 6578,05 |
| PWY-241: C4 photosynthetic carbon assimilation cycle, NADP-ME typ | 939,64 | 4015,02 |

| | | |
|---|---|---|
| PWY-7111: pyruvate fermentation to isobutanol (engineered) | 2204,03 | 8772,66 |
| PWY-3001: superpathway of L-isoleucine biosynthesis I | 1686,75 | 6004,15 |
| THRESYN-PWY: superpathway of L-threonine biosynthesis | 2232,97 | 4996,77 |
| PWY-6147: 6-hydroxymethyl-dihydropterin diphosphate biosynthesis | 1630,35 | 4777,08 |
| ASPASN-PWY: superpathway of L-aspartate and L-asparagine biosynt | 2860,73 | 6551,53 |
| NONMEVIPP-PWY: methylerythritol phosphate pathway I | 6852,18 | 11469,30 |
| PWY-6385: peptidoglycan biosynthesis III (mycobacteria) | 8290,14 | 12442,50 |
| CALVIN-PWY: Calvin-Benson-Bassham cycle | 2665,37 | 10435,22 |
| PWY-7199: pyrimidine deoxyribonucleosides salvage | 3137,05 | 6049,79 |
| PEPTIDOGLYCANSYN-PWY: peptidoglycan biosynthesis I (meso-diami | 9777,11 | 12163,55 |
| PWY-7953: UDP-N-acetylmuramoyl-pentapeptide biosynthesis III (me | 7622,32 | 10859,13 |
| PWY-6700: queuosine biosynthesis I (de novo) | 8555,05 | 13626,14 |
| PWY-5103: L-isoleucine biosynthesis III | 3031,58 | 7977,56 |
| PWY-5154: L-arginine biosynthesis III (via N-acetyl-L-citrulline) | 1721,46 | 4110,29 |
| PWY-7117: C4 photosynthetic carbon assimilation cycle, PEPCK type | 969,17 | 3428,54 |
| SER-GLYSYN-PWY: superpathway of L-serine and glycine biosynthesis | 1881,09 | 6282,79 |
| PHOSLIPSYN-PWY: superpathway of phospholipid biosynthesis I (bact | 2921,11 | 7568,37 |
| ARGSYN-PWY: L-arginine biosynthesis I (via L-ornithine) | 2395,45 | 5829,63 |
| PWY-5973: cis-vaccenate biosynthesis | 6181,13 | 12731,70 |
| ARGSYNBSUB-PWY: L-arginine biosynthesis II (acetyl cycle) | 2066,24 | 5206,47 |
| PWY-7977: L-methionine biosynthesis IV | 7760,96 | 13671,91 |
| PWY-6630: superpathway of L-tyrosine biosynthesis | 73,63 | 1314,98 |
| PWY0-1296: purine ribonucleosides degradation | 4062,68 | 8155,02 |
| PWY-7357: thiamine phosphate formation from pyrithiamine and oxy | 1192,15 | 3804,10 |
| PWY0-1297: superpathway of purine deoxyribonucleosides degradat | 552,46 | 4366,00 |
| PWY-7323: superpathway of GDP-mannose-derived O-antigen buildi | 1209,15 | 4762,81 |
| PWY-1269: CMP-3-deoxy-D-manno-octulosonate biosynthesis | 1604,10 | 5256,14 |
| PWY-7328: superpathway of UDP-glucose-derived O-antigen building | 768,19 | 2189,91 |
| POLYISOPRENSYN-PWY: polyisoprenoid biosynthesis (E. coli) | 969,68 | 1916,99 |
| PWY-1042: glycolysis IV | 8124,09 | 12336,54 |
| PWY-4984: urea cycle | 48,29 | 355,51 |
| PWY0-1586: peptidoglycan maturation (meso-diaminopimelate conta | 2147,64 | 9478,29 |
| PWY-6386: UDP-N-acetylmuramoyl-pentapeptide biosynthesis II (lysi | 8209,15 | 11213,19 |
| PWY-6703: preQ0 biosynthesis | 5810,01 | 10575,16 |
| PWY-3841: folate transformations II (plants) | 12057,26 | 15119,89 |
| PWY-4041: γ-glutamyl cycle | 726,30 | 2982,61 |
| PWY-5659: GDP-mannose biosynthesis | 1417,04 | 4936,37 |
| PWY-7663: gondoate biosynthesis (anaerobic) | 5685,07 | 12009,15 |
| PWY-5097: L-lysine biosynthesis VI | 6980,88 | 10822,19 |
| GLYCOLYSIS: glycolysis I (from glucose 6-phosphate) | 6952,18 | 9273,61 |
| PENTOSE-P-PWY: pentose phosphate pathway | 1183,13 | 4987,19 |
| PWY-6901: superpathway of glucose and xylose degradation | 2000,02 | 5880,26 |
| PWY-7234: inosine-5'-phosphate biosynthesis III | 3671,00 | 7725,79 |
| TRNA-CHARGING-PWY: tRNA charging | 11663,13 | 14714,38 |
| PWY-6123: inosine-5'-phosphate biosynthesis I | 8259,10 | 11888,17 |
| LACTOSECAT-PWY: lactose and galactose degradation I | 3024,69 | 1197,71 |

| | | |
|---|---|---|
| PANTO-PWY: phosphopantothenate biosynthesis I | 8037,79 | 12925,36 |
| PWY-6126: superpathway of adenosine nucleotides de novo biosynth | 7449,56 | 10686,46 |
| PWY-7228: superpathway of guanosine nucleotides de novo biosynth | 6773,42 | 10140,51 |
| PWY-7229: superpathway of adenosine nucleotides de novo biosynth | 8739,60 | 11884,61 |
| PWY-5130: 2-oxobutanoate degradation I | 285,89 | 547,31 |
| PWY-6897: thiamine diphosphate salvage II | 1630,27 | 4998,95 |
| PWY4FS-7: phosphatidylglycerol biosynthesis I (plastidic) | 2114,68 | 5145,81 |
| PWY4FS-8: phosphatidylglycerol biosynthesis II (non-plastidic) | 2114,68 | 5145,81 |
| PWY-7356: thiamine diphosphate salvage IV (yeast) | 164,34 | 937,73 |
| PWY0-1261: anhydromuropeptides recycling I | 1150,53 | 4034,08 |
| PWY-7184: pyrimidine deoxyribonucleotides de novo biosynthesis I | 1440,70 | 4671,13 |
| PWY0-166: superpathway of pyrimidine deoxyribonucleotides de nov | 1563,73 | 4713,30 |
| PWY-6892: thiazole component of thiamine diphosphate biosynthesi | 2698,14 | 6222,32 |
| ANAEROFRUCAT-PWY: homolactic fermentation | 4181,32 | 7356,66 |
| PWY-7456: β-(1,4)-mannan degradation | 1041,36 | 2652,31 |
| PWY0-1241: ADP-L-glycero-β-D-manno-heptose biosynthesis | 20,59 | 477,49 |
| PWY-7761: NAD salvage pathway II (PNC IV cycle) | 1108,00 | 3693,89 |
| ARGININE-SYN4-PWY: L-ornithine biosynthesis II | 1491,43 | 2952,58 |
| PWY-7560: methylerythritol phosphate pathway II | 4988,80 | 8172,84 |
| PWY-6270: isoprene biosynthesis I | 5144,67 | 8174,71 |
| GLYCOGENSYNTH-PWY: glycogen biosynthesis I (from ADP-D-Glucose | 3421,73 | 6207,63 |
| PWY-5686: UMP biosynthesis I | 11514,24 | 14163,39 |
| PWY-7790: UMP biosynthesis II | 11514,24 | 14163,39 |
| PWY-7791: UMP biosynthesis III | 11514,24 | 14163,39 |
| PWY-6125: superpathway of guanosine nucleotides de novo biosynth | 6178,84 | 9344,85 |
| HSERMETANA-PWY: L-methionine biosynthesis III | 5118,26 | 6949,31 |
| COLANSYN-PWY: colanic acid building blocks biosynthesis | 2484,73 | 5686,20 |
| GLYCOLYSIS-E-D: superpathway of glycolysis and the Entner-Doudoro | 2895,15 | 5852,25 |
| GLYCOCAT-PWY: glycogen degradation I | 1995,44 | 3644,56 |
| PWY-5022: 4-aminobutanoate degradation V | 613,85 | 1803,27 |
| PWY-7238: sucrose biosynthesis II | 2353,65 | 6541,08 |
| PWY-6628: superpathway of L-phenylalanine biosynthesis | 214,33 | 1915,28 |
| GLUDEG-I-PWY: GABA shunt | 687,59 | 2074,57 |
| PWY-5695: inosine 5'-phosphate degradation | 9114,24 | 14092,54 |
| ANAGLYCOLYSIS-PWY: glycolysis III (from glucose) | 7921,70 | 11792,50 |
| PWY-5484: glycolysis II (from fructose 6-phosphate) | 6740,83 | 8370,41 |
| PWY-7221: guanosine ribonucleotides de novo biosynthesis | 9966,31 | 13575,46 |
| PWY-841: superpathway of purine nucleotides de novo biosynthesis | 6615,24 | 10207,22 |
| 1CMET2-PWY: folate transformations III (E. coli) | 8467,10 | 12005,60 |
| GLUCONEO-PWY: gluconeogenesis I | 6729,70 | 9158,79 |
| PWY-7220: adenosine deoxyribonucleotides de novo biosynthesis II | 5246,29 | 7809,40 |
| PWY-7222: guanosine deoxyribonucleotides de novo biosynthesis II | 5246,29 | 7809,40 |
| PANTOSYN-PWY: superpathway of coenzyme A biosynthesis I (bacter | 8132,05 | 12223,39 |
| THISYN-PWY: superpathway of thiamine diphosphate biosynthesis I | 3116,20 | 7300,08 |
| HISTSYN-PWY: L-histidine biosynthesis | 5009,93 | 7402,96 |
| PWY66-409: superpathway of purine nucleotide salvage | 1451,69 | 6159,29 |

| | | |
|---|---|---|
| COBALSYN-PWY: superpathway of adenosylcobalamin salvage from c | 876,44 | 2474,31 |
| PWY-2942: L-lysine biosynthesis III | 5864,00 | 8720,32 |
| PWY-6124: inosine-5'-phosphate biosynthesis II | 8046,83 | 11691,21 |
| PWY-7208: superpathway of pyrimidine nucleobases salvage | 5669,87 | 7646,28 |
| PWY-6151: S-adenosyl-L-methionine salvage I | 8553,35 | 9128,93 |
| DTDPRHAMSYN-PWY: dTDP-β-L-rhamnose biosynthesis | 12946,75 | 14779,23 |
| GLCMANNANAUT-PWY: superpathway of N-acetylglucosamine, N-ac | 2118,33 | 5562,81 |
| RIBOSYN2-PWY: flavin biosynthesis I (bacteria and plants) | 2629,06 | 5312,28 |
| P122-PWY: heterolactic fermentation | 47,51 | 789,24 |
| PWY-6607: guanosine nucleotides degradation I | 193,64 | 1688,07 |
| PYRIDNUCSYN-PWY: NAD de novo biosynthesis I (from aspartate) | 6798,39 | 9902,08 |
| COA-PWY-1: superpathway of coenzyme A biosynthesis III (mammals | 8470,10 | 11451,10 |
| PWY-6595: superpathway of guanosine nucleotides degradation (pla | 267,86 | 1567,66 |
| PWY-6545: pyrimidine deoxyribonucleotides de novo biosynthesis III | 1058,57 | 2510,59 |
| PWY-6612: superpathway of tetrahydrofolate biosynthesis | 395,57 | 1930,98 |
| PWY-6823: molybdopterin biosynthesis | 115,30 | 1234,35 |
| CENTFERM-PWY: pyruvate fermentation to butanoate | 120,96 | 549,10 |
| PWY-6317: D-galactose degradation I (Leloir pathway) | 4205,22 | 5617,37 |
| PWY-6590: superpathway of Clostridium acetobutylicum acidogenic f | 160,27 | 711,26 |
| UDPNAGSYN-PWY: UDP-N-acetyl-D-glucosamine biosynthesis I | 4935,10 | 6701,69 |
| COA-PWY: coenzyme A biosynthesis I (prokaryotic) | 9748,58 | 13151,56 |
| METH-ACETATE-PWY: methanogenesis from acetate | 94,39 | 169,88 |
| PWY-5121: superpathway of geranylgeranyl diphosphate biosynthesi | 1356,19 | 2465,40 |
| PWY-8004: Entner-Doudoroff pathway I | 3357,71 | 5179,76 |
| PWY-6936: seleno-amino acid biosynthesis (plants) | 2784,62 | 3697,80 |
| PWY-7197: pyrimidine deoxyribonucleotide phosphorylation | 4208,11 | 5614,99 |
| PWY-7851: coenzyme A biosynthesis II (eukaryotic) | 8517,39 | 11398,86 |
| PWY-6292: superpathway of L-cysteine biosynthesis (mammalian) | 689,34 | 634,70 |
| FOLSYN-PWY: superpathway of tetrahydrofolate biosynthesis and sal | 570,12 | 2699,74 |
| PWY-6859: all-trans-farnesol biosynthesis | 557,03 | 1035,47 |
| PWY-7237: myo-, chiro- and scyllo-inositol degradation | 1701,11 | 2709,23 |
| PWY-8187: L-arginine degradation XIII (reductive Stickland reaction) | 971,45 | 1575,32 |
| PWY-5030: L-histidine degradation III | 1061,50 | 2564,82 |
| GOLPDLCAT-PWY: superpathway of glycerol degradation to 1,3-propa | 44,06 | 65,22 |
| PWY-6527: stachyose degradation | 4740,89 | 5941,88 |
| PWY-6143: CMP-pseudaminate biosynthesis | 153,04 | 193,35 |
| PWY-7392: taxadiene biosynthesis (engineered) | 1064,65 | 638,53 |
| PWY-5910: superpathway of geranylgeranyldiphosphate biosynthesis | 617,76 | 297,04 |
| PWY-6293: superpathway of L-cysteine biosynthesis (fungi) | 409,73 | 885,73 |
| PWY-801: homocysteine and cysteine interconversion | 254,47 | 558,45 |
| THISYNARA-PWY: superpathway of thiamine diphosphate biosynthes | 1380,51 | 2344,52 |
| HISDEG-PWY: L-histidine degradation I | 5031,39 | 5947,74 |
| PWY-2941: L-lysine biosynthesis II | 2136,66 | 2634,35 |
| PWY-7316: dTDP-N-acetylviosamine biosynthesis | 120,38 | 141,30 |
| PWY-6902: chitin degradation II (Vibrio) | 1364,49 | 2361,04 |
| PWY-5100: pyruvate fermentation to acetate and lactate II | 4799,98 | 5072,41 |

| | | |
|---|---|---|
| OANTIGEN-PWY: O-antigen building blocks biosynthesis (E. coli) | 6097,72 | 5912,68 |
| PWY-5981: CDP-diacylglycerol biosynthesis III | 5126,47 | 3615,10 |
| PWY-7383: anaerobic energy metabolism (invertebrates, cytosol) | 493,70 | 493,40 |
| PWY66-399: gluconeogenesis III | 820,79 | 695,42 |
| P41-PWY: pyruvate fermentation to acetate and (S)-lactate I | 4799,98 | 5070,34 |
| PWY-922: mevalonate pathway I (eukaryotes and bacteria) | 750,08 | 326,88 |

| dian | | log2FC | | |
|---|---|---|---|---|
| FMD (T1) | FMD (T2) | T2 / T1 (SD) | T2 / T1 (FMD) | FMD / SD (T1) |
| 258,89 | 251,53 | 2,09 | -0,04 | 0,11 |
| 46,53 | 88,94 | 2,51 | 0,92 | -0,80 |
| 77,50 | 324,72 | 4,39 | 2,05 | -1,19 |
| 5126,61 | 6931,65 | 0,91 | 0,44 | -0,32 |
| 21,14 | 181,31 | 7,35 | 3,04 | 0,21 |
| 10,78 | 637,38 | 4,98 | 5,76 | -2,03 |
| 264,24 | 452,26 | 1,59 | 0,77 | -0,66 |
| 47,08 | 90,82 | 2,59 | 0,93 | -0,18 |
| 7,04 | 284,78 | 7,97 | 5,15 | -0,85 |
| 31,97 | 97,28 | 5,72 | 1,58 | 0,16 |
| 55,35 | 166,89 | 5,62 | 1,58 | 0,16 |
| 62,56 | 186,52 | 5,51 | 1,56 | 0,14 |
| 19,22 | 282,68 | 4,83 | 3,81 | -2,20 |
| 44,32 | 193,49 | 4,63 | 2,10 | -0,64 |
| 28,89 | 315,34 | 4,16 | 3,40 | -1,66 |
| 88,12 | 372,43 | 3,95 | 2,07 | -1,20 |
| 144,87 | 386,29 | 3,35 | 1,41 | -0,46 |
| 775,28 | 1105,22 | 1,50 | 0,51 | -0,52 |
| 7416,32 | 7158,01 | 0,73 | -0,05 | -0,02 |
| 7416,32 | 7158,01 | 0,73 | -0,05 | -0,02 |
| 57,79 | 577,56 | 5,96 | 3,30 | -0,88 |
| 57,76 | 575,99 | 5,95 | 3,30 | -0,88 |
| 67,24 | 660,44 | 5,85 | 3,28 | -0,88 |
| 76,65 | 740,13 | 5,75 | 3,25 | -0,88 |
| 57,15 | 544,55 | 5,72 | 3,23 | -0,88 |
| 121,08 | 683,91 | 4,79 | 2,49 | -0,93 |
| 35,66 | 306,26 | 4,67 | 3,07 | -1,33 |
| 5655,00 | 6187,75 | 0,99 | 0,13 | -0,08 |
| 113,53 | 86,62 | 2,52 | -0,39 | 0,97 |
| 12,00 | 703,44 | 7,54 | 5,76 | -0,40 |
| 112,38 | 840,58 | 5,94 | 2,89 | 0,19 |
| 21,14 | 1032,45 | 5,94 | 5,54 | -1,26 |
| 69,45 | 649,51 | 5,51 | 3,21 | -0,88 |
| 62,99 | 188,67 | 5,50 | 1,57 | 0,15 |
| 329,86 | 658,48 | 3,63 | 1,00 | 0,29 |
| 1294,73 | 1992,88 | 1,70 | 0,62 | -0,02 |
| 2001,13 | 3202,06 | 1,47 | 0,68 | -0,12 |
| 951,04 | 914,97 | 3,02 | -0,06 | 0,12 |
| 7,77 | 288,90 | 5,72 | 5,05 | -1,76 |
| 87,04 | 773,59 | 5,19 | 3,14 | -0,88 |
| 25,38 | 678,39 | 5,19 | 4,69 | -1,26 |
| 108,67 | 1163,73 | 3,89 | 3,41 | -1,01 |
| 71,74 | 271,20 | 3,86 | 1,90 | -1,02 |
| 113,31 | 462,41 | 3,76 | 2,02 | -1,12 |

| | | | | |
|---|---|---|---|---|
| 175,62 | 540,03 | 3,11 | 1,62 | -0,82 |
| 308,38 | 705,39 | 2,99 | 1,19 | -0,42 |
| 4699,15 | 4966,62 | 1,00 | 0,08 | -0,16 |
| 4699,15 | 4966,62 | 1,00 | 0,08 | -0,16 |
| 5957,94 | 7158,13 | 0,66 | 0,26 | -0,21 |
| 45,55 | 1551,71 | 6,35 | 5,06 | -0,19 |
| 15,46 | 503,76 | 5,40 | 4,94 | -1,82 |
| 124,37 | 594,92 | 4,75 | 2,25 | -0,12 |
| 52,06 | 1248,39 | 4,60 | 4,56 | -1,86 |
| 234,48 | 1080,43 | 3,73 | 2,20 | -0,99 |
| 897,11 | 1652,73 | 3,01 | 0,88 | 0,35 |
| 897,11 | 1652,73 | 3,01 | 0,88 | 0,35 |
| 23,58 | 950,74 | 5,47 | 5,28 | -1,89 |
| 64,01 | 1501,76 | 4,55 | 4,53 | -1,88 |
| 29,05 | 183,62 | 3,80 | 2,62 | -1,03 |
| 140,41 | 339,67 | 3,23 | 1,27 | -0,72 |
| 3530,56 | 6047,30 | 0,96 | 0,78 | -0,33 |
| 393,66 | 754,07 | 3,26 | 0,94 | -0,07 |
| 321,73 | 954,12 | 2,97 | 1,57 | -0,48 |
| 341,58 | 581,28 | 2,64 | 0,77 | 0,09 |
| 3395,20 | 4461,57 | 1,41 | 0,39 | -0,13 |
| 2785,44 | 3549,07 | 1,00 | 0,35 | -0,33 |
| 142,78 | 257,28 | 2,01 | 0,85 | -0,25 |
| 5338,61 | 5638,19 | 1,33 | 0,08 | 0,00 |
| 8264,23 | 9259,64 | 0,53 | 0,16 | -0,15 |
| 48,38 | 33,24 | 2,21 | -0,53 | 0,81 |
| 84,96 | 1924,20 | 4,29 | 4,49 | -1,65 |
| 371,27 | 329,25 | 0,48 | -0,17 | 0,06 |
| 899,10 | 1176,96 | -1,57 | 0,39 | -0,42 |
| 1081,40 | 2668,40 | 2,41 | 1,30 | -0,38 |
| 2571,45 | 3439,46 | 1,24 | 0,42 | 0,07 |
| 2505,39 | 3505,53 | 1,18 | 0,48 | 0,08 |
| 1015,17 | 1053,17 | 2,19 | 0,05 | 0,33 |
| 729,27 | 937,95 | 2,08 | 0,36 | -0,11 |
| 1312,67 | 932,04 | 1,50 | -0,49 | 0,37 |
| 3498,79 | 4472,24 | 1,34 | 0,35 | -0,12 |
| 753,89 | 847,11 | 2,44 | 0,17 | 0,18 |
| 1552,87 | 1646,04 | 1,22 | 0,08 | -0,78 |
| 5280,62 | 6493,64 | 0,72 | 0,30 | -0,18 |
| 1578,81 | 2913,37 | 1,52 | 0,88 | 0,08 |
| 430,09 | 564,42 | 3,44 | 0,39 | 0,93 |
| 320,96 | 1283,24 | 2,99 | 2,00 | -0,15 |
| 8242,22 | 7754,51 | 0,35 | -0,09 | -0,21 |
| 379,91 | 663,39 | 2,92 | 0,80 | 0,18 |
| 706,69 | 1982,92 | 2,83 | 1,49 | -0,38 |
| 1289,28 | 1257,80 | 2,09 | -0,04 | 0,46 |

| | | | | |
|---|---|---|---|---|
| 2403,47 | 3052,10 | 1,99 | 0,34 | 0,12 |
| 1529,49 | 1985,43 | 1,83 | 0,38 | -0,14 |
| 1881,33 | 1897,82 | 1,16 | 0,01 | -0,25 |
| 1544,13 | 1628,49 | 1,55 | 0,08 | -0,08 |
| 2239,40 | 2804,71 | 1,20 | 0,32 | -0,35 |
| 6001,90 | 7258,24 | 0,74 | 0,27 | -0,19 |
| 7530,55 | 6962,61 | 0,59 | -0,11 | -0,14 |
| 2240,90 | 4344,65 | 1,97 | 0,95 | -0,25 |
| 2922,87 | 4590,49 | 0,95 | 0,65 | -0,10 |
| 8368,95 | 7910,25 | 0,32 | -0,08 | -0,22 |
| 6959,36 | 6365,12 | 0,51 | -0,13 | -0,13 |
| 7130,24 | 8188,50 | 0,67 | 0,20 | -0,26 |
| 2765,17 | 3580,96 | 1,40 | 0,37 | -0,13 |
| 1599,92 | 1566,19 | 1,26 | -0,03 | -0,11 |
| 1181,58 | 1171,02 | 1,82 | -0,01 | 0,29 |
| 2118,60 | 2012,70 | 1,74 | -0,07 | 0,17 |
| 2840,46 | 3062,33 | 1,37 | 0,11 | -0,04 |
| 2719,29 | 4220,25 | 1,28 | 0,63 | 0,18 |
| 6384,73 | 5854,16 | 1,04 | -0,13 | 0,05 |
| 2204,45 | 4026,10 | 1,33 | 0,87 | 0,09 |
| 6047,74 | 6178,84 | 0,82 | 0,03 | -0,36 |
| 15,47 | 295,05 | 4,14 | 4,17 | -2,18 |
| 4139,94 | 5655,74 | 1,01 | 0,45 | 0,03 |
| 1810,95 | 1867,16 | 1,67 | 0,04 | 0,60 |
| 471,13 | 1929,46 | 2,98 | 2,03 | -0,23 |
| 1099,51 | 1405,18 | 1,98 | 0,35 | -0,14 |
| 1868,24 | 1647,31 | 1,71 | -0,18 | 0,22 |
| 871,68 | 1005,15 | 1,51 | 0,21 | 0,18 |
| 781,58 | 1000,44 | 0,98 | 0,36 | -0,31 |
| 6331,25 | 6973,00 | 0,60 | 0,14 | -0,36 |
| 36,99 | 119,18 | 2,85 | 1,66 | -0,38 |
| 2751,97 | 4543,57 | 2,14 | 0,72 | 0,36 |
| 7433,47 | 6764,00 | 0,45 | -0,14 | -0,14 |
| 5080,12 | 5391,36 | 0,86 | 0,09 | -0,19 |
| 9833,93 | 9645,48 | 0,33 | -0,03 | -0,29 |
| 852,38 | 1647,97 | 2,04 | 0,95 | 0,23 |
| 1578,55 | 1589,02 | 1,80 | 0,01 | 0,16 |
| 5910,27 | 5455,11 | 1,08 | -0,12 | 0,06 |
| 6473,09 | 6644,54 | 0,63 | 0,04 | -0,11 |
| 5851,17 | 5945,64 | 0,42 | 0,02 | -0,25 |
| 1052,54 | 1960,38 | 2,07 | 0,90 | -0,17 |
| 1812,19 | 2824,81 | 1,56 | 0,64 | -0,14 |
| 2968,44 | 2926,61 | 1,07 | -0,02 | -0,31 |
| 9958,02 | 9813,81 | 0,34 | -0,02 | -0,23 |
| 7203,01 | 6947,05 | 0,53 | -0,05 | -0,20 |
| 1786,78 | 2448,03 | -1,34 | 0,45 | -0,76 |

| | | | | |
|---|---|---|---|---|
| 6967,65 | 8033,01 | 0,69 | 0,21 | -0,21 |
| 5726,11 | 6512,61 | 0,52 | 0,19 | -0,38 |
| 5110,30 | 5903,16 | 0,58 | 0,21 | -0,41 |
| 7193,42 | 7568,13 | 0,44 | 0,07 | -0,28 |
| 267,55 | 216,76 | 0,93 | -0,30 | -0,10 |
| 2309,80 | 2422,64 | 1,62 | 0,07 | 0,50 |
| 2377,20 | 2371,51 | 1,28 | 0,00 | 0,17 |
| 2377,20 | 2371,51 | 1,28 | 0,00 | 0,17 |
| 184,55 | 413,94 | 2,51 | 1,16 | 0,17 |
| 1387,19 | 1207,30 | 1,81 | -0,20 | 0,27 |
| 1171,11 | 1663,75 | 1,70 | 0,51 | -0,30 |
| 1246,50 | 1761,54 | 1,59 | 0,50 | -0,33 |
| 2583,83 | 3487,74 | 1,21 | 0,43 | -0,06 |
| 4383,81 | 4030,80 | 0,81 | -0,12 | 0,07 |
| 1048,99 | 1035,54 | 1,35 | -0,02 | 0,01 |
| 19,24 | 157,32 | 4,47 | 2,97 | -0,09 |
| 1204,63 | 1737,93 | 1,74 | 0,53 | 0,12 |
| 1354,30 | 1063,75 | 0,98 | -0,35 | -0,14 |
| 4711,18 | 5032,92 | 0,71 | 0,10 | -0,08 |
| 4842,11 | 5056,47 | 0,67 | 0,06 | -0,09 |
| 3341,35 | 4872,24 | 0,86 | 0,54 | -0,03 |
| 10434,26 | 10129,65 | 0,30 | -0,04 | -0,14 |
| 10434,26 | 10129,65 | 0,30 | -0,04 | -0,14 |
| 10434,26 | 10129,65 | 0,30 | -0,04 | -0,14 |
| 4587,22 | 5390,94 | 0,60 | 0,23 | -0,43 |
| 4176,93 | 4208,59 | 0,44 | 0,01 | -0,29 |
| 2217,47 | 2364,29 | 1,19 | 0,09 | -0,16 |
| 2904,97 | 2564,69 | 1,02 | -0,18 | 0,00 |
| 1723,79 | 2122,89 | 0,87 | 0,30 | -0,21 |
| 318,91 | 666,02 | 1,55 | 1,06 | -0,94 |
| 2495,25 | 4718,16 | 1,47 | 0,92 | 0,08 |
| 313,99 | 648,44 | 3,15 | 1,04 | 0,55 |
| 600,34 | 775,56 | 1,59 | 0,37 | -0,20 |
| 8383,92 | 8161,33 | 0,63 | -0,04 | -0,12 |
| 6666,06 | 6528,71 | 0,57 | -0,03 | -0,25 |
| 5500,90 | 5769,81 | 0,31 | 0,07 | -0,29 |
| 8937,18 | 8490,43 | 0,45 | -0,07 | -0,16 |
| 5687,70 | 5499,29 | 0,63 | -0,05 | -0,22 |
| 6685,91 | 6837,14 | 0,50 | 0,03 | -0,34 |
| 5684,59 | 5592,83 | 0,44 | -0,02 | -0,24 |
| 3612,60 | 4482,43 | 0,57 | 0,31 | -0,54 |
| 3612,60 | 4482,43 | 0,57 | 0,31 | -0,54 |
| 7047,98 | 8041,17 | 0,59 | 0,19 | -0,21 |
| 3253,73 | 3581,98 | 1,23 | 0,14 | 0,06 |
| 4797,98 | 4801,11 | 0,56 | 0,00 | -0,06 |
| 1170,75 | 3555,24 | 2,08 | 1,60 | -0,31 |

| | | | | |
|---|---|---|---|---|
| 1060,49 | 1342,03 | 1,50 | 0,34 | 0,27 |
| 5649,34 | 5537,54 | 0,57 | -0,03 | -0,05 |
| 6986,03 | 6749,70 | 0,54 | -0,05 | -0,20 |
| 4083,69 | 4977,49 | 0,43 | 0,29 | -0,47 |
| 7552,69 | 7310,97 | 0,09 | -0,05 | -0,18 |
| 10547,66 | 10270,91 | 0,19 | -0,04 | -0,30 |
| 2736,01 | 2148,06 | 1,39 | -0,35 | 0,37 |
| 2900,11 | 3421,42 | 1,01 | 0,24 | 0,14 |
| 39,17 | 378,45 | 4,03 | 3,24 | -0,27 |
| 108,79 | 600,44 | 3,12 | 2,45 | -0,83 |
| 5649,21 | 5596,32 | 0,54 | -0,01 | -0,27 |
| 8217,10 | 8461,51 | 0,43 | 0,04 | -0,04 |
| 143,43 | 650,22 | 2,54 | 2,17 | -0,90 |
| 619,81 | 942,50 | 1,25 | 0,60 | -0,77 |
| 344,20 | 1220,48 | 2,28 | 1,82 | -0,20 |
| 182,76 | 856,65 | 3,41 | 2,22 | 0,66 |
| 124,48 | 265,99 | 2,17 | 1,09 | 0,04 |
| 3853,04 | 3949,67 | 0,42 | 0,04 | -0,13 |
| 164,17 | 344,26 | 2,14 | 1,06 | 0,03 |
| 3684,26 | 4893,18 | 0,44 | 0,41 | -0,42 |
| 8708,14 | 8877,35 | 0,43 | 0,03 | -0,16 |
| 123,90 | 118,66 | 0,84 | -0,06 | 0,39 |
| 1149,47 | 1474,14 | 0,86 | 0,36 | -0,24 |
| 3381,89 | 3149,14 | 0,63 | -0,10 | 0,01 |
| 2318,87 | 2893,08 | 0,41 | 0,32 | -0,26 |
| 2644,17 | 3515,11 | 0,42 | 0,41 | -0,67 |
| 7995,06 | 8174,64 | 0,42 | 0,03 | -0,09 |
| 675,64 | 973,81 | -0,12 | 0,53 | -0,03 |
| 500,97 | 1586,88 | 2,24 | 1,66 | -0,19 |
| 477,02 | 592,47 | 0,89 | 0,31 | -0,22 |
| 2066,18 | 2139,26 | 0,67 | 0,05 | 0,28 |
| 1019,70 | 1478,89 | 0,70 | 0,54 | 0,07 |
| 1038,25 | 755,09 | 1,27 | -0,46 | -0,03 |
| 38,43 | 216,94 | 0,56 | 2,47 | -0,19 |
| 3853,80 | 4241,11 | 0,33 | 0,14 | -0,30 |
| 195,66 | 195,72 | 0,34 | 0,00 | 0,35 |
| 788,09 | 1049,91 | -0,74 | 0,41 | -0,43 |
| 397,69 | 585,01 | -1,05 | 0,56 | -0,63 |
| 422,84 | 600,63 | 1,11 | 0,51 | 0,05 |
| 267,62 | 387,61 | 1,13 | 0,53 | 0,07 |
| 1902,87 | 1894,62 | 0,76 | -0,01 | 0,46 |
| 4014,34 | 4152,40 | 0,24 | 0,05 | -0,33 |
| 1459,36 | 2802,26 | 0,30 | 0,94 | -0,55 |
| 78,24 | 62,49 | 0,23 | -0,32 | -0,62 |
| 1428,94 | 1202,59 | 0,79 | -0,25 | 0,07 |
| 3609,98 | 4366,82 | 0,08 | 0,27 | -0,41 |

| | | | | |
|---|---|---|---|---|
| 4304,15 | 5551,52 | -0,04 | 0,37 | -0,50 |
| 3640,84 | 4198,42 | -0,50 | 0,21 | -0,49 |
| 421,53 | 535,39 | 0,00 | 0,34 | -0,23 |
| 522,44 | 801,20 | -0,24 | 0,62 | -0,65 |
| 3609,98 | 4366,82 | 0,08 | 0,27 | -0,41 |
| 512,33 | 650,14 | -1,20 | 0,34 | -0,55 |

| | P-value | | |
|---|---|---|---|
| **FMD / SD (T2)** | **T2 / T1 (SD)** | **T2 / T1 (FMD)** | **FMD / SD (T1)** |
| -2,02 | 3,91E-03 | 6,95E-01 | 6,61E-01 |
| -2,39 | 7,81E-03 | 3,22E-01 | 7,06E-01 |
| -3,52 | 3,91E-03 | 8,40E-02 | 7,89E-02 |
| -0,79 | 3,91E-03 | 3,75E-01 | 2,11E-01 |
| -4,10 | 3,91E-03 | 6,45E-02 | 4,81E-01 |
| -1,25 | 3,91E-03 | 1,78E-02 | 8,91E-02 |
| -1,47 | 7,81E-03 | 4,88E-02 | 5,49E-01 |
| -1,83 | 3,91E-03 | 6,25E-01 | 8,42E-01 |
| -3,67 | 3,91E-03 | 2,73E-02 | 3,40E-01 |
| -3,98 | 3,91E-03 | 8,40E-02 | 6,51E-01 |
| -3,89 | 3,91E-03 | 8,40E-02 | 6,51E-01 |
| -3,80 | 3,91E-03 | 8,40E-02 | 6,51E-01 |
| -3,22 | 3,91E-03 | 8,40E-02 | 7,87E-02 |
| -3,17 | 3,91E-03 | 2,73E-02 | 1,78E-01 |
| -2,41 | 3,91E-03 | 1,95E-02 | 7,89E-02 |
| -3,08 | 3,91E-03 | 1,05E-01 | 1,82E-01 |
| -2,40 | 3,91E-03 | 2,73E-02 | 5,49E-01 |
| -1,51 | 3,91E-03 | 4,92E-01 | 3,15E-01 |
| -0,80 | 7,81E-03 | 7,70E-01 | 3,56E-01 |
| -0,80 | 7,81E-03 | 7,70E-01 | 3,56E-01 |
| -3,54 | 3,91E-03 | 2,73E-02 | 2,20E-01 |
| -3,53 | 3,91E-03 | 2,73E-02 | 2,20E-01 |
| -3,46 | 3,91E-03 | 2,73E-02 | 2,20E-01 |
| -3,38 | 3,91E-03 | 2,73E-02 | 2,20E-01 |
| -3,37 | 3,91E-03 | 2,73E-02 | 2,20E-01 |
| -3,23 | 3,91E-03 | 4,88E-02 | 7,20E-01 |
| -2,93 | 3,91E-03 | 2,73E-02 | 3,56E-01 |
| -0,94 | 7,81E-03 | 3,22E-01 | 4,47E-01 |
| -1,93 | 1,43E-02 | 7,70E-01 | 4,62E-01 |
| -2,18 | 3,91E-03 | 3,71E-02 | 6,45E-01 |
| -2,87 | 3,91E-03 | 4,88E-02 | 4,81E-01 |
| -1,66 | 3,91E-03 | 1,95E-02 | 3,47E-01 |
| -3,18 | 3,91E-03 | 2,73E-02 | 2,20E-01 |
| -3,78 | 3,91E-03 | 8,40E-02 | 6,51E-01 |
| -2,34 | 3,91E-03 | 3,71E-02 | 6,04E-01 |
| -1,10 | 3,91E-03 | 6,45E-02 | 3,56E-01 |
| -0,91 | 3,91E-03 | 4,88E-02 | 4,47E-01 |
| -2,95 | 7,81E-03 | 4,32E-01 | 6,04E-01 |
| -2,44 | 3,91E-03 | 3,71E-02 | 6,38E-02 |
| -2,94 | 3,91E-03 | 2,73E-02 | 2,20E-01 |
| -1,77 | 3,91E-03 | 3,71E-02 | 3,47E-01 |
| -1,49 | 3,91E-03 | 1,37E-02 | 1,13E-01 |
| -2,97 | 3,91E-03 | 1,31E-01 | 2,05E-01 |
| -2,86 | 3,91E-03 | 1,05E-01 | 1,82E-01 |

| | | | |
|---|---|---|---|
| -2,31 | 3,91E-03 | 1,05E-01 | 3,07E-01 |
| -2,22 | 7,81E-03 | 4,88E-02 | 2,43E-01 |
| -1,08 | 1,17E-02 | 3,75E-01 | 4,00E-01 |
| -1,08 | 1,17E-02 | 3,75E-01 | 4,00E-01 |
| -0,61 | 1,95E-02 | 2,75E-01 | 2,43E-01 |
| -1,48 | 3,91E-03 | 1,37E-02 | 4,00E-01 |
| -2,29 | 3,91E-03 | 3,71E-02 | 6,38E-02 |
| -2,63 | 3,91E-03 | 8,40E-02 | 6,61E-01 |
| -1,90 | 3,91E-03 | 1,95E-02 | 9,36E-02 |
| -2,52 | 3,91E-03 | 4,88E-02 | 7,20E-01 |
| -1,78 | 3,91E-03 | 2,32E-01 | 3,56E-01 |
| -1,78 | 3,91E-03 | 2,32E-01 | 3,56E-01 |
| -2,08 | 3,91E-03 | 1,95E-02 | 9,36E-02 |
| -1,90 | 3,91E-03 | 1,95E-02 | 9,36E-02 |
| -2,21 | 3,91E-03 | 8,40E-02 | 1,11E-01 |
| -2,68 | 3,91E-03 | 1,93E-01 | 7,80E-01 |
| -0,52 | 3,91E-03 | 4,88E-02 | 4,47E-01 |
| -2,39 | 7,81E-03 | 8,40E-02 | 9,05E-01 |
| -1,88 | 7,81E-03 | 1,95E-02 | 6,61E-01 |
| -1,78 | 7,81E-03 | 1,60E-01 | 7,80E-01 |
| -1,14 | 7,81E-03 | 2,75E-01 | 6,61E-01 |
| -0,98 | 7,81E-03 | 5,57E-01 | 7,80E-01 |
| -1,42 | 1,17E-02 | 1,31E-01 | 7,20E-01 |
| -1,24 | 1,17E-02 | 4,92E-01 | 9,68E-01 |
| -0,51 | 1,17E-02 | 1,93E-01 | 2,43E-01 |
| -1,93 | 1,43E-02 | 6,95E-01 | 1,51E-01 |
| -1,46 | 3,91E-03 | 2,73E-02 | 3,07E-01 |
| -0,59 | 9,77E-02 | 9,22E-01 | 7,80E-01 |
| 1,54 | 1,64E-01 | 2,32E-01 | 6,04E-01 |
| -1,49 | 3,91E-03 | 2,73E-02 | 1,82E-01 |
| -0,75 | 3,91E-03 | 1,60E-01 | 7,20E-01 |
| -0,62 | 3,91E-03 | 1,31E-01 | 9,68E-01 |
| -1,80 | 7,81E-03 | 2,75E-01 | 1,00E+00 |
| -1,83 | 7,81E-03 | 5,57E-01 | 9,68E-01 |
| -1,63 | 7,81E-03 | 6,95E-01 | 1,33E-01 |
| -1,11 | 7,81E-03 | 3,22E-01 | 6,04E-01 |
| -2,09 | 1,95E-02 | 4,32E-01 | 5,49E-01 |
| -1,92 | 1,95E-02 | 5,57E-01 | 1,13E-01 |
| -0,60 | 1,95E-02 | 2,75E-01 | 4,97E-01 |
| -0,56 | 3,91E-02 | 8,40E-02 | 9,05E-01 |
| -2,12 | 3,91E-03 | 2,75E-01 | 1,53E-01 |
| -1,15 | 3,91E-03 | 1,37E-02 | 5,49E-01 |
| -0,65 | 5,47E-02 | 6,95E-01 | 4,97E-01 |
| -1,94 | 3,91E-03 | 1,95E-02 | 4,97E-01 |
| -1,73 | 3,91E-03 | 3,71E-02 | 1,13E-01 |
| -1,67 | 7,81E-03 | 4,32E-01 | 2,43E-01 |

| | | | |
|---|---|---|---|
| -1,52 | 7,81E-03 | 1,05E-01 | 9,05E-01 |
| -1,60 | 7,81E-03 | 1,93E-01 | 9,05E-01 |
| -1,40 | 1,17E-02 | 6,25E-01 | 4,47E-01 |
| -1,55 | 1,95E-02 | 6,95E-01 | 7,80E-01 |
| -1,22 | 1,95E-02 | 2,32E-01 | 1,56E-01 |
| -0,66 | 2,73E-02 | 2,75E-01 | 3,56E-01 |
| -0,84 | 2,73E-02 | 5,57E-01 | 4,97E-01 |
| -1,26 | 3,91E-03 | 2,73E-02 | 1,56E-01 |
| -0,40 | 3,91E-03 | 1,05E-01 | 5,49E-01 |
| -0,62 | 5,47E-02 | 7,70E-01 | 4,97E-01 |
| -0,77 | 5,47E-02 | 6,25E-01 | 5,49E-01 |
| -0,73 | 9,77E-02 | 6,25E-01 | 1,13E-01 |
| -1,16 | 7,81E-03 | 1,93E-01 | 4,97E-01 |
| -1,39 | 7,81E-03 | 4,92E-01 | 9,68E-01 |
| -1,55 | 1,17E-02 | 4,32E-01 | 4,47E-01 |
| -1,64 | 1,17E-02 | 6,25E-01 | 9,68E-01 |
| -1,31 | 1,95E-02 | 4,92E-01 | 9,68E-01 |
| -0,47 | 1,95E-02 | 1,05E-01 | 1,00E+00 |
| -1,12 | 1,95E-02 | 6,25E-01 | 6,61E-01 |
| -0,37 | 3,91E-02 | 6,45E-02 | 9,68E-01 |
| -1,15 | 3,91E-02 | 7,70E-01 | 1,13E-01 |
| -2,15 | 3,91E-03 | 3,71E-02 | 7,69E-01 |
| -0,53 | 3,91E-03 | 6,45E-02 | 7,20E-01 |
| -1,03 | 7,81E-03 | 3,22E-01 | 9,05E-01 |
| -1,18 | 1,17E-02 | 1,95E-02 | 2,78E-01 |
| -1,76 | 1,95E-02 | 2,75E-01 | 7,80E-01 |
| -1,67 | 1,95E-02 | 5,57E-01 | 7,80E-01 |
| -1,12 | 1,95E-02 | 6,95E-01 | 7,80E-01 |
| -0,94 | 2,73E-02 | 4,92E-01 | 9,05E-01 |
| -0,82 | 2,73E-02 | 4,32E-01 | 9,47E-02 |
| -1,57 | 3,91E-03 | 1,37E-02 | 1,78E-01 |
| -1,06 | 7,81E-03 | 6,45E-02 | 9,68E-01 |
| -0,73 | 7,42E-02 | 6,95E-01 | 6,04E-01 |
| -0,97 | 7,42E-02 | 6,25E-01 | 2,43E-01 |
| -0,65 | 3,59E-01 | 9,22E-01 | 2,78E-01 |
| -0,86 | 7,81E-03 | 4,88E-02 | 9,05E-01 |
| -1,63 | 1,95E-02 | 4,92E-01 | 6,61E-01 |
| -1,14 | 1,95E-02 | 6,25E-01 | 6,61E-01 |
| -0,70 | 2,73E-02 | 4,32E-01 | 2,43E-01 |
| -0,64 | 3,91E-02 | 6,25E-01 | 4,47E-01 |
| -1,35 | 7,81E-03 | 6,45E-02 | 3,15E-01 |
| -1,06 | 7,81E-03 | 1,05E-01 | 4,47E-01 |
| -1,40 | 5,47E-02 | 1,00E+00 | 4,97E-01 |
| -0,58 | 7,42E-02 | 8,46E-01 | 4,00E-01 |
| -0,77 | 9,77E-02 | 1,00E+00 | 4,00E-01 |
| 1,03 | 1,64E-01 | 6,25E-01 | 7,80E-01 |

| | | | |
|---|---|---|---|
| -0,69 | 1,64E-01 | 5,57E-01 | 3,56E-01 |
| -0,71 | 1,64E-01 | 6,95E-01 | 1,56E-01 |
| -0,78 | 1,64E-01 | 6,95E-01 | 1,56E-01 |
| -0,65 | 1,64E-01 | 5,57E-01 | 1,82E-01 |
| -1,33 | 2,50E-01 | 5,57E-01 | 1,00E+00 |
| -1,04 | 1,17E-02 | 3,22E-01 | 8,42E-01 |
| -1,12 | 1,17E-02 | 5,57E-01 | 9,68E-01 |
| -1,12 | 1,17E-02 | 5,57E-01 | 9,68E-01 |
| -1,18 | 1,95E-02 | 2,73E-02 | 9,68E-01 |
| -1,74 | 1,95E-02 | 5,57E-01 | 6,61E-01 |
| -1,49 | 1,95E-02 | 1,93E-01 | 9,68E-01 |
| -1,42 | 1,95E-02 | 1,93E-01 | 9,68E-01 |
| -0,83 | 1,95E-02 | 3,22E-01 | 9,68E-01 |
| -0,87 | 1,95E-02 | 6,25E-01 | 7,80E-01 |
| -1,36 | 3,91E-02 | 3,75E-01 | 7,20E-01 |
| -1,60 | 3,91E-03 | 9,77E-03 | 8,68E-01 |
| -1,09 | 1,95E-02 | 1,93E-01 | 1,00E+00 |
| -1,47 | 5,47E-02 | 8,46E-01 | 1,00E+00 |
| -0,70 | 5,47E-02 | 4,32E-01 | 6,04E-01 |
| -0,69 | 7,42E-02 | 5,57E-01 | 4,97E-01 |
| -0,35 | 9,77E-02 | 1,31E-01 | 9,68E-01 |
| -0,48 | 9,77E-02 | 6,95E-01 | 3,15E-01 |
| -0,48 | 9,77E-02 | 6,95E-01 | 3,15E-01 |
| -0,48 | 9,77E-02 | 6,95E-01 | 3,15E-01 |
| -0,79 | 1,64E-01 | 6,25E-01 | 1,13E-01 |
| -0,72 | 2,03E-01 | 9,22E-01 | 1,56E-01 |
| -1,27 | 1,95E-02 | 4,32E-01 | 4,97E-01 |
| -1,19 | 1,95E-02 | 9,22E-01 | 7,20E-01 |
| -0,78 | 1,95E-02 | 3,22E-01 | 9,05E-01 |
| -1,44 | 2,73E-02 | 1,31E-01 | 6,04E-01 |
| -0,47 | 3,91E-02 | 1,05E-01 | 8,42E-01 |
| -1,56 | 1,17E-02 | 3,71E-02 | 5,40E-01 |
| -1,42 | 1,95E-02 | 1,93E-01 | 8,42E-01 |
| -0,79 | 5,47E-02 | 6,95E-01 | 6,04E-01 |
| -0,85 | 7,42E-02 | 6,25E-01 | 4,00E-01 |
| -0,54 | 9,77E-02 | 6,95E-01 | 4,47E-01 |
| -0,68 | 9,77E-02 | 8,46E-01 | 2,78E-01 |
| -0,89 | 9,77E-02 | 6,95E-01 | 4,47E-01 |
| -0,81 | 1,29E-01 | 7,70E-01 | 2,43E-01 |
| -0,71 | 1,29E-01 | 7,70E-01 | 3,15E-01 |
| -0,80 | 1,64E-01 | 5,57E-01 | 1,33E-01 |
| -0,80 | 1,64E-01 | 5,57E-01 | 1,33E-01 |
| -0,60 | 2,50E-01 | 5,57E-01 | 3,15E-01 |
| -1,03 | 1,95E-02 | 4,32E-01 | 9,05E-01 |
| -0,62 | 3,91E-02 | 5,57E-01 | 5,49E-01 |
| -0,79 | 1,95E-02 | 3,71E-02 | 4,47E-01 |

| | | | |
|---|---|---|---|
| -0,88 | 1,95E-02 | 3,22E-01 | 3,15E-01 |
| -0,66 | 5,47E-02 | 4,92E-01 | 4,97E-01 |
| -0,79 | 2,03E-01 | 9,22E-01 | 4,00E-01 |
| -0,62 | 2,03E-01 | 6,25E-01 | 2,43E-01 |
| -0,32 | 2,50E-01 | 7,70E-01 | 7,80E-01 |
| -0,52 | 5,70E-01 | 1,00E+00 | 2,11E-01 |
| -1,37 | 1,95E-02 | 4,92E-01 | 7,20E-01 |
| -0,63 | 1,95E-02 | 1,60E-01 | 8,42E-01 |
| -1,06 | 3,91E-03 | 4,88E-02 | 3,84E-01 |
| -1,49 | 3,91E-03 | 4,88E-02 | 9,05E-01 |
| -0,82 | 1,29E-01 | 7,70E-01 | 4,00E-01 |
| -0,44 | 3,59E-01 | 6,95E-01 | 4,47E-01 |
| -1,27 | 3,91E-03 | 6,45E-02 | 9,68E-01 |
| -1,41 | 2,73E-02 | 1,93E-01 | 6,04E-01 |
| -0,66 | 7,81E-03 | 1,95E-02 | 1,00E+00 |
| -0,53 | 3,91E-03 | 1,37E-02 | 9,05E-01 |
| -1,04 | 5,47E-02 | 1,31E-01 | 9,05E-01 |
| -0,51 | 5,47E-02 | 4,32E-01 | 5,49E-01 |
| -1,04 | 5,47E-02 | 1,31E-01 | 9,05E-01 |
| -0,45 | 7,42E-02 | 4,92E-01 | 7,20E-01 |
| -0,57 | 3,59E-01 | 7,70E-01 | 4,97E-01 |
| -0,51 | 4,26E-01 | 7,70E-01 | 3,56E-01 |
| -0,74 | 5,47E-02 | 3,22E-01 | 8,42E-01 |
| -0,72 | 1,29E-01 | 6,95E-01 | 6,04E-01 |
| -0,35 | 1,64E-01 | 7,70E-01 | 8,42E-01 |
| -0,68 | 2,50E-01 | 4,32E-01 | 2,78E-01 |
| -0,48 | 3,59E-01 | 7,70E-01 | 6,04E-01 |
| 0,62 | 8,20E-01 | 2,75E-01 | 9,68E-01 |
| -0,77 | 7,81E-03 | 1,95E-02 | 1,00E+00 |
| -0,80 | 5,47E-02 | 4,32E-01 | 7,80E-01 |
| -0,34 | 7,81E-03 | 5,57E-01 | 6,04E-01 |
| -0,09 | 1,95E-02 | 1,05E-01 | 9,05E-01 |
| -1,76 | 2,50E-01 | 1,00E+00 | 7,20E-01 |
| 1,72 | 3,59E-01 | 1,95E-02 | 5,40E-01 |
| -0,49 | 2,03E-01 | 4,92E-01 | 5,49E-01 |
| 0,02 | 4,26E-01 | 8,46E-01 | 4,97E-01 |
| 0,72 | 4,96E-01 | 6,95E-01 | 1,00E+00 |
| 0,98 | 5,70E-01 | 1,00E+00 | 9,68E-01 |
| -0,56 | 7,42E-02 | 1,31E-01 | 4,97E-01 |
| -0,53 | 7,42E-02 | 1,05E-01 | 4,97E-01 |
| -0,31 | 1,64E-01 | 6,95E-01 | 6,04E-01 |
| -0,52 | 9,10E-01 | 8,46E-01 | 2,11E-01 |
| 0,09 | 1,00E+00 | 1,93E-01 | 6,04E-01 |
| -1,16 | 1,00E+00 | 6,95E-01 | 2,79E-02 |
| -0,97 | 1,29E-01 | 6,25E-01 | 6,04E-01 |
| -0,22 | 3,01E-01 | 6,25E-01 | 6,04E-01 |

| | | | |
|---|---|---|---|
| -0,09 | 5,70E-01 | 6,25E-01 | 6,61E-01 |
| 0,22 | 1,00E+00 | 7,70E-01 | 6,61E-01 |
| 0,12 | 6,52E-01 | 5,57E-01 | 9,68E-01 |
| 0,20 | 7,34E-01 | 3,22E-01 | 6,04E-01 |
| -0,22 | 8,20E-01 | 6,25E-01 | 6,04E-01 |
| 0,99 | 6,52E-01 | 9,22E-01 | 1,00E+00 |

| | Adjusted p-value | | |
|---|---|---|---|
| **FMD / SD (T2)** | **T2 / T1 (SD)** | **T2 / T1 (FMD)** | **FMD / SD (T1)** |
| 2,99E-03 | 1,34E-02 | 7,98E-01 | 9,66E-01 |
| 5,67E-03 | 1,88E-02 | 6,18E-01 | 9,85E-01 |
| 1,01E-02 | 1,34E-02 | 2,85E-01 | 9,66E-01 |
| 1,01E-02 | 1,34E-02 | 6,96E-01 | 9,66E-01 |
| 1,33E-02 | 1,34E-02 | 2,56E-01 | 9,66E-01 |
| 1,33E-02 | 1,34E-02 | 1,94E-01 | 9,66E-01 |
| 1,33E-02 | 1,88E-02 | 2,20E-01 | 9,66E-01 |
| 1,60E-02 | 1,34E-02 | 7,95E-01 | 1,00E+00 |
| 1,72E-02 | 1,34E-02 | 1,94E-01 | 9,66E-01 |
| 1,72E-02 | 1,34E-02 | 2,85E-01 | 9,66E-01 |
| 1,72E-02 | 1,34E-02 | 2,85E-01 | 9,66E-01 |
| 1,72E-02 | 1,34E-02 | 2,85E-01 | 9,66E-01 |
| 1,72E-02 | 1,34E-02 | 2,85E-01 | 9,66E-01 |
| 1,72E-02 | 1,34E-02 | 1,94E-01 | 9,66E-01 |
| 1,72E-02 | 1,34E-02 | 1,94E-01 | 9,66E-01 |
| 1,72E-02 | 1,34E-02 | 3,12E-01 | 9,66E-01 |
| 1,72E-02 | 1,34E-02 | 1,94E-01 | 9,66E-01 |
| 1,72E-02 | 1,34E-02 | 7,78E-01 | 9,66E-01 |
| 1,72E-02 | 1,88E-02 | 8,26E-01 | 9,66E-01 |
| 1,72E-02 | 1,88E-02 | 8,26E-01 | 9,66E-01 |
| 2,20E-02 | 1,34E-02 | 1,94E-01 | 9,66E-01 |
| 2,20E-02 | 1,34E-02 | 1,94E-01 | 9,66E-01 |
| 2,20E-02 | 1,34E-02 | 1,94E-01 | 9,66E-01 |
| 2,20E-02 | 1,34E-02 | 1,94E-01 | 9,66E-01 |
| 2,20E-02 | 1,34E-02 | 1,94E-01 | 9,66E-01 |
| 2,20E-02 | 1,34E-02 | 2,20E-01 | 9,85E-01 |
| 2,20E-02 | 1,34E-02 | 1,94E-01 | 9,66E-01 |
| 2,20E-02 | 1,88E-02 | 6,18E-01 | 9,66E-01 |
| 2,74E-02 | 2,98E-02 | 8,26E-01 | 9,66E-01 |
| 2,79E-02 | 1,34E-02 | 2,07E-01 | 9,66E-01 |
| 2,79E-02 | 1,34E-02 | 2,20E-01 | 9,66E-01 |
| 2,79E-02 | 1,34E-02 | 1,94E-01 | 9,66E-01 |
| 2,79E-02 | 1,34E-02 | 1,94E-01 | 9,66E-01 |
| 2,79E-02 | 1,34E-02 | 2,85E-01 | 9,66E-01 |
| 2,79E-02 | 1,34E-02 | 2,07E-01 | 9,66E-01 |
| 2,79E-02 | 1,34E-02 | 2,56E-01 | 9,66E-01 |
| 2,79E-02 | 1,34E-02 | 2,20E-01 | 9,66E-01 |
| 2,79E-02 | 1,88E-02 | 7,32E-01 | 9,66E-01 |
| 3,50E-02 | 1,34E-02 | 2,07E-01 | 9,66E-01 |
| 3,50E-02 | 1,34E-02 | 1,94E-01 | 9,66E-01 |
| 3,50E-02 | 1,34E-02 | 2,07E-01 | 9,66E-01 |
| 3,50E-02 | 1,34E-02 | 1,94E-01 | 9,66E-01 |
| 3,50E-02 | 1,34E-02 | 3,52E-01 | 9,66E-01 |
| 3,50E-02 | 1,34E-02 | 3,12E-01 | 9,66E-01 |

| | | | |
|---|---|---|---|
| 3,50E-02 | 1,34E-02 | 3,12E-01 | 9,66E-01 |
| 3,50E-02 | 1,88E-02 | 2,20E-01 | 9,66E-01 |
| 3,50E-02 | 2,49E-02 | 6,96E-01 | 9,66E-01 |
| 3,50E-02 | 2,49E-02 | 6,96E-01 | 9,66E-01 |
| 3,50E-02 | 3,20E-02 | 5,75E-01 | 9,66E-01 |
| 4,35E-02 | 1,34E-02 | 1,94E-01 | 9,66E-01 |
| 4,35E-02 | 1,34E-02 | 2,07E-01 | 9,66E-01 |
| 4,35E-02 | 1,34E-02 | 2,85E-01 | 9,66E-01 |
| 4,35E-02 | 1,34E-02 | 1,94E-01 | 9,66E-01 |
| 4,35E-02 | 1,34E-02 | 2,20E-01 | 9,85E-01 |
| 4,35E-02 | 1,34E-02 | 5,23E-01 | 9,66E-01 |
| 4,35E-02 | 1,34E-02 | 5,23E-01 | 9,66E-01 |
| 5,35E-02 | 1,34E-02 | 1,94E-01 | 9,66E-01 |
| 5,35E-02 | 1,34E-02 | 1,94E-01 | 9,66E-01 |
| 5,35E-02 | 1,34E-02 | 2,85E-01 | 9,66E-01 |
| 5,35E-02 | 1,34E-02 | 4,52E-01 | 9,92E-01 |
| 5,35E-02 | 1,34E-02 | 2,20E-01 | 9,66E-01 |
| 5,35E-02 | 1,88E-02 | 2,85E-01 | 1,00E+00 |
| 5,35E-02 | 1,88E-02 | 1,94E-01 | 9,66E-01 |
| 5,35E-02 | 1,88E-02 | 4,16E-01 | 9,92E-01 |
| 5,35E-02 | 1,88E-02 | 5,75E-01 | 9,66E-01 |
| 5,35E-02 | 1,88E-02 | 7,85E-01 | 9,92E-01 |
| 5,35E-02 | 2,49E-02 | 3,52E-01 | 9,85E-01 |
| 5,35E-02 | 2,49E-02 | 7,78E-01 | 1,00E+00 |
| 5,35E-02 | 2,49E-02 | 4,52E-01 | 9,66E-01 |
| 5,93E-02 | 2,98E-02 | 7,98E-01 | 9,66E-01 |
| 6,53E-02 | 1,34E-02 | 1,94E-01 | 9,66E-01 |
| 5,35E-02 | 1,22E-01 | 9,42E-01 | 9,92E-01 |
| 5,35E-02 | 1,88E-01 | 5,23E-01 | 9,66E-01 |
| 6,53E-02 | 1,34E-02 | 1,94E-01 | 9,66E-01 |
| 6,53E-02 | 1,34E-02 | 4,16E-01 | 9,85E-01 |
| 6,53E-02 | 1,34E-02 | 3,52E-01 | 1,00E+00 |
| 6,53E-02 | 1,88E-02 | 5,75E-01 | 1,00E+00 |
| 6,53E-02 | 1,88E-02 | 7,85E-01 | 1,00E+00 |
| 6,53E-02 | 1,88E-02 | 7,98E-01 | 9,66E-01 |
| 6,53E-02 | 1,88E-02 | 6,18E-01 | 9,66E-01 |
| 6,53E-02 | 3,20E-02 | 7,32E-01 | 9,66E-01 |
| 6,53E-02 | 3,20E-02 | 7,85E-01 | 9,66E-01 |
| 6,53E-02 | 3,20E-02 | 5,75E-01 | 9,66E-01 |
| 6,53E-02 | 5,82E-02 | 2,85E-01 | 1,00E+00 |
| 7,89E-02 | 1,34E-02 | 5,75E-01 | 9,66E-01 |
| 7,89E-02 | 1,34E-02 | 1,94E-01 | 9,66E-01 |
| 6,53E-02 | 7,53E-02 | 7,98E-01 | 9,66E-01 |
| 7,89E-02 | 1,34E-02 | 1,94E-01 | 9,66E-01 |
| 7,89E-02 | 1,34E-02 | 2,07E-01 | 9,66E-01 |
| 7,89E-02 | 1,88E-02 | 7,32E-01 | 9,66E-01 |

| | | | |
|---|---|---|---|
| 7,89E-02 | 1,88E-02 | 3,12E-01 | 1,00E+00 |
| 7,89E-02 | 1,88E-02 | 4,52E-01 | 1,00E+00 |
| 7,89E-02 | 2,49E-02 | 7,95E-01 | 9,66E-01 |
| 7,89E-02 | 3,20E-02 | 7,98E-01 | 9,92E-01 |
| 7,89E-02 | 3,20E-02 | 5,23E-01 | 9,66E-01 |
| 7,89E-02 | 4,27E-02 | 5,75E-01 | 9,66E-01 |
| 7,89E-02 | 4,27E-02 | 7,85E-01 | 9,66E-01 |
| 9,47E-02 | 1,34E-02 | 1,94E-01 | 9,66E-01 |
| 9,47E-02 | 1,34E-02 | 3,12E-01 | 9,66E-01 |
| 7,89E-02 | 7,53E-02 | 8,26E-01 | 9,66E-01 |
| 7,89E-02 | 7,53E-02 | 7,95E-01 | 9,66E-01 |
| 7,89E-02 | 1,22E-01 | 7,95E-01 | 9,66E-01 |
| 9,47E-02 | 1,88E-02 | 4,52E-01 | 9,66E-01 |
| 9,47E-02 | 1,88E-02 | 7,78E-01 | 1,00E+00 |
| 9,47E-02 | 2,49E-02 | 7,32E-01 | 9,66E-01 |
| 9,47E-02 | 2,49E-02 | 7,95E-01 | 1,00E+00 |
| 9,47E-02 | 3,20E-02 | 7,78E-01 | 1,00E+00 |
| 9,47E-02 | 3,20E-02 | 3,12E-01 | 1,00E+00 |
| 9,47E-02 | 3,20E-02 | 7,95E-01 | 9,66E-01 |
| 9,47E-02 | 5,82E-02 | 2,56E-01 | 1,00E+00 |
| 9,47E-02 | 5,82E-02 | 8,26E-01 | 9,66E-01 |
| 1,13E-01 | 1,34E-02 | 2,07E-01 | 9,92E-01 |
| 1,13E-01 | 1,34E-02 | 2,56E-01 | 9,85E-01 |
| 1,13E-01 | 1,88E-02 | 6,18E-01 | 1,00E+00 |
| 1,13E-01 | 2,49E-02 | 1,94E-01 | 9,66E-01 |
| 1,13E-01 | 3,20E-02 | 5,75E-01 | 9,92E-01 |
| 1,13E-01 | 3,20E-02 | 7,85E-01 | 9,92E-01 |
| 1,13E-01 | 3,20E-02 | 7,98E-01 | 9,92E-01 |
| 1,13E-01 | 4,27E-02 | 7,78E-01 | 1,00E+00 |
| 1,13E-01 | 4,27E-02 | 7,32E-01 | 9,66E-01 |
| 1,33E-01 | 1,34E-02 | 1,94E-01 | 9,66E-01 |
| 1,33E-01 | 1,88E-02 | 2,56E-01 | 1,00E+00 |
| 1,13E-01 | 9,76E-02 | 7,98E-01 | 9,66E-01 |
| 1,13E-01 | 9,76E-02 | 7,95E-01 | 9,66E-01 |
| 1,13E-01 | 3,84E-01 | 9,42E-01 | 9,66E-01 |
| 1,33E-01 | 1,88E-02 | 2,20E-01 | 1,00E+00 |
| 1,33E-01 | 3,20E-02 | 7,78E-01 | 9,66E-01 |
| 1,33E-01 | 3,20E-02 | 7,95E-01 | 9,66E-01 |
| 1,33E-01 | 4,27E-02 | 7,32E-01 | 9,66E-01 |
| 1,33E-01 | 5,82E-02 | 7,95E-01 | 9,66E-01 |
| 1,56E-01 | 1,88E-02 | 2,56E-01 | 9,66E-01 |
| 1,56E-01 | 1,88E-02 | 3,12E-01 | 9,66E-01 |
| 1,33E-01 | 7,53E-02 | 1,00E+00 | 9,66E-01 |
| 1,33E-01 | 9,76E-02 | 8,87E-01 | 9,66E-01 |
| 1,33E-01 | 1,22E-01 | 1,00E+00 | 9,66E-01 |
| 1,33E-01 | 1,88E-01 | 7,95E-01 | 9,92E-01 |

| | | | |
|---|---|---|---|
| 1,33E-01 | 1,88E-01 | 7,85E-01 | 9,66E-01 |
| 1,33E-01 | 1,88E-01 | 7,98E-01 | 9,66E-01 |
| 1,33E-01 | 1,88E-01 | 7,98E-01 | 9,66E-01 |
| 1,33E-01 | 1,88E-01 | 7,85E-01 | 9,66E-01 |
| 1,33E-01 | 2,75E-01 | 7,85E-01 | 1,00E+00 |
| 1,56E-01 | 2,49E-02 | 6,18E-01 | 1,00E+00 |
| 1,56E-01 | 2,49E-02 | 7,85E-01 | 1,00E+00 |
| 1,56E-01 | 2,49E-02 | 7,85E-01 | 1,00E+00 |
| 1,56E-01 | 3,20E-02 | 1,94E-01 | 1,00E+00 |
| 1,56E-01 | 3,20E-02 | 7,85E-01 | 9,66E-01 |
| 1,56E-01 | 3,20E-02 | 4,52E-01 | 1,00E+00 |
| 1,56E-01 | 3,20E-02 | 4,52E-01 | 1,00E+00 |
| 1,56E-01 | 3,20E-02 | 6,18E-01 | 1,00E+00 |
| 1,56E-01 | 3,20E-02 | 7,95E-01 | 9,92E-01 |
| 1,56E-01 | 5,82E-02 | 6,96E-01 | 9,85E-01 |
| 1,82E-01 | 1,34E-02 | 1,94E-01 | 1,00E+00 |
| 1,82E-01 | 3,20E-02 | 4,52E-01 | 1,00E+00 |
| 1,56E-01 | 7,53E-02 | 8,87E-01 | 1,00E+00 |
| 1,56E-01 | 7,53E-02 | 7,32E-01 | 9,66E-01 |
| 1,56E-01 | 9,76E-02 | 7,85E-01 | 9,66E-01 |
| 1,56E-01 | 1,22E-01 | 3,52E-01 | 1,00E+00 |
| 1,56E-01 | 1,22E-01 | 7,98E-01 | 9,66E-01 |
| 1,56E-01 | 1,22E-01 | 7,98E-01 | 9,66E-01 |
| 1,56E-01 | 1,22E-01 | 7,98E-01 | 9,66E-01 |
| 1,56E-01 | 1,88E-01 | 7,95E-01 | 9,66E-01 |
| 1,56E-01 | 2,29E-01 | 9,42E-01 | 9,66E-01 |
| 1,82E-01 | 3,20E-02 | 7,32E-01 | 9,66E-01 |
| 1,82E-01 | 3,20E-02 | 9,42E-01 | 9,85E-01 |
| 1,82E-01 | 3,20E-02 | 6,18E-01 | 1,00E+00 |
| 1,82E-01 | 4,27E-02 | 3,52E-01 | 9,66E-01 |
| 1,82E-01 | 5,82E-02 | 3,12E-01 | 1,00E+00 |
| 2,11E-01 | 2,49E-02 | 2,07E-01 | 9,66E-01 |
| 2,11E-01 | 3,20E-02 | 4,52E-01 | 1,00E+00 |
| 1,82E-01 | 7,53E-02 | 7,98E-01 | 9,66E-01 |
| 1,82E-01 | 9,76E-02 | 7,95E-01 | 9,66E-01 |
| 1,82E-01 | 1,22E-01 | 7,98E-01 | 9,66E-01 |
| 1,82E-01 | 1,22E-01 | 8,87E-01 | 9,66E-01 |
| 1,82E-01 | 1,22E-01 | 7,98E-01 | 9,66E-01 |
| 1,82E-01 | 1,56E-01 | 8,26E-01 | 9,66E-01 |
| 1,82E-01 | 1,56E-01 | 8,26E-01 | 9,66E-01 |
| 1,82E-01 | 1,88E-01 | 7,85E-01 | 9,66E-01 |
| 1,82E-01 | 1,88E-01 | 7,85E-01 | 9,66E-01 |
| 1,82E-01 | 2,75E-01 | 7,85E-01 | 9,66E-01 |
| 2,11E-01 | 3,20E-02 | 7,32E-01 | 1,00E+00 |
| 2,11E-01 | 5,82E-02 | 7,85E-01 | 9,66E-01 |
| 2,43E-01 | 3,20E-02 | 2,07E-01 | 9,66E-01 |

| | | | |
|---|---|---|---|
| 2,43E-01 | 3,20E-02 | 6,18E-01 | 9,66E-01 |
| 2,11E-01 | 7,53E-02 | 7,78E-01 | 9,66E-01 |
| 2,11E-01 | 2,29E-01 | 9,42E-01 | 9,66E-01 |
| 2,11E-01 | 2,29E-01 | 7,95E-01 | 9,66E-01 |
| 2,11E-01 | 2,75E-01 | 8,26E-01 | 9,92E-01 |
| 2,11E-01 | 5,93E-01 | 1,00E+00 | 9,66E-01 |
| 2,43E-01 | 3,20E-02 | 7,78E-01 | 9,85E-01 |
| 2,43E-01 | 3,20E-02 | 4,16E-01 | 1,00E+00 |
| 2,78E-01 | 1,34E-02 | 2,20E-01 | 9,66E-01 |
| 2,78E-01 | 1,34E-02 | 2,20E-01 | 1,00E+00 |
| 2,43E-01 | 1,56E-01 | 8,26E-01 | 9,66E-01 |
| 2,43E-01 | 3,84E-01 | 7,98E-01 | 9,66E-01 |
| 2,78E-01 | 1,34E-02 | 2,56E-01 | 1,00E+00 |
| 2,78E-01 | 4,27E-02 | 4,52E-01 | 9,66E-01 |
| 3,56E-01 | 1,88E-02 | 1,94E-01 | 1,00E+00 |
| 4,00E-01 | 1,34E-02 | 1,94E-01 | 1,00E+00 |
| 2,78E-01 | 7,53E-02 | 3,52E-01 | 1,00E+00 |
| 2,78E-01 | 7,53E-02 | 7,32E-01 | 9,66E-01 |
| 2,78E-01 | 7,53E-02 | 3,52E-01 | 1,00E+00 |
| 2,78E-01 | 9,76E-02 | 7,78E-01 | 9,85E-01 |
| 2,78E-01 | 3,84E-01 | 8,26E-01 | 9,66E-01 |
| 3,07E-01 | 4,51E-01 | 8,26E-01 | 9,66E-01 |
| 3,15E-01 | 7,53E-02 | 6,18E-01 | 1,00E+00 |
| 3,15E-01 | 1,56E-01 | 7,98E-01 | 9,66E-01 |
| 3,15E-01 | 1,88E-01 | 8,26E-01 | 1,00E+00 |
| 3,15E-01 | 2,75E-01 | 7,32E-01 | 9,66E-01 |
| 3,15E-01 | 3,84E-01 | 8,26E-01 | 9,66E-01 |
| 3,15E-01 | 8,35E-01 | 5,75E-01 | 1,00E+00 |
| 4,00E-01 | 1,88E-02 | 1,94E-01 | 1,00E+00 |
| 3,56E-01 | 7,53E-02 | 7,32E-01 | 9,92E-01 |
| 4,47E-01 | 1,88E-02 | 7,85E-01 | 9,66E-01 |
| 6,04E-01 | 3,20E-02 | 3,12E-01 | 1,00E+00 |
| 4,00E-01 | 2,75E-01 | 1,00E+00 | 9,85E-01 |
| 4,35E-02 | 3,84E-01 | 1,94E-01 | 9,66E-01 |
| 4,47E-01 | 2,29E-01 | 7,78E-01 | 9,66E-01 |
| 4,47E-01 | 4,51E-01 | 8,87E-01 | 9,66E-01 |
| 4,47E-01 | 5,23E-01 | 7,98E-01 | 1,00E+00 |
| 4,97E-01 | 5,93E-01 | 1,00E+00 | 1,00E+00 |
| 5,49E-01 | 9,76E-02 | 3,52E-01 | 9,66E-01 |
| 5,49E-01 | 9,76E-02 | 3,12E-01 | 9,66E-01 |
| 5,49E-01 | 1,88E-01 | 7,98E-01 | 9,66E-01 |
| 5,49E-01 | 9,22E-01 | 8,87E-01 | 9,66E-01 |
| 5,49E-01 | 1,00E+00 | 4,52E-01 | 9,66E-01 |
| 3,70E-02 | 1,00E+00 | 7,98E-01 | 9,66E-01 |
| 6,61E-01 | 1,56E-01 | 7,95E-01 | 9,66E-01 |
| 7,20E-01 | 3,29E-01 | 7,95E-01 | 9,66E-01 |

| | | | |
|---|---|---|---|
| 7,20E-01 | 5,93E-01 | 7,95E-01 | 9,66E-01 |
| 7,80E-01 | 1,00E+00 | 8,26E-01 | 9,66E-01 |
| 8,42E-01 | 6,72E-01 | 7,85E-01 | 1,00E+00 |
| 8,42E-01 | 7,54E-01 | 6,18E-01 | 9,66E-01 |
| 8,42E-01 | 8,35E-01 | 7,95E-01 | 9,66E-01 |
| 9,05E-01 | 6,72E-01 | 9,42E-01 | 1,00E+00 |

| FMD / SD (T2) |
|---|
| 1,67E-01 |
| 1,67E-01 |
| 1,67E-01 |
| 1,67E-01 |
| 1,67E-01 |
| 1,67E-01 |
| 1,67E-01 |
| 1,67E-01 |
| 1,67E-01 |
| 1,67E-01 |
| 1,67E-01 |
| 1,67E-01 |
| 1,67E-01 |
| 1,67E-01 |
| 1,67E-01 |
| 1,67E-01 |
| 1,67E-01 |
| 1,67E-01 |
| 1,67E-01 |
| 1,67E-01 |
| 1,67E-01 |
| 1,67E-01 |
| 1,67E-01 |
| 1,67E-01 |
| 1,67E-01 |
| 1,67E-01 |
| 1,67E-01 |
| 1,67E-01 |
| 1,67E-01 |
| 1,67E-01 |
| 1,67E-01 |
| 1,67E-01 |
| 1,67E-01 |
| 1,67E-01 |
| 1,67E-01 |
| 1,67E-01 |
| 1,67E-01 |
| 1,67E-01 |
| 1,67E-01 |
| 1,67E-01 |
| 1,67E-01 |
| 1,67E-01 |
| 1,67E-01 |
| 1,67E-01 |

| |
|---|
| 1,67E-01 |
| 1,67E-01 |
| 1,67E-01 |
| 1,67E-01 |
| 1,67E-01 |
| 1,71E-01 |
| 1,71E-01 |
| 1,71E-01 |
| 1,71E-01 |
| 1,71E-01 |
| 1,71E-01 |
| 1,71E-01 |
| 1,71E-01 |
| 1,71E-01 |
| 1,71E-01 |
| 1,71E-01 |
| 1,71E-01 |
| 1,71E-01 |
| 1,71E-01 |
| 1,71E-01 |
| 1,71E-01 |
| 1,71E-01 |
| 1,71E-01 |
| 1,71E-01 |
| 1,71E-01 |
| 1,76E-01 |
| 1,76E-01 |
| 1,71E-01 |
| 1,71E-01 |
| 1,76E-01 |
| 1,76E-01 |
| 1,76E-01 |
| 1,76E-01 |
| 1,76E-01 |
| 1,76E-01 |
| 1,76E-01 |
| 1,76E-01 |
| 1,76E-01 |
| 1,76E-01 |
| 1,76E-01 |
| 1,81E-01 |
| 1,81E-01 |
| 1,76E-01 |
| 1,81E-01 |
| 1,81E-01 |
| 1,81E-01 |

| |
|---|
| 1,81E-01 |
| 1,81E-01 |
| 1,81E-01 |
| 1,81E-01 |
| 1,81E-01 |
| 1,81E-01 |
| 1,81E-01 |
| 1,96E-01 |
| 1,96E-01 |
| 1,81E-01 |
| 1,81E-01 |
| 1,81E-01 |
| 1,96E-01 |
| 1,96E-01 |
| 1,96E-01 |
| 1,96E-01 |
| 1,96E-01 |
| 1,96E-01 |
| 1,96E-01 |
| 1,96E-01 |
| 1,96E-01 |
| 2,11E-01 |
| 2,11E-01 |
| 2,11E-01 |
| 2,11E-01 |
| 2,11E-01 |
| 2,11E-01 |
| 2,11E-01 |
| 2,11E-01 |
| 2,11E-01 |
| 2,21E-01 |
| 2,21E-01 |
| 2,11E-01 |
| 2,11E-01 |
| 2,11E-01 |
| 2,21E-01 |
| 2,21E-01 |
| 2,21E-01 |
| 2,21E-01 |
| 2,21E-01 |
| 2,26E-01 |
| 2,26E-01 |
| 2,21E-01 |
| 2,21E-01 |
| 2,21E-01 |
| 2,21E-01 |

| |
|---|
| 2,21E-01 |
| 2,21E-01 |
| 2,21E-01 |
| 2,21E-01 |
| 2,21E-01 |
| 2,26E-01 |
| 2,26E-01 |
| 2,26E-01 |
| 2,26E-01 |
| 2,26E-01 |
| 2,26E-01 |
| 2,26E-01 |
| 2,26E-01 |
| 2,26E-01 |
| 2,26E-01 |
| 2,38E-01 |
| 2,38E-01 |
| 2,26E-01 |
| 2,26E-01 |
| 2,26E-01 |
| 2,26E-01 |
| 2,26E-01 |
| 2,26E-01 |
| 2,26E-01 |
| 2,26E-01 |
| 2,26E-01 |
| 2,38E-01 |
| 2,38E-01 |
| 2,38E-01 |
| 2,38E-01 |
| 2,38E-01 |
| 2,63E-01 |
| 2,63E-01 |
| 2,38E-01 |
| 2,38E-01 |
| 2,38E-01 |
| 2,38E-01 |
| 2,38E-01 |
| 2,38E-01 |
| 2,38E-01 |
| 2,38E-01 |
| 2,38E-01 |
| 2,38E-01 |
| 2,63E-01 |
| 2,63E-01 |
| 2,93E-01 |

| |
|---|
| 2,93E-01 |
| 2,63E-01 |
| 2,63E-01 |
| 2,63E-01 |
| 2,63E-01 |
| 2,63E-01 |
| 2,93E-01 |
| 2,93E-01 |
| 3,20E-01 |
| 3,20E-01 |
| 2,93E-01 |
| 2,93E-01 |
| 3,20E-01 |
| 3,20E-01 |
| 3,93E-01 |
| 4,36E-01 |
| 3,20E-01 |
| 3,20E-01 |
| 3,20E-01 |
| 3,20E-01 |
| 3,20E-01 |
| 3,51E-01 |
| 3,51E-01 |
| 3,51E-01 |
| 3,51E-01 |
| 3,51E-01 |
| 3,51E-01 |
| 3,51E-01 |
| 4,36E-01 |
| 3,93E-01 |
| 4,78E-01 |
| 6,25E-01 |
| 4,36E-01 |
| 1,71E-01 |
| 4,78E-01 |
| 4,78E-01 |
| 4,78E-01 |
| 5,28E-01 |
| 5,71E-01 |
| 5,71E-01 |
| 5,71E-01 |
| 5,71E-01 |
| 5,71E-01 |
| 1,71E-01 |
| 6,81E-01 |
| 7,35E-01 |

| |
|---|
| 7,35E-01 |
| 7,94E-01 |
| 8,46E-01 |
| 8,46E-01 |
| 8,46E-01 |
| 9,05E-01 |
